# *SNCA* triplication disrupts proteostasis and extracellular architecture prior to neurodegeneration in human midbrain organoids

**DOI:** 10.1101/2025.05.28.656641

**Authors:** Elpida Statoulla, Maria Zafeiri, Kleanthi Chalkiadaki, Georgia Voudouri, Karmel S. Gkika, Christos Papazoglou, Thomas Durcan, Arkady Khoutorsky, Seyed Mehdi Jafarnejad, Sarah Maguire, Alexander Dityatev, Christos G. Gkogkas

## Abstract

Synucleinopathies, including Parkinson’s disease, are characterized by α-synuclein (SNCA) aggregation and progressive neurodegeneration, yet the early molecular events linking *SNCA* gene dosage to disrupted proteostasis remain poorly understood. Here, we used human midbrain organoids derived from induced pluripotent stem cells (iPSC) carrying an *SNCA* triplication (*SNCA* Trip) and the isogenic corrected line (*SNCA* Isog) to dissect early pathogenic mechanisms in a 3D human model of synucleinopathy. We combined immunohistochemistry, immunoblotting, tandem mass tag proteomics, bulk RNA sequencing, and ribosome profiling to systematically characterize molecular alterations in *SNCA* Trip organoids at day 50 (D50) and day 100 (D100) of maturation. *SNCA* Trip organoids exhibited increased α-synuclein accumulation, neuromelanin deposition, and activation of mTORC1 (p-rpS6), ERK1/2, AKT and p-eIF2α signalling pathways by D100. Proteomic and transcriptomic analyses revealed upregulation of cytoskeletal, synaptic, and axonal development pathways, alongside significant downregulation of extracellular matrix (ECM) components and upregulation of perineuronal net (PNN) genes. Ribosome profiling showed minimal global translational changes but uncovered selective translational buffering of neuronal and ECM-associated transcripts. Confocal imaging confirmed progressive disorganization of pericellular and interstitial ECM structures around neurons in *SNCA* Trip organoids. Our findings demonstrate that *SNCA* triplication induces early proteostatic disruption and extracellular matrix remodelling prior to neurodegeneration and suggest that altered gene expression and ECM homeostasis may contribute to disease initiation and progression. Targeting these early aberrant mechanisms may offer new therapeutic opportunities for synucleinopathies, such as Parkinson’s Disease.

## Introduction

Synucleinopathies, including Parkinson’s disease (PD), dementia with Lewy bodies, and multiple system atrophy, are characterized by the pathological accumulation of α-synuclein protein, encoded by the *SNCA* gene, which disrupts cellular homeostasis and drives neurodegeneration. *SNCA* multiplications, such as triplications and duplications, exemplify this process, with triplication linked to early-onset parkinsonism (∼35 years), cognitive decline, and severe striatal dopaminergic deficits, while duplication mirrors idiopathic PD with later onset (∼50 years) and milder progression^1–3^. Central to these disorders is SNCA protein aggregation, which perturbs proteostasis, the balance of protein synthesis, folding, and degradation, leading to cellular stress and neuronal loss, particularly in dopaminergic systems^4,5^. Studies across animal and human models reveal that SNCA overexpression exacerbates this imbalance, activating stress response pathways like the unfolded protein response (UPR) and integrated stress response (ISR), which attempt to mitigate proteotoxic damage but may ultimately amplify pathology^6,7^.

Invertebrate models such as Drosophila and Caenorhabditis elegans have proven invaluable for dissecting the cellular toxicity of α-synuclein, successfully recapitulating protein aggregation, synaptic deficits, and neuronal dysfunction while enabling high-throughput modifier screens^8,9^. Mammalian systems, including transgenic mice, rats, and viral vector-based models, have advanced our understanding by demonstrating how SNCA overexpression triggers proteostatic collapse, mitochondrial dysfunction, and progressive dopaminergic cell death^10,11^. These rodent models faithfully reproduce key disease hallmarks, including progressive aggregation, phosphorylation at S129, motor impairments, synaptic dysfunction, and compromised autophagy^10,12,13^. Non-human primate studies have validated the cross-species relevance of these pathological mechanisms, though high costs and ethical considerations limit their widespread application.

To complement these animal models, human iPSC-derived systems offer genetic precision for modelling early pathogenic events in patient-specific backgrounds^4^. These human models bridge genetic fidelity with multicellular interactions absent in simpler systems, providing critical insights into PD mechanisms^4,14–16^. Human iPSC-derived models of *SNCA* triplication recapitulate proteostatic collapse and α-synuclein aggregation in patient-relevant systems^16^. More recently, 3D midbrain organoids have been developed to better mimic the spatial architecture and cell diversity of the human midbrain^14,15^. Across several studies, iPSC-derived dopaminergic neurons and 3D midbrain/cortical organoids with *SNCA* triplication exhibit 2- to 4-fold increases in α-synuclein mRNA and protein, yielding oligomeric and phosphorylated aggregates^14,17,18^. These models display disease hallmarks: mitochondrial dysfunction, synaptic decline, and dopaminergic (tyrosine hydroxylase; TH) neuron loss, alongside Endoplasmic Reticulum (ER) stress and UPR activation, underscoring proteostatic failure^7,15,19^. Patient brain tissue and cerebrospinal fluid (CSF) corroborate these findings, showing 2- to 3-fold α-synuclein elevations with aggregation linked to nuclear aging and lipid metabolism alterations^5,20,21^.

Translational control is a master regulator of proteostasis. Translational regulators like eEF2K, tied to the ISR, show increased activity in PD tissue, amplifying toxicity, while microRNAs (miR-7, miR-153) and iron-dependent mechanisms reduce α-synuclein levels^22–24^. Translational control studies further show that pathologic α-synuclein activates mammalian target of rapamycin complex 1 (mTORC1) via Tuberous sclerosis complex 2 (TSC2) binding, enhancing protein synthesis and aggregation, while its inhibition rescues neurodegeneration^6^. Although direct evidence of UPR/ISR activation is less frequent in these models, the heightened protein load suggests a stress response, aligning with cellular efforts to restore homeostasis.

Herein, we studied a human midbrain organoid model derived from human iPSC (hiPSC) harbouring *SNCA* Triplication (*SNCA* Trip) and the isogenic CRISPR/Cas9 corrected line (*SNCA* Isog)^14^ to investigate if altered proteostasis could be a driver of pathology in a 3D model of synucleinopathy. *SNCA* Trip organoids displayed increased SNCA and p-SNCA (S129) expression and prominent neuromelanin depositions both at 50 and 100 days (D) in culture. We investigated signalling related to protein synthesis, a key player in the proteostatic pathway and detected elevated activity of mTORC1 (phospho-rpS6 S240/244), ERK (phospho-T202/Y204), AKT (phospho-S473) and eIF2α (phospho-S51) at D100, but only p-rpS6 (S240/244) at D50. To further assess the effect of dysregulated signalling pathways on proteostasis we performed a multiomic analysis. Bulk proteomics of D100 midbrain organoids showed increased expression in *SNCA* Trip of axonogenesis, axonal development, neurotransmitter, synaptic vesicle cycle, synapse organization and membrane transporter-related proteins, together with decreased expression of actin-binding, cytoskeletal motor activity and extracellular matrix proteins, compared with *SNCA* Isog. Bulk RNA-seq corroborated these findings, showing significant downregulation in the expression of ECM-related mRNAs and upregulation of mRNAs coding for proteins abundant in Perineuronal Nets (PNNs). Strikingly bulk translatomics of actively translated mRNAs measured with Ribosome Profiling revealed modest changes in overall translation in *SNCA* Trip compared with *SNCA* Isog, however we observed evidence of translational buffering in mRNAs coding for ECM, PNN and neuronal, synaptic and axonal proteins. Using confocal imaging and immunofluorescence we confirmed increased staining for pericellular (PNN) and interstitial ECM in microtubule-associated protein 2 (MAP2)-positive cells in *SNCA* Trip at D100 but not D50, compared with *SNCA* Isog organoids. Interestingly, TH^+^ cells displayed significantly increased PNN and interstitial ECM staining earlier, at D50, with a strong trend for increase at D100, but not reaching statistical significance. Together these data reveal progressive phenotypes downstream of SNCA triplication, impacting synapses, axonal transport and ECM structure, concomitant with disrupted proteostasis.

## Materials and methods

### iPSC lines

iPSC lines *SNCA* Triplication (AST23 P35+P7 MT) and *SNCA* Isogenic (AST23-2KO-8B P11MT) were obtained from the Biobank of the Montreal Neurological Hospital and Institute (C-BIG). Sanger sequencing and pluripotency assessments were performed for each batch, using PCR for genetic validation and quantitative PCR (qPCR) and immunofluorescence (IF) to evaluate the expression of pluripotency markers OCT3/4 and NANOG. Karyotype analysis was conducted approximately every 10 passages using the Comparative Genome Hybridisation (CGH) array method. All experiments involving iPSCs were approved by the FORTH Ethics and Deontology Committee.

### Cell culture and midbrain organoids generation

iPSC lines were maintained at 37°C with 5% CO₂ in mTeSR Plus (STEMCELL Technologies, #05825) on Matrigel®-coated plates (Corning, #354277) and passaged using 0.5 mM EDTA (Thermo Fisher Scientific, #15575020). For generation of human midbrain organoids (hMOs), iPSCs were used only after at least two passages following thawing and were not passaged more than ten times. hMOs were generated as previously described^14^. Briefly, 10,000 cells from a single-cell suspension of iPSCs were seeded into each well of a Nunclon™ Sphera™ 96-well, U-bottom, Sphera-treated microplate (Thermo Scientific™, #174925) containing neuronal induction medium [DMEM/F12:Neurobasal (1:1) (Gibco, #21103049), supplemented with 1:100 N2 (Thermo Fisher Scientific, #17502048), 1:50 B27 without vitamin A (Thermo Fisher Scientific, #12587010), 1% GlutaMAX (Thermo Fisher Scientific, #35050061), 1% MEM Non-Essential Amino Acids (Thermo Fisher Scientific, #11140-050), and 0.1% β-mercaptoethanol (Gibco, #21985-023), along with 1 μg/mL heparin (Sigma-Aldrich, #H3149-10KU), 10 μM SB431542 (Selleck Chemicals, #S1067), 200 ng/mL Noggin (ImmunoTools, #11344834-rh), 0.8 μM CHIR99021 (Cayman, #13122), and 10 μM ROCK inhibitor Y27632 (Cayman, #129830-38-2)]. Plates were centrifuged for 3 min at 1200 rpm to aggregate the cells. After 48 h, embryoid bodies (EBs) were formed, and the medium was replaced with fresh neuronal induction medium lacking ROCK inhibitor. Two days later, the medium was replaced with midbrain patterning medium [neuronal induction medium supplemented with 200 ng/mL SHH-C25II (ImmunoTools, #11344074) and 100 ng/mL FGF8 (ImmunoTools, #11344834)] to promote midbrain identity. After 3 additional days, EBs were embedded in reduced growth factor Matrigel®GFR (Corning, #354230) and incubated for 24 h in tissue induction medium [neurobasal medium supplemented with 1:100 N2, 1:50 B27 without vitamin A, 1% GlutaMAX, 1% MEM Non-Essential Amino Acids, 0.1% β-mercaptoethanol, 2.5 μg/mL Insulin (Capricorn Scientific, #INS-K), 200 ng/mL laminin (Sigma-Aldrich, #L2020), 100 ng/mL SHH-C25II, and 100 ng/mL FGF8]. The following day, embedded hMOs were transferred to six-well ultra-low attachment plates containing final differentiation medium [neurobasal medium supplemented with 1:100 N2, 1:50 B27 without vitamin A, 1% GlutaMAX, 1% MEM Non-Essential Amino Acids, 0.1% β-mercaptoethanol, 10 ng/mL BDNF (ImmunoTools, #11343375), 10 ng/mL GDNF (ImmunoTools, #11343793), 100 μM ascorbic acid (Sigma-Aldrich, #A8960-5g), and 125 μM db-cAMP (Santa Cruz Biotechnology, #sc-201567A)] and cultured on an orbital shaker (Heathrow Scientific, #5003396). Full media exchanges were performed 3 times per week. For quality control, mycoplasma testing was performed monthly on both iPSCs and hMOs to confirm the absence of contamination.

### Brightfield Image analysis

Brightfield images were acquired using an EVOS™ XL Core microscope (Invitrogen). Quantification of projected surface area and diameter of the organoids was performed using OrgM, a Jython-based macro for automated measurement of organoid size (diameter and area) and shape (roundness and circularity) from brightfield images. OrgM was developed by Eddie Cai and Rhalena A. Thomas and is openly available from the Montreal Neurological Institute (MNI), Canada.

### Immunofluorescence and confocal imaging

D50 and D100 midbrain organoids were fixed overnight (o/n) in 4% (w/v) paraformaldehyde (PFA) in PBS at 4°C, cryoprotected in 20% (w/v) sucrose in PBS o/n or until they sank, at 4°C, and embedded in optimal cutting temperature (O.C.T.) compound (Shakura Finetek USA Inc., #4583). O.C.T.-embedded organoids were frozen using a dry ice/ethanol bath and stored at -80°C until further processing. Organoids were sectioned at 18 μm thickness using a cryostat (Leica Cryostat), blocked in 10% (v/v) normal goat serum (NGS) solution containing 0.3% Triton X-100 in PBS, and incubated o/n with primary antibodies at 4°C. All primary antibodies were detected using Alexa Fluor-conjugated secondary antibodies (Invitrogen), incubated for 90 min at RT. A detailed list of antibodies can be found in Sup. Table 4. DRAQ5 (Abcam, #ab108410) and DAPI (Abcam, #ab228549) were used for nuclear staining. Imaging was performed using a LEICA SP5 and a NIKON A1R HD confocal microscope, with ×40 oil immersion, ×20 air, and ×100 oil immersion objectives. For all immunofluorescence experiments, a minimum of 3 slices per organoid were imaged and analysed.

### Immunoblotting

D50 or D100 midbrain organoids were transferred from culture plates, briefly washed in ice-cold DPBS (PAN-Biotech, #P04-36500), and subsequently homogenized in RIPA buffer [50 mM Tris-HCl pH 8, 150 mM NaCl, 1% NP-40, 0.5% sodium deoxycholate (SDC), 1% sodium dodecyl sulphate (SDS)] supplemented with protease (Sigma-Aldrich, #P8340-1ML) and phosphatase inhibitors (Sigma-Aldrich, #P0044-1ML), using a motorised pestle homogenizer and sonicator. For each biological replicate, 3–5 organoids were pooled and lysed together, sonicated twice at 13% amplitude for 10 s (Digital Sonifier 250, Marshall Scientific), incubated on ice for 15 min with occasional vortexing, and finally centrifuged for 20 min at 16,000 × *g* at 4°C. Protein concentration of each sample was determined using the BCA protein assay (Pierce™ BCA Protein Assay, ThermoFisher Scientific). A total of 50 μg protein per lane was prepared in SDS sample buffer (50 mM Tris pH 6.8, 100 mM DTT, 2% SDS, 10% glycerol, 0.1% bromophenol blue), heated at 95°C for 5 min, and resolved on polyacrylamide gels. Proteins were transferred to 0.2 μm nitrocellulose membranes (Bio-Rad), blocked for 1 h at RT in 5% bovine serum albumin (BSA) in TBS-T, and incubated o/n with primary antibodies at 4°C. Fluorescent secondary antibodies were used for all immunoblotting experiments. A detailed list of antibodies can be found in Sup. Table 4. Blots were imaged using an Azure imaging system (Azure Biosystems) and quantified using Image Studio Software (LI-COR Biosciences) by measuring the intensity of each protein band. HSC70 or β-actin was used as a loading control. Data are shown as arbitrary units (AU) as a proxy for protein expression, normalized to the isogenic control group. For protein phosphorylation, phospho-protein values were divided by the corresponding normalized total protein values after subtraction of immunoblot background intensity (Image Studio Software, LI-COR Biosciences). For each experiment, values from SNCA triplication organoids were normalized to the mean of the SNCA isogenic group.

### TMT-proteomics and bioinformatics analysis

D100 organoids were transferred from culture plates, briefly washed in ice-cold DPBS (PAN-Biotech, #P04-36500), centrifuged for 1 min at 600 × *g* at 4°C, and DPBS was removed completely. Afterwards, organoids were lysed in a buffer consisting of 4% SDS, 0.1 M DTT, and 0.1 M Tris pH 7.4, supplemented with protease and phosphatase inhibitors. For each biological replicate, 3–4 organoids were pooled and lysed together. Samples were sonicated (Digital Sonifier 250, Marshall Scientific), heated at 95°C for 3 min, and centrifuged for 15 min at 16,000 × g. Protein concentration was measured using tryptophan fluorescence. Albumin standards were prepared in a concentration range of 0–2000 μg/mL to generate a standard curve. For each sample, 10 μL of lysate was diluted into 490 μL of urea buffer (8 M urea in 10 mM HEPES pH 8.5) in a quartz cuvette, and absorbance was measured using a spectrofluorometer (FP-8300, Jasco).

#### Sample Preparation

Samples were subjected to an in-solution tryptic digest, following a modified version of the Single-Pot Solid-Phase-enhanced Sample Preparation (SP3) technology^25,26^. 20 µL of a slurry of hydrophilic and hydrophobic Sera-Mag Beads (Thermo Scientific, #4515-2105-050250, 6515-2105-050250) were mixed, washed with water and were then reconstituted in 100 µL water. 5 µL of the prepared bead slurry were added to 50 µL of the eluate following the addition of 55 µL of acetonitrile. All further steps were prepared using the King Fisher Apex System (Thermo Scientific). After binding to beads, beads were washed three times with 100 µl of 80% ethanol before they were transferred to 100 µL of digestion buffer (50 mM HEPES/NaOH pH 8.4 supplemented with 5 mM TCEP, 20 mM chloroacetamide (Sigma-Aldrich, #C0267), and 0.25 µg trypsin (Promega, #V5111)). Samples were digested over night at 37°C, beads were removed, and the remaining peptides were dried down and subsequently reconstituted in 10 µl of water. Peptides were reconstituted in 10 µL of H2O and reacted for 1 h at room temperature with 80 µg of TMT6plex (Thermo Scientific, #90066) label reagent dissolved in 4 µL of acetonitrile. Excess TMT reagent was quenched by the addition of 4 µL of an aqueous 5% hydroxylamine solution (Sigma, 438227). Peptides were reconstituted in 0.1 % formic acid and mixed to achieve a 1:1 ratio across all TMT-channels. Mixed peptides were purified by a reverse phase clean-up step (OASIS HLB 96-well µElution Plate, Waters #186001828BA). Peptides were subjected to an off-line fractionation under high pH conditions^25^. The resulting 12 fractions were then analysed by LC-MS/MS on an Orbitrap Fusion Lumos mass spectrometer (Thermo Scentific). LC-MS/MS analysis. Peptides were analysed by LC-MS/MS on an Orbitrap Fusion Lumos mass spectrometer (Thermo Scentific). To this end, peptides were separated using an Ultimate 3000 nano RSLC system (Dionex) equipped with a trapping cartridge (Precolumn C18 PepMap100, 5 mm, 300 μm i.d., 5 μm, 100 Å) and an analytical column (Acclaim PepMap 100. 75 × 50 cm C18, 3 mm, 100 Å) connected to a nanospray-Flex ion source. The peptides were loaded onto the trap column at 30 µl per min using solvent A (0.1% formic acid) and eluted using a gradient from 2 to 38% Solvent B (0.1% formic acid in acetonitrile) over 90 min at 0.3 µl per min (all solvents were of LC-MS grade). The Orbitrap Fusion Lumos was operated in positive ion mode with a spray voltage of 2.4 kV and capillary temperature of 275 °C. Full scan MS spectra with a mass range of 375–1500 m/z were acquired in profile mode using a resolution of 60,000 (maximum fill time of 50 ms; AGC Target was set to Standard) and a RF lens setting of 30%. Fragmentation was triggered for 3 s cycle time for peptide like features with charge states of 2–7 on the MS scan (data-dependent acquisition). Precursors were isolated using the quadrupole with a window of 0.7 m/z and fragmented with a normalized collision energy of 36%. Fragment mass spectra were acquired in profile mode and an orbitrap resolution of 15,000. Maximum fill time was set to 54 ms. AGC target was set to 200%. The dynamic exclusion was set to 60 s.

#### Data analysis

Acquired data were analysed using FragPipe^27^ and a Uniprot Homo sapiens fasta database (UP000005640, ID9606, 20.594 entries, date: 26.10.2022, downloaded: 11.01.2023) including common contaminants. The following modifications were considered: Carbamidomethyl (C, fixed), TMT10plex (K, fixed), Acetyl (N-term, variable), Oxidation (M, variable) and TMT6plex (N-term, variable). The mass error tolerance for full scan MS spectra was set to 10 ppm and for MS/MS spectra to 0.02 Da. A maximum of 2 missed cleavages were allowed. A minimum of 2 unique peptides with a peptide length of at least seven amino acids and a false discovery rate below 0.01 were required on the peptide and protein level (PMID: 25987413).

#### Bioinformatics analysis

Raw output files from FragPipe were processed using the R programming environment. Initial filtering removed reverse proteins and known contaminants. Only proteins with at least two quantified razor peptides (Razor.Peptides ≥ 2) were retained for downstream analysis, yielding 1,738 high-confidence proteins. TMT reporter ion intensities (log₂-transformed) were corrected for batch effects using the removeBatchEffect function from the limma package^28^. Data normalization was performed using variance stabilization normalization (normalizeVSN)^29^ from the same package. Differential expression analysis was conducted using limma’s moderated t-test, with replicate information included as a factor in the design matrix. Proteins were deemed differentially expressed (“hits”) if they had a False Discovery Rate (FDR) <0.05 and absolute fold change >2. Proteins with FDR <0.2 and absolute fold change >1.5 were considered candidate changes.

Gene ontology (GO) enrichment analysis was performed with the compareCluster function from the clusterProfiler R package^30^, using the org.Hs.eg.db database as reference. GO terms were analyzed for over-representation across Biological Process (BP), Molecular Function (MF), and Cellular Component (CC) categories. Enrichment significance was assessed using odds ratios, calculated from the ratio of term-associated genes in the dataset to those in the background set (GeneRatio vs BgRatio), with values >1 indicating enrichment. Network analyses of differentially abundant proteins were performed using Metascape (metascape.org)^31^. Protein lists (upregulated and downregulated hits, FDR ≤0.05, |log₂FC| ≥1) were submitted to Metascape’s Express Analysis workflow for protein-protein interaction (PPI) network construction using integrated databases including STRING, BioGRID, and OmniPath. Dense network components were identified using the MCODE algorithm with default parameters (degree cutoff = 2, node score cutoff = 0.2, k-core = 2, max depth = 100), and each MCODE complex was automatically annotated with enriched biological functions through Metascape’s integrated pathway databases.

### RNA sequencing and bioinformatics analysis

D100 organoids were transferred from culture plates to 1.5 mL tubes and briefly washed in ice-cold DPBS. For each biological replicate, 3 organoids were pooled in the same tube. After complete removal of DPBS, organoids were homogenized using QIAshredder homogenizers (Qiagen, #79656), and total RNA was extracted using the RNeasy Micro Kit (Qiagen, #74004) according to the manufacturer’s instructions. RNA was suspended in RNase-free water, and the concentration and purity of each sample were determined using a NanoDrop instrument (Thermo Fisher Scientific, NanoDrop One C).

Library preparation and RNA sequencing were performed as a service by GENEWIZ/AZENTA and sequenced on a NovaSeq 6000 instrument (Illumina). Sequence reads were trimmed to remove adapter sequences and low-quality nucleotides using Trimmomatic v0.36. Trimmed reads were mapped to the Homo sapiens GRCh38 reference genome available on ENSEMBL using the STAR aligner v2.5.2b. BAM files were generated at this step. Unique gene hit counts were calculated using featureCounts from the Subread package v1.5.2. Hit counts were summarised and reported using the gene_id feature in the annotation file. Only unique reads that mapped to exon regions were included.

After extraction, the gene hit count table was used for downstream differential expression analysis. Using DESeq2, a comparison of gene expression between Trip. and Isog. sample groups was performed. The Wald test was used to compute p-values and log₂ fold changes. Genes with an adjusted p-value <0.05 and an absolute log₂ fold change > 1 were considered differentially expressed genes (DEGs) in each comparison. Gene ontology (GO) analysis was performed on the statistically significant gene set using GeneSCF v1.1-p2. The goa_human GO list was used to cluster genes based on biological process terms and determine their statistical significance.

### Ribosome Profiling and bioinformatics analysis

Ribosome profiling was performed using the ALL-In-ONE RiboLace Gel-Free Kit (Immagina, #GF001) with modifications. *Sample preparation.* Organoids were incubated in maturation medium supplemented with 10 μg/mL cycloheximide (CHX) for 5 min, washed twice with cold DPBS containing 20 μg/mL CHX, and snap-frozen in liquid nitrogen. Ten frozen organoids per sample were pooled and pulverized under liquid nitrogen using a mortar and pestle. The powder was collected in a 1.5 mL tube and resuspended in 400 μL Tissue Lysis Buffer (Immagina, #RL001-02) supplemented with 10% sodium deoxycholate (Thermo Scientific, #89904), 1 U/μl DNase I (Invitrogen, #AM2222), and 40 U/μl RiboLock RNase Inhibitor (Thermo Fisher Scientific, #EO0381), and centrifuged according to the manufacturer’s protocol. The absorbance units (AU) of the samples were measured using a NanoDrop spectrophotometer at 260 nm. *Ribosome-protected fragment pulldown.* High Affinity ribosome beads (hiRB; Immagina, #GF001-04) were prepared for pulldown according to the functionalization protocol. Tissue lysates were subjected to nuclease digestion during the bead functionalization step. A fraction of each lysate was retained as an internal mRNA control (total mRNA). The digested lysates were incubated with functionalized beads for 70 min to allow ribosome binding. Ribosome-protected fragments (RPFs; length: 28–32 nt) were then purified using RNA Clean & Concentrator-5 (Zymo Research, #R1015). *Library preparation.* RPF library preparation was carried out according to the manufacturer’s instructions and included the following steps: A. 5′ phosphorylation, B. Adapter ligation, C. Circularization, D. Reverse transcription, E. PCR amplification, F. PCR amplification 2, G. Library quality check, H. Sequencing. Sequencing was performed as a service by Immagina Biotechnology (Italy) on a NovaSeq 6000 instrument (Illumina). *Bioinformatics analysis*. Preprocessing and P-site determination. Raw ribosome profiling reads were processed using the riboWaltz package (v2.0). Following adapter trimming and size selection (28-32 nt), reads were aligned to the GRCh38 transcriptome using STAR (v2.7.10b). riboWaltz’s psite function was employed to calculate read-length specific P-site offsets using a flanking region of 6 nucleotides around start codons, with automatic extremity selection (extremity=“auto”). This generated corrected P-site positions that showed strong 3-nt periodicity across coding sequences (CDS), confirming proper ribosome positioning. Translational efficiency analysis. P-site-mapped reads were analyzed using Xtail (v1.2.0) to identify genes with differential translation between *SNCA* Trip and Isog organoids. RNA-seq counts (TPM) were paired with ribosome-protected fragment (RPF) counts, normalized via median-of-ratios, and analyzed using Xtail’s dual statistical framework evaluating both ratio-of-fold-changes (RFC) and fold-change-of-ratios (FCR). Genes with FDR <0.1 in either metric were considered significant. Gene ontology analysis was performed using g:profiler^32^ and SynGO^33^ as described in ref.^34^ Word clouds were generated using the Python wordcloud library (version 1.9.4) in combination with Matplotlib (version 3.10.3).

### ECM staining, imaging and analysis

Midbrain organoids at D50 and D100 were fixed and stained with Wisteria floribunda agglutinin (WFA) to visualize extracellular matrix (ECM) components, and with MAP2 to label neuronal somas, as described in the Immunofluorescence and Confocal Imaging section. Confocal z-stack images were acquired and processed using ImageJ (NIH). Neuronal somas were identified based on MAP2 signal, and binary masks were generated for each soma. Using the freehand selection tool, a region of interest (ROI) was drawn around the soma to define the base area (Z₀). This ROI was then expanded by 1 μm (Z₁) and 4 μm (Z₂) to create concentric perisomatic rings around each neuron. The Z₁ ring was defined as the pericellular ECM compartment (corresponding to perineuronal nets; PNN), while the region between Z₁ and Z₂ (i.e., Z₂–Z₁) was defined as the interstitial ECM compartment. Masks corresponding to these regions were overlaid onto the WFA fluorescence channel to measure integrated intensity, allowing for quantification of ECM localization in pericellular versus interstitial zones.

### Reverse transcription-quantitative PCR (RT-qPCR)

Total RNA was extracted from *SNCA* Trip and isogenic iPSCs using TRI Reagent (Sigma-Aldrich, #T9424) according to the manufacturer’s instructions. RT-qPCR was performed using a two-step protocol with the LunaScript RT SuperMix Kit (NEB, #E3010) and the Luna Universal qPCR Master Mix (NEB, #M3003) on an AriaMx Real-Time PCR System (Agilent Technologies, G8830A). Raw Ct values were normalized to GAPDH using the ΔΔCt method. Primer sequences (F: forward, R: reverse) used in this study are as follows:

OCT4: F: 5΄-GGAGGAAGCTGACAACAATGAAA-3΄, R: 5΄-GGCCTGCACGAGGGTTT-3΄,

NANOG: F: 5΄-ACAACTGGCCGAAGAATAGCA-3΄, R: 5΄-GGTTCCCAGTCGGGTTCAC-3΄,

GAPDH: F: 5΄-ACCACAGTCCATGCCATCAC-3΄, R: 5΄-TCCACCACCCTGTTGCTGTA-3΄.

## Results

### A human iPSC-derived *SNCA* triplication midbrain organoid model

To investigate early pathological mechanisms associated with *SNCA* gene triplication and their link to proteostasis in 3D brain organoids, we utilized a human midbrain organoid model derived from induced pluripotent stem cells (iPSCs) carrying an *SNCA* triplication (*SNCA* Trip) and the isogenic CRISPR/Cas9-corrected counterpart (*SNCA* Isog)^14^. This model was established using a differentiation protocol previously described^14^ and subsequently adopted in related studies^15^. We selected day 50 (D50) and day 100 (D100) of organoid maturation for analysis, aiming to capture early cellular alterations prior to the onset of overt neurodegeneration. This approach was motivated by evidence from animal models and patient-derived neurons indicating that proteostasis dysregulation, including aberrant activation of mTORC1 signalling, is implicated in the pathogenesis of synucleinopathies^6^. The *SNCA* Trip and *SNCA* Isog lines exhibited comparable expression of pluripotency markers OCT3/4 and Nanog at the iPSC stage, as confirmed by quantitative PCR analyses and immunofluorescence (Fig. 1A, B). Differentiation into midbrain organoids followed a stepwise protocol involving neural induction, midbrain patterning, and tissue induction phases, leading to long-term organoid maintenance up to 100 days in culture (Fig. 1C). Immunofluorescence analysis at D50 revealed the presence of TH-positive dopaminergic neurons and MAP2-positive neuronal networks in both genotypes, indicating successful midbrain specification (Fig. 1D). Western blot analysis showed significantly elevated α-synuclein protein levels in *SNCA* Trip organoids compared to *SNCA* Isog at both D50 (314.29% increase) and D100 (117.9% increase) (Fig. 1E), together with increased phosphorylation of SNCA S129 (Sup. Fig. 1B). Morphological assessment across differentiation stages demonstrated consistent organoid formation between lines, with no significant differences observed in organoid diameter or projected area at D50 (Fig. 1F). However, by D100, *SNCA* Trip organoids displayed a modest but statistically significant reduction in both parameters (area: 13.86%; diameter: 7.19%) relative to *SNCA* Isog controls (Fig. 1F). Furthermore, spontaneous neuromelanin deposition, a hallmark of maturing midbrain dopaminergic neurons, was observed in *SNCA* Trip organoids at D100 but was less pronounced in *SNCA* Isog controls (Fig. 1G).

**Figure 1.**
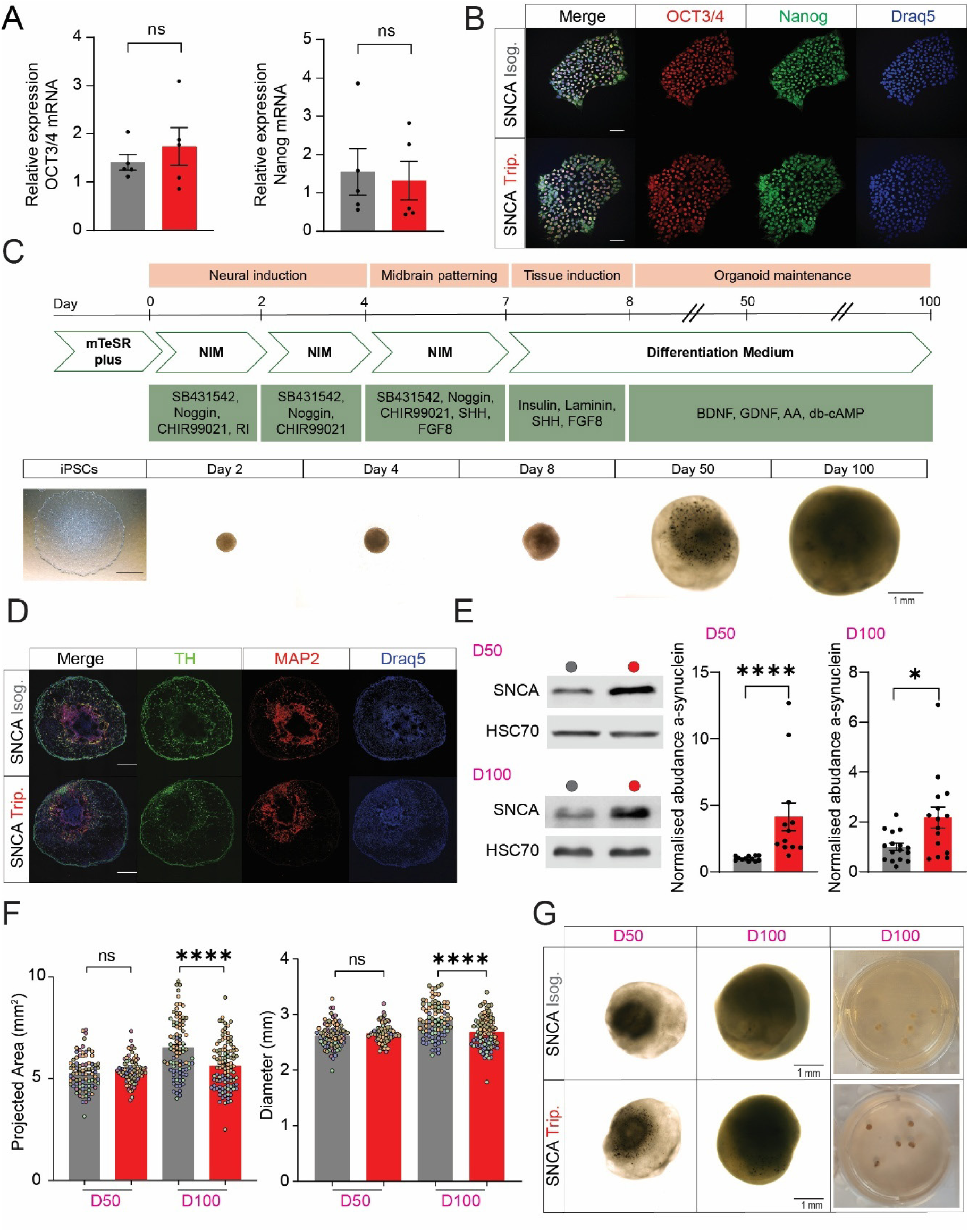
Human midbrain organoid model of *SNCA* triplication recapitulates early synucleinopathy features. (A) qPCR analysis of pluripotency markers *OCT3/4* and *Nanog* in *SNCA* Trip and *SNCA* Isog iPSCs. (B) Immunofluorescence staining of OCT3/4 (red) and Nanog (green) in *SNCA* Trip and *SNCA* Isog iPSCs; nuclei counterstained with Draq5 (blue). (C) Schematic illustration of the midbrain organoid differentiation protocol including neural induction, midbrain patterning, and organoid maintenance stages. Brightfield images depict representative morphological development from iPSCs up to D100 organoids. (D) Immunofluorescence staining of D50 organoid cryosections for TH (green), MAP2 (red), and Draq5 (blue), showing dopaminergic and neuronal populations. Scale bar = 500 μm. (E) Immunoblot analysis of α-synuclein levels in D50 and D100 organoids. Left: representative immunoblots. Right: quantification of α-synuclein protein levels normalized to HSC70 (D50; n = 12 per group; 3 organoid batches, D100; n= 15-16 per group; 3 organoid batches). (F) Quantification of projected surface area and diameter in D50 and D100 organoids, showing no differences at D50 and reduced size in *SNCA* Trip organoids at D100. (G) Brightfield images of organoids at D50 and D100, with high-magnification images at D100 showing spontaneous neuromelanin accumulation in *SNCA* Trip organoids. Scale bars as indicated. One-way ANOVA or Student’s *t*-test with parametric or non-parametric (*Mann-Whitney*) test. Data shown as mean ± SEM. \**P* <0.05, \*\*\*\**P* <0.0001; ns, not significant. See also Sup. Fig. 1 and Sup. Table 5 for details on statistical analyses.

Collectively, these results establish that the *SNCA* Triplication midbrain organoid model recapitulates key features of early synucleinopathy, including α-synuclein accumulation and S129 phosphorylation together with neuromelanin formation, within the first 100 days of *in vitro* development.

### Dysregulated signalling linked to proteostasis in D100 *SNCA* Trip organoids

We next examined signalling pathways regulating proteostasis, focusing on key mediators of protein synthesis and cellular stress responses (Fig. 2A, E). Western blot analysis revealed a significant increase in phosphorylation of ribosomal protein S6 at serines 240/244 (p-rpS6 S240/244), indicative of mTORC1 activation, in SNCA Trip organoids compared to *SNCA* Isog controls at both D50 (42.93%) and D100 (37.25%) (Fig. 2B). In contrast, phosphorylation at serines 235/236 (p-rpS6 S235/236), which is mostly mediated by RSK^35^ and to a smaller extent by mTORC1, remained unchanged at both timepoints (Fig. 2B).

**Figure 2.**
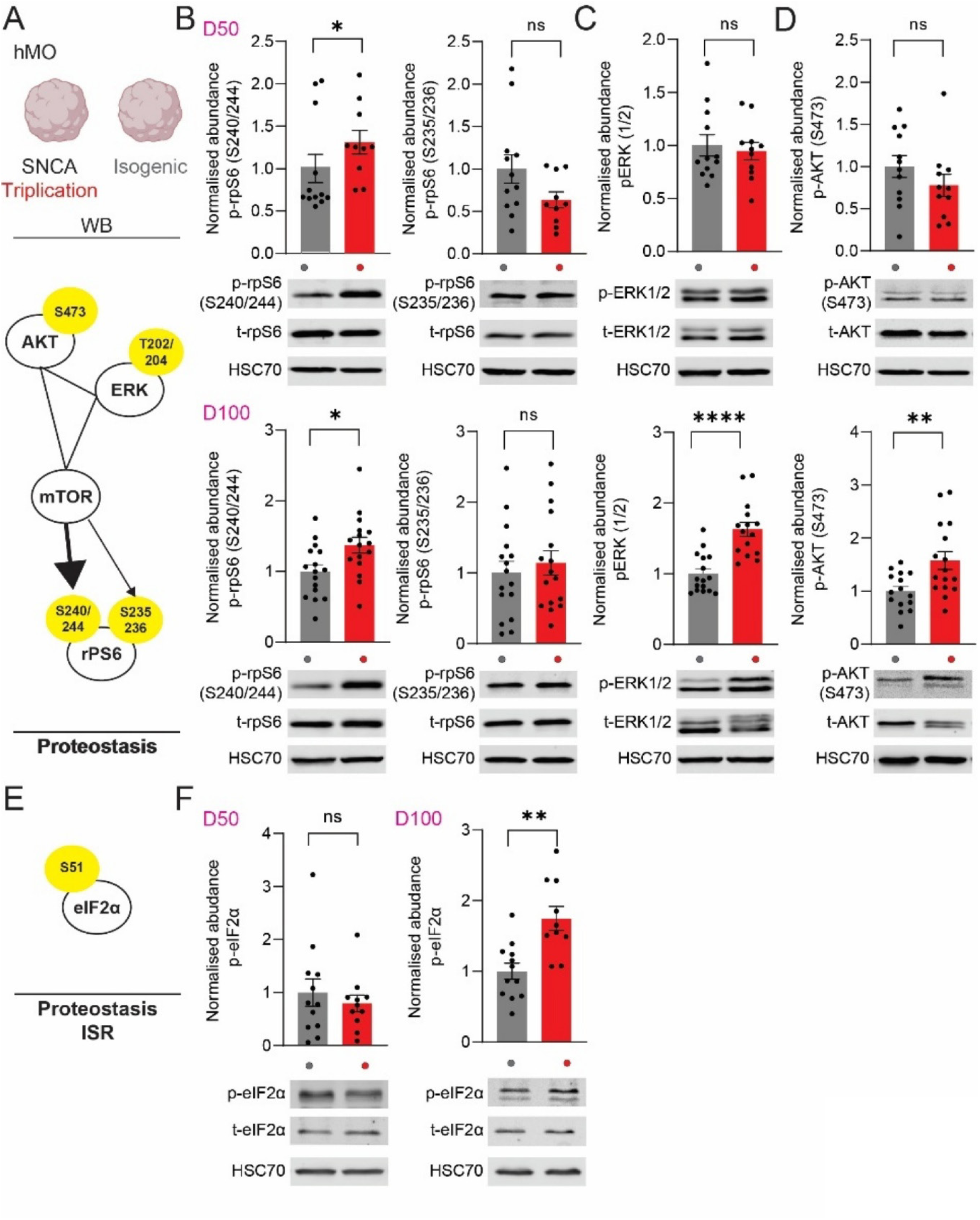
Proteostasis and signalling pathway alterations in *SNCA* Triplication midbrain organoids at D50 and D100. (A) Schematic illustration of the “AKT-ERK-mTOR” triangle of signaling upstream of proteostasis. Immunoblot analysis of PI3K/AKT/mTOR signalling in D50 and D100 isogenic and triplication organoids. Quantification of (B) phospho-rpS6 (S240/244, 235/236), (C) phospho-ERK 1/2 and (D) phospho-AKT (S473) for the indicated groups (*n* = 10-16 per group, 3 organoid batches for D50 and D100. (E) eIF2α signalling and ISR, proteostasis. (F) Quantification of phospho-eIF2α for the indicated groups (*n* = 10-12pergroup, 3 organoid batches for D50 and D100. For B, C, D, F (bottom): Representative immunoblots of midbrain organoids lysates, probed with antisera against the indicated proteins. HSC70/GAPDH: loading control. Student’s *t*-test with parametric or non-parametric (*Mann-Whitney*) test. All data are shown as mean ± SEM. \**P* <0.05, \*\**P* <0.01, \*\*\*\**P* <0.0001. See also Sup. Fig. 2 for raw WB data and Sup. Table 5 for details on statistical analyses.

At D50, no differences were observed in ERK1/2 (p-ERK T202/Y204) or AKT (p-AKT S473) phosphorylation (Fig. 2C, D). However, by D100, *SNCA* Trip organoids exhibited pronounced upregulation of p-ERK1/2 (75.45%) and p-AKT (57.75%), suggesting broader activation of growth and survival pathways as maturation progressed (Fig. 2C, D). In parallel, phosphorylation of eukaryotic initiation factor 2 alpha (p-eIF2α), a marker of integrated stress response (ISR) activation, was significantly elevated in *SNCA* Trip organoids at D100 (74.68% increase) but not at D50 (Fig. 2F).

These findings indicate that *SNCA* triplication leads to early and progressive dysregulation of signalling pathways linked to proteostasis, characterized by mTORC1 hyperactivation at prodromal stages, followed by broader perturbations in mTOR, AKT, ERK and eIF2α signalling, as organoids mature.

### Proteomic alterations reveal disrupted cytoskeletal, synaptic, and extracellular matrix pathways in *SNCA* Trip organoids

Given the changes in signalling upstream of proteostasis at D100, we performed quantitative proteomic analysis of D100 midbrain organoids using tandem mass tag (TMT) labelling and liquid chromatography–mass spectrometry (LC-MS/MS) (Fig. 3A). Principal component analysis of the proteomics dataset confirmed robust separation between *SNCA* Trip and *SNCA* Isog organoids (Supplementary Figure 3A). Differential abundance analysis identified 187 proteins significantly upregulated, and 79 proteins significantly downregulated in *SNCA* Trip organoids relative to *SNCA* Isog controls (FDR ≤ 0.05; |log₂FC| ≥ 1), with an additional set of candidates showing moderate fold changes (Fig. 3B).

**Figure 3.**
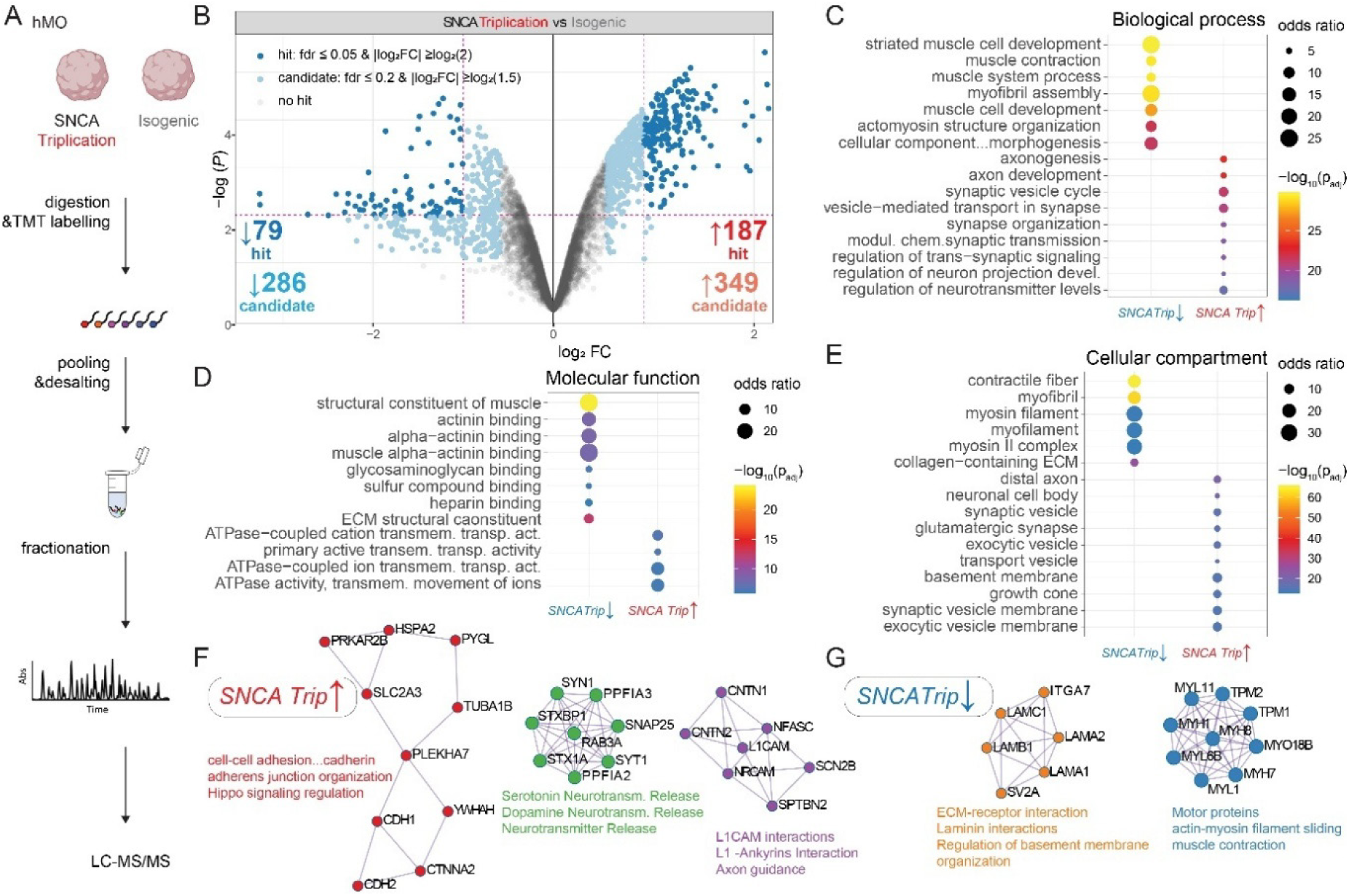
Global TMT proteomic analysis identifies altered neuronal and ECM-associated pathways in *SNCA* Trip organoids. (A) Schematic overview of the tandem mass tag (TMT)-based quantitative proteomics workflow used for analysis of D100 *SNCA* Trip and *SNCA* Isog midbrain organoids. (B) Volcano plot of differentially abundant proteins in *SNCA* Trip vs. *SNCA* Isog organoids. Proteins with false discovery rate (FDR) ≤0.05 and |log₂ fold change| ≥log₂(2) are considered significantly changed (blue = downregulated, red = upregulated); proteins with FDR ≤0.2 and |log₂FC| ≥ log₂(1.5) are shown as candidate changes (light blue/red). (C–E) Gene ontology (GO) enrichment analyses of significantly altered proteins categorized by biological process (C), molecular function (D), and cellular compartment (E). Circle size reflects odds ratio, and colour denotes adjusted p-value (*Padj*). (F–G) Metascape protein–protein interaction (PPI) network analysis of upregulated (F) and downregulated (G) proteins. Upregulated modules (F) include cadherin-mediated cell adhesion, Hippo signalling, and neurotransmitter release pathways. Downregulated modules (G) include ECM-receptor interaction, laminin complexes, axon guidance, and cytoskeletal motor proteins. See also Sup. Fig. 3 and Sup Table 1

Gene ontology enrichment analysis of proteomics hits revealed that upregulated proteins in *SNCA* Trip organoids (*SNCA* Trip↑) were primarily associated with neuronal functions, including synaptic vesicle cycling, neurotransmitter release, synapse organization, and axon development (Fig. 3C). These were reflected in molecular function categories such as ATPase-coupled transport, and cellular compartments such as synaptic vesicles, distal axons, and glutamatergic synapses (Fig. 3D–E). In contrast, downregulated proteins (SNCA Trip↓) were enriched in ECM-associated and actomyosin-related pathways, including muscle system processes, actin–myosin structure organization, and cytoskeletal and actin-binding activities, including alpha-actinin and glycosaminoglycan binding. These were accompanied by reduced representation in ECM structural components and contractile elements such as myofibrils, contractile fibres, and collagen-containing ECM (Fig. 3C–E). SynGO analysis revealed that *SNCA* Trip upregulated proteins were significantly enriched in synaptic compartments, particularly postsynaptic densities, whereas *SNCA* Trip downregulated proteins showed minimal synaptic annotation (Sup. Fig. 3C).

Network analyses of proteins upregulated in *SNCA* Trip organoids proteins using Metascape identified protein networks involved in cadherin-mediated cell-cell adhesion, Hippo signalling and serotonin and dopamine neurotransmitter release (Fig. 3F, left panel), as well as the actin/spectrin cytoskeleton and axon guidance (Fig. 3F, right panel). In contrast, networks of proteins downregulated in *SNCA* Trip organoids implicated key ECM components, laminin-receptor interactions, and motor protein complexes essential for cytoskeletal integrity (Fig. 3G).

These findings reveal that *SNCA* triplication induces broad proteomic changes, affecting cytoskeletal dynamics, synaptic/neurotransmitter function, and ECM organization at D100, concomitant with mTOR, ERK, AKT and eIF2α pathways hyperactivation.

### Transcriptional changes in D100 *SNCA* Trip organoids align with proteomic alterations

To complement the proteomic findings, we performed bulk RNA sequencing on D100 *SNCA* Trip and *SNCA* Isog midbrain organoids (Fig. 4A). Principal component analysis confirmed distinct transcriptional profiles between SNCA Trip and SNCA Isog organoids at D100 (Sup. Fig. 4A, B). Differential expression analysis revealed 397 transcripts (differentially expressed genes; DEG) significantly upregulated and 157 transcripts significantly downregulated (*Padj* ≤ 10⁻⁴, |log₂FC| ≥2), with an additional set of candidate genes showing moderate fold changes (Fig. 4B).

**Figure 4.**
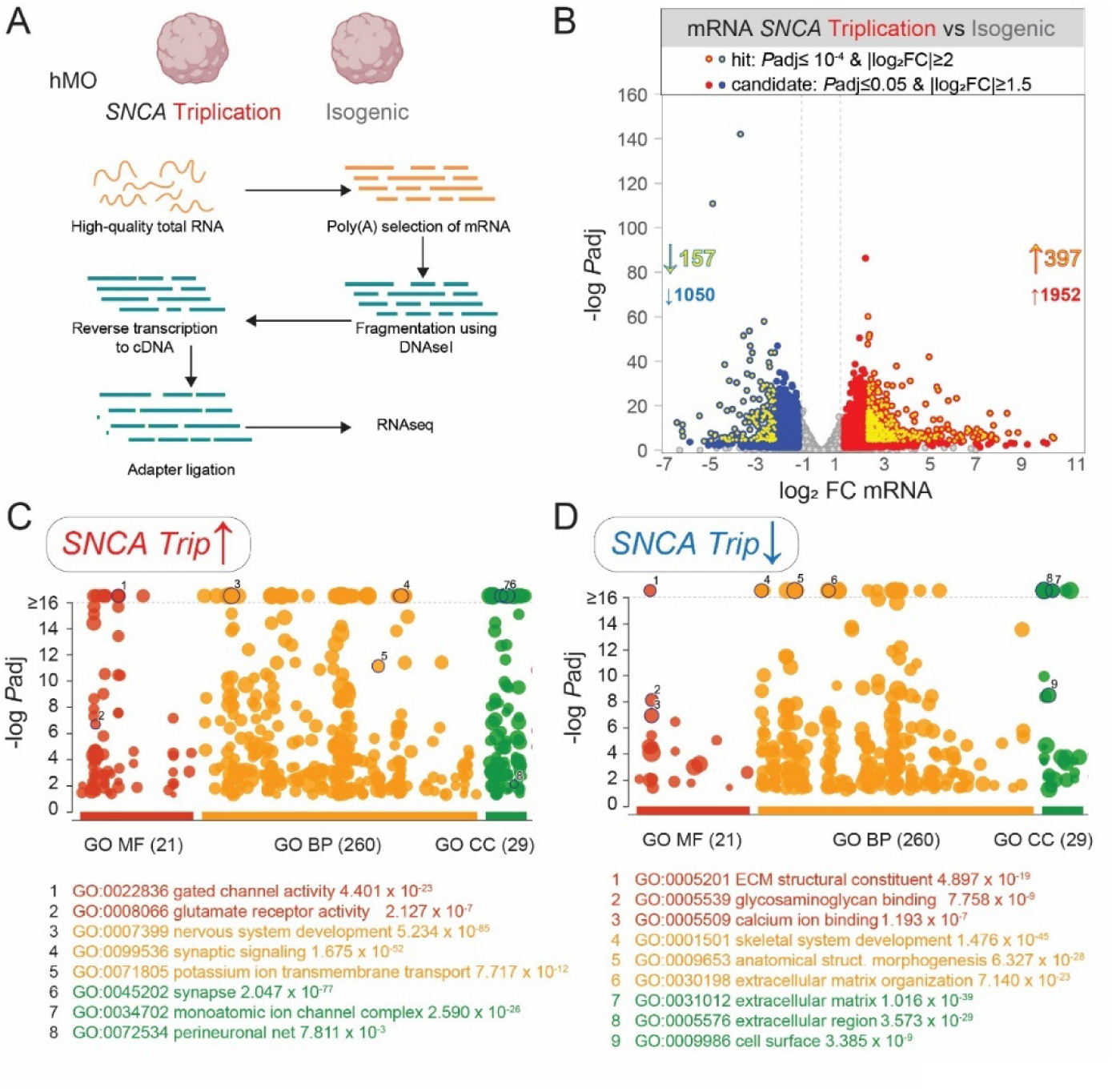
Transcriptomic profiling links *SNCA* triplication to synaptic activation and ECM suppression. (A) Schematic overview of the bulk RNA sequencing workflow used to compare gene expression profiles between D100 *SNCA* Trip and *SNCA* Isog midbrain organoids. (B) Volcano plot showing differential gene expression (*SNCA* Trip vs. *SNCA* Isog). Genes with adjusted p-value (*Padj*) ≤10⁻⁴ and |log₂ fold change|≥ 2 are shown as significant hits (red = upregulated, blue = downregulated); additional genes with *Padj* ≤0.05 and |log₂FC| ≥1.5 are shown as candidates. (C–D) Gene ontology (GO) enrichment analysis of significantly upregulated (C) and downregulated (D) transcripts categorized by molecular function (MF), biological process (BP), and cellular component (CC). Select top terms are annotated, highlighting enrichment in synaptic signalling, ion channel activity and perineuronal nets (PNN) among upregulated genes (C), and structural ECM components among downregulated genes (D). Circle size represents gene set size; vertical position reflects significance (-log₁₀*Padj*). See also Sup. Fig. 4 and Sup Table 2.

Gene ontology analysis of upregulated DEG in *SNCA* Trip organoids highlighted enrichment in biological processes related to neuronal signalling, including gated channel activity, glutamate receptor activity, nervous system development, and synaptic signalling (Fig. 4C). These findings are consistent with the proteomic evidence of enhanced synaptic and axonal pathways. Conversely, downregulated DEG were predominantly associated with ECM structure and organization, including decreased expression of genes involved in ECM-receptor interactions, glycosaminoglycan binding, and skeletal system development (Fig. 4D), mirroring the loss of ECM components observed in the proteome. Interestingly genes coding for perineuronal net (PNN) proteins such as brevican (BCAN), tenascin R (TNR), aggrecan (ACAN), neurocan (NCAN) and versican (VCAN), were among the upregulated DEG (Sup. Table 2). SynGO analysis revealed that upregulated DEG were significantly enriched in synaptic gene ontologies, contrasting with minimal synaptic representation among downregulated DEG (Sup. Fig. 4C) Together, the transcriptomic data corroborate the proteomic alterations, indicating that *SNCA* triplication perturbs both neuronal signalling and ECM/PNN homeostasis at the transcriptional level in midbrain organoids.

### Translational landscape remains largely unchanged in *SNCA* Trip organoids but reveals evidence of buffering

To investigate whether the observed proteomic alterations were driven by changes in mRNA translation, which is a key determinant of proteostasis, we performed ribosome profiling (Ribo-seq) on D100 *SNCA* Trip and *SNCA* Isog organoids (Fig. 5A). Translational efficiency (TE) was assessed by integrating Ribo-seq and RNA-seq data using the XTAIL framework (Fig. 5B).

**Figure 5.**
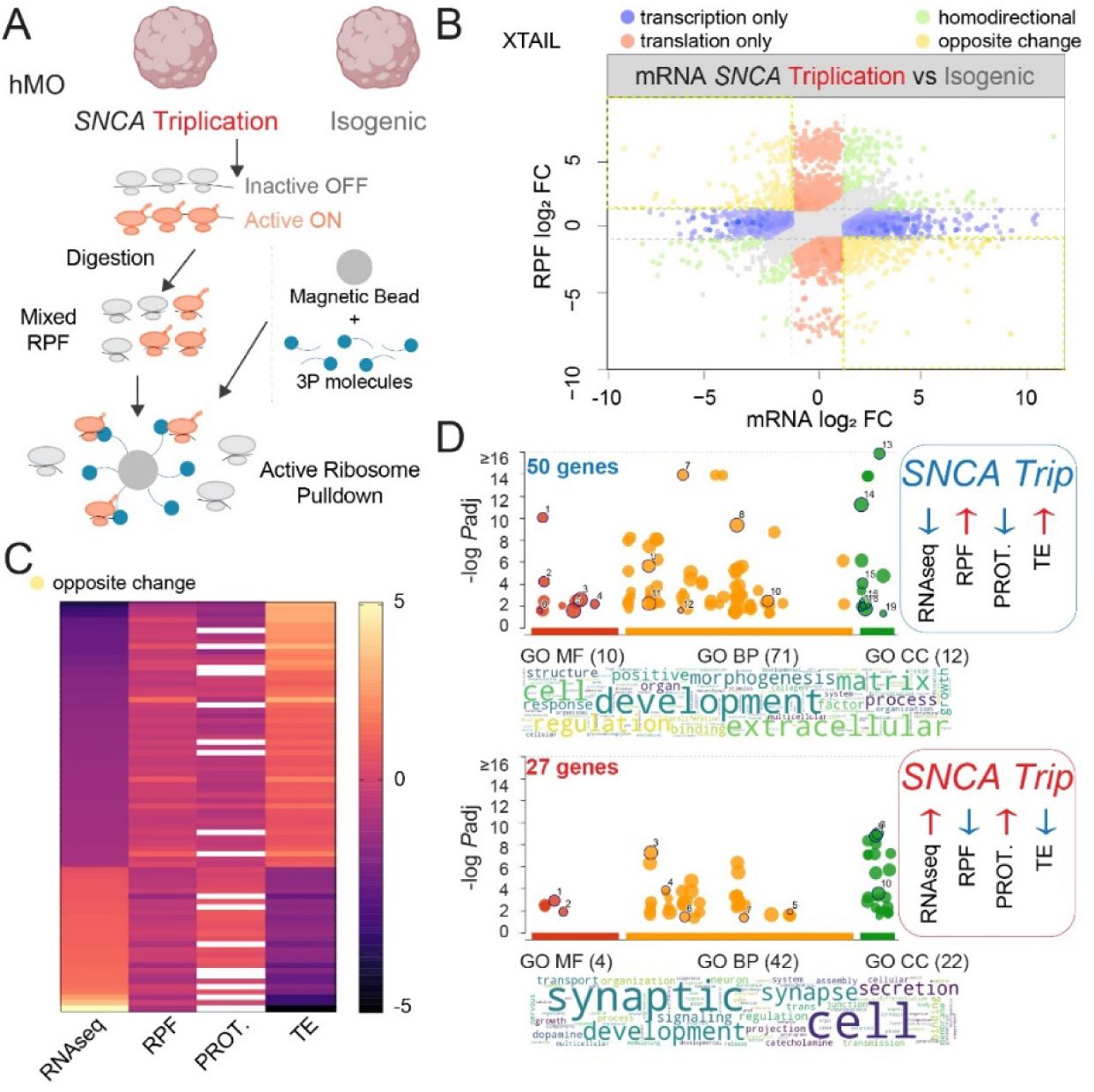
Ribosome profiling reveals selective translational buffering of ECM and synaptic transcripts in *SNCA* Trip organoids. (A) Schematic overview of the active ribosome profiling workflow used to isolate ribosome-protected fragments (RPFs) and identify actively translated transcripts in *SNCA* Trip and *SNCA* Isog D100 midbrain organoids. (B) Biplot comparing changes in mRNA abundance (RNA-seq, x-axis) and ribosome occupancy (RPF, y-axis). Transcripts showing transcription-only (blue), translation-only (red), homodirectional (orange), or opposite (yellow) changes are annotated based on XTAIL classification. (C) Heatmap showing gene-level changes across RNA abundance (RNA-seq), ribosome occupancy (RPF), protein abundance (PROT.), and translational efficiency (TE), highlighting transcripts with evidence of opposite-direction changes. (D) Gene ontology enrichment analysis of buffered transcripts using g:profiler, showing either increased (top) or decreased (bottom) translational efficiency in *SNCA* Trip organoids. Word clouds represent key enriched biological themes; GO terms are plotted by adjusted p-value across molecular function (MF), biological process (BP), and cellular component (CC). Summary insets indicate general directional trends for RNA, RPF, protein, and TE values for each gene group. See also Sup. Fig. 5 and Sup Table 3

Comparative analysis revealed that most changes in protein abundance could be attributed to transcriptional alterations, with minimal contribution from differential translation. Most transcripts exhibited homodirectional changes between mRNA and ribosome-protected fragment (RPF) levels (Fig. 5B), indicating concordant regulation at the transcriptional level. A subset of genes (77), however, demonstrated opposite regulatory patterns between RNA abundance and translation, suggestive of translational buffering (Fig. 5C). Gene ontology analysis of buffered transcripts highlighted processes related to neuronal development and extracellular matrix organization, with distinct subsets showing reduced or enhanced translational efficiency relative to RNA abundance (Fig. 5D).

These results indicate that, while *SNCA* triplication does not induce widespread changes in translational control, selective buffering mechanisms may modulate the translation of key transcripts involved in synaptic and extracellular matrix pathways.

### ECM disorganization in *SNCA* Triplication organoids precedes neurodegeneration

ECM is a critical regulator of brain architecture and function, providing structural support, modulating synaptic stability, and influencing neuronal plasticity^36^. In the midbrain, specialized ECM structures, including perineuronal nets, are essential for maintaining dopaminergic neuron health and protecting against neurodegenerative processes^37^. Given the transcriptomic, translatomic and proteomic evidence of ECM dysregulation in *SNCA* Trip organoids, we sought to examine ECM organization at the cellular level.

Using immunofluorescence and confocal imaging, we assessed ECM integrity by staining with Wisteria floribunda agglutinin (WFA), a marker of chondroitin sulfate proteoglycan-rich ECM and aggrecan (ACAN), a proteoglycan found in PNNs, in combination with neuronal markers MAP2 and TH (Fig. 6A, B). At D100, *SNCA* Trip organoids exhibited prominent WFA staining around MAP2^+^ neuronal somata (42.32% increase), indicative of increased pericellular ECM deposition (PNN), as well as enhanced interstitial ECM (44.07% increase) in the parenchyma, compared with *SNCA* Isog. (Fig. 6C, D).

**Figure 6.**
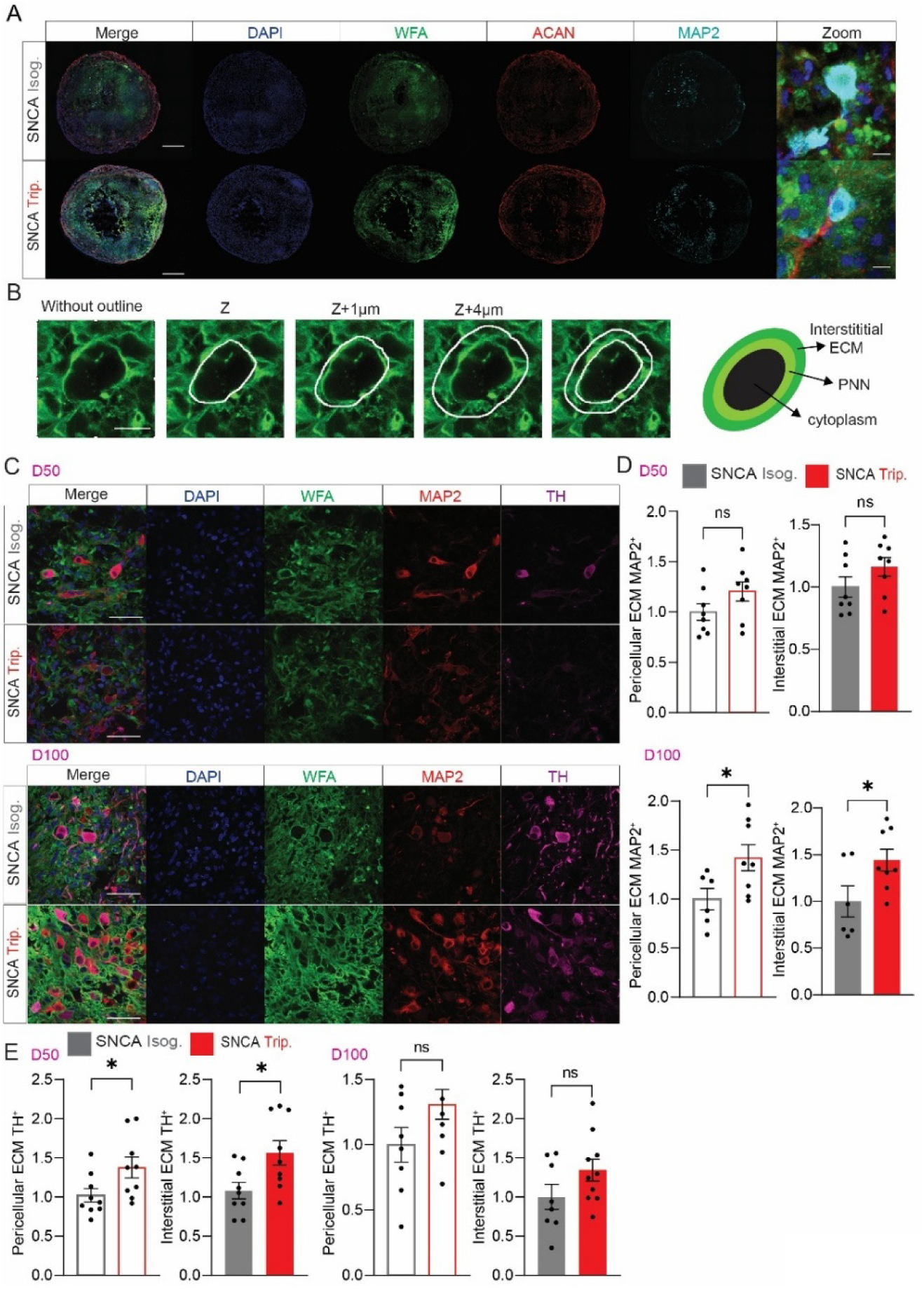
Disorganization and accumulation of pericellular and interstitial extracellular matrix (ECM) in SNCA Triplication midbrain organoids. (A) Representative confocal images of whole organoid cryosections at D100 stained with DAPI (nuclei, blue), WFA (ECM, green), aggrecan (ACAN, red), and MAP2 (neurons, cyan) in *SNCA* Isog and *SNCA* Trip organoids. Zoom panel highlights pericellular WFA^+^ ECM surrounding MAP2^+^ neuronal somata. Scale bars: 500 µm (whole section), 10 µm (zoom). (B) Orthogonal projection views (Z₀–Zₙ) showing 3D organization of pericellular ECM nets stained with WFA. White outlines delineate individual neuronal cell bodies encased by WFA^+^ ECM. (C) Representative higher-magnification confocal images of organoid parenchyma at D50 and D100 showing WFA (green), MAP2 (red), and TH (magenta) staining in *SNCA* Isog and *SNCA* Trip organoids. Quantification of perineuronal (PNN) and interstitial ECM signal intensity at D50 (top) and D100 (bottom) in (D) MAP2^+^ cells reveals significantly increased PNN and interstitial ECM deposition in *SNCA* Trip organoids at D100 but not at D50, and, in (E) TH^+^, significantly increased PNN and interstitial ECM deposition in *SNCA* Trip organoids at D50 and a strong trend for increase at D100. Data are shown as mean ± SEM. \**P* < 0.05, Student’s *t*-test. ns, not significant. Scale bars: 50 µm. See also Sup. Table 5 for details on statistical analyses.

Interstitial ECM area was similarly elevated (Fig. 6C, D). Notably, these changes were not observed at D50 (Fig. 6D), suggesting that ECM remodelling emerges progressively during organoid maturation during a period which is prodromal to pervasive neurodegeneration and cell death. Strikingly, compared to control, *SNCA* Trip TH^+^ cells displayed increased pericellular (35%) and interstitial ECM (44.85%) WFA staining from D50, with a strong trend for increased staining at D100 too (pericellular: 31%, interstitial: 34.2%), yet not statistically significant.

These findings demonstrate that *SNCA* triplication leads to abnormal accumulation and organization of ECM components around neurons, in a cell-type specific manner, supporting the notion that proteostatic imbalance extends beyond intracellular compartments to the extracellular environment in early stages of synucleinopathy.

## Discussion

In this study, we employed human midbrain organoids harbouring *SNCA* triplication as a predictive model for early synucleinopathy, revealing progressive proteostatic collapse, ECM dysregulation, and cell-type-specific vulnerabilities preceding neurodegeneration (D50-D100). By integrating multiomic profiling (Figs. 3, 4, 5) with ECM analysis with imaging, we identify: hyperactivation of PI3K/AKT/mTORC1-driven signalling cascades (Fig. 2) and transcriptional-translational decoupling (Fig. 5) as key mechanisms possibly linking SNCA accumulation to PNN and ECM remodelling in dopaminergic circuits.

### Proteostatic stress and signalling network dysregulation

*SNCA* Trip organoids recapitulate hallmark features of prodromal Parkinson’s disease (PD), including α-synuclein aggregation, neuromelanin deposition (Fig. 1), and mTORC1 hyperactivation (p-rpS6 S240/244) at D50–D100 (Fig. 2). While prior studies in *SNCA* models^7,14–19,21^ emphasized synaptic deficits, ER stress and mitochondrial dysfunction, our data reveal early co-activation of AKT/mTOR and ERK pathways (Fig. 2B–D), mirroring findings in LRRK2 G2019S models where dual kinase activation exacerbates proteotoxicity^38^. The delayed emergence of p-eIF2α elevation at D100 (Fig. 2F) suggests ISR activation occurs secondary to chronic proteostatic overload, consistent with iPSC-derived neuron studies showing temporal decoupling of UPR initiation and resolution^7^.

Despite evidence of AKT/mTORC1 hyperactivation and elevated ERK and p-eIF2α signalling at D100, our ribosome profiling analysis revealed minimal global changes in translational output (Fig. 5). Instead, we observed selective translational buffering, particularly in transcripts related to neuronal development and ECM organization (Fig. 5C). Plausibly, in early synucleinopathy, cells may engage compensatory translational mechanisms to maintain homeostasis in the face of transcriptional dysregulation, a phenomenon previously hinted at in models of neurodegeneration^39^ but not systematically demonstrated in *SNCA* triplication systems. On the other hand, bulk omics analysis performed herein may mask cell-type specific phenotypes, requiring single-cell analysis of gene expression.

### ECM remodelling as a compensatory and pathogenic driver

Transcriptomic and proteomic analyses converged on ECM dyshomeostasis, with PNN component upregulation (BCAN, TNR, ACAN) contrasting broad ECM protein downregulation (Fig. 3, 4). This divergence aligns with analyses in several PD models^40–42^, such as E326K-*GBA1* (Glucosylceramidase Beta 1) and *PINK1/PRKN* (PTEN-Induced Putative Kinase 1/ Parkin RBR E3 Ubiquitin Protein Ligase), collectively showing ECM gene expression modulation coupled with synaptic dysfunction. Our work goes beyond; spatial quantification revealed TH^+^ neurons develop pericellular/interstitial ECM accumulation by D50; 6 weeks earlier than MAP2^+^ populations (Fig. 6D–E). This cell-type-specific phenotype mirrors clinical neuropathology where dopaminergic neurons exhibit heightened ECM receptor (e.g., integrin α3β1) expression prior to Lewy pathology^43^. These findings underscore the value of 3D organoid models in studying structures such as the ECM, whereby planar systems e.g. 2D cultures, lack the 3D cytoarchitecture and stromal interactions necessary for matrix maturation. PNN-like structures develop in 2D neuronal cultures, but lack maturity and require longer culture time, however some recent models address this issue^44^.

### PNN dynamics in early pathogenesis

The downregulation of ECM-related proteins and transcripts, together with increased WFA staining surrounding MAP2^+^ neurons (Fig. 6), is in line with aberrant accumulation of perineuronal nets and interstitial ECM, that may reflect changes in ECM turnover and synthesis. Strikingly, this phenotype emerges sooner (D50 for TH^+^ cells). This ECM remodelling could be explained by changes in the expression of ECM-degrading enzymes in *SNCA* Trip, such as matrix metalloproteinases (MMPs) and ADAMTS proteases, which are found in our omics (Sup. Table 1, 2, 3). Upregulated PNN gene expression (BCAN, NCAN, VCAN) alongside postsynaptic density enrichment (Sup. Fig. 4C) suggests attempted synaptic stabilization, ascribing a neuroprotective role of PNN upregulation^45^. However, excessive perineuronal matrix deposition may entrap SNCA oligomers, creating a feedforward loop of proteostatic stress. The earlier ECM dysregulation in TH^+^ cells (Fig. 6E) may reflect their unique reliance on CSPG-mediated BDNF signalling^46^, which *SNCA* aggregates could disrupt via mechanisms such as cellular prior protein (PrP^C^) sequestration^47^.

### Implications for PD progression and therapeutic development

The D50–D100 window of ECM remodelling mirrors α-synuclein’s neuroprotective-to-toxic transition and identifies a potential therapeutic niche for matrix-stabilizing agents like chondroitinase ABC^48^. However, broad ECM modulation risks disrupting neuroprotective interactions; our data advocate precision approaches targeting TH^+^ neuron-specific PNN or ECM may be a viable therapeutic strategy.

### Limitations and future directions

While our 3D model captures cell-ECM interactions absent in 2D cultures, particularly PNN formation and interstitial matrix stratification, the lack of microglia and vascular components likely attenuates ECM turnover rates observed in vivo. ECM remodelling in the brain is highly cell-type dependent. Neurons locally secrete ECM molecules and enzymes (e.g., reelin, brevican, MMPs) that remodel the perisynaptic space to support synaptic plasticity. Astrocytes, which are present in our model, are major producers of CSPGs, tenascins, and laminins, particularly under stress or injury. OPCs, also present in our model, respond to ECM cues that regulate their migration and maturation. In pathological contexts, inhibitory ECM components like aggrecan can hinder OPC differentiation. Both Mohamed et al.^14^ and Patikas et al.^15^ reported the presence of oligodendrocyte precursor cells (OPCs) in *SNCA* Trip and Isog midbrain organoid systems. OPCs express ECM receptors such as integrins and interact with fibronectin and CSPGs and thus may participate in the ECM phenotype observed here, particularly in the interstitial compartment. Therefore, the ECM accumulation observed in our model likely results from an imbalance in matrix production, degradation, and may be modified by the absence of microglial and vascular contributions.

By mapping the spatiotemporal progression of proteostatic and ECM changes in SNCA triplication organoids, we identify proteostasis-ECM axis dysregulation as an early driver of dopaminergic vulnerability. The preferential ECM remodelling in TH^+^ neurons, undetectable in 2D systems, highlights the necessity of 3D models to study matrix biology in neurodegeneration. These findings underscore the therapeutic potential of matrix-focused strategies in predegenerative PD stages, prioritizing interventions that preserve neuroprotective matrix interactions while inhibiting maladaptive remodelling.

## Acknowledgments

We wish to thank C-BIG at MNI, McGill University for assistance with iPSC cell lines, P. Haberkant and F. Stein for proteomics analysis at the EMBL proteomics facility (Heidelberg, Germany), Azenta/GeneWiz (Germany) for RNA-seq and Immagina SRL (Italy) for sequencing ribosome profiling libraries. We thank T. Kunath (University of Edinburgh, UK) for the *SNCA* lines.

## Author Contributions

ES and MZ contributed equally to this work. ES, MZ, SMJ, AD and CGG were responsible for conceptualization. ES, MZ, AD and CGG wrote the original draft of the article. CGG acquired funding. TD, AK, SMJ, SM and CGG were responsible for supervision. KC, GV, KSG and CP performed molecular/cellular experiments. All authors were responsible for investigation/methodology and reviewing and editing the article.

## Supplementary Material

**Supplementary Figure 1.**
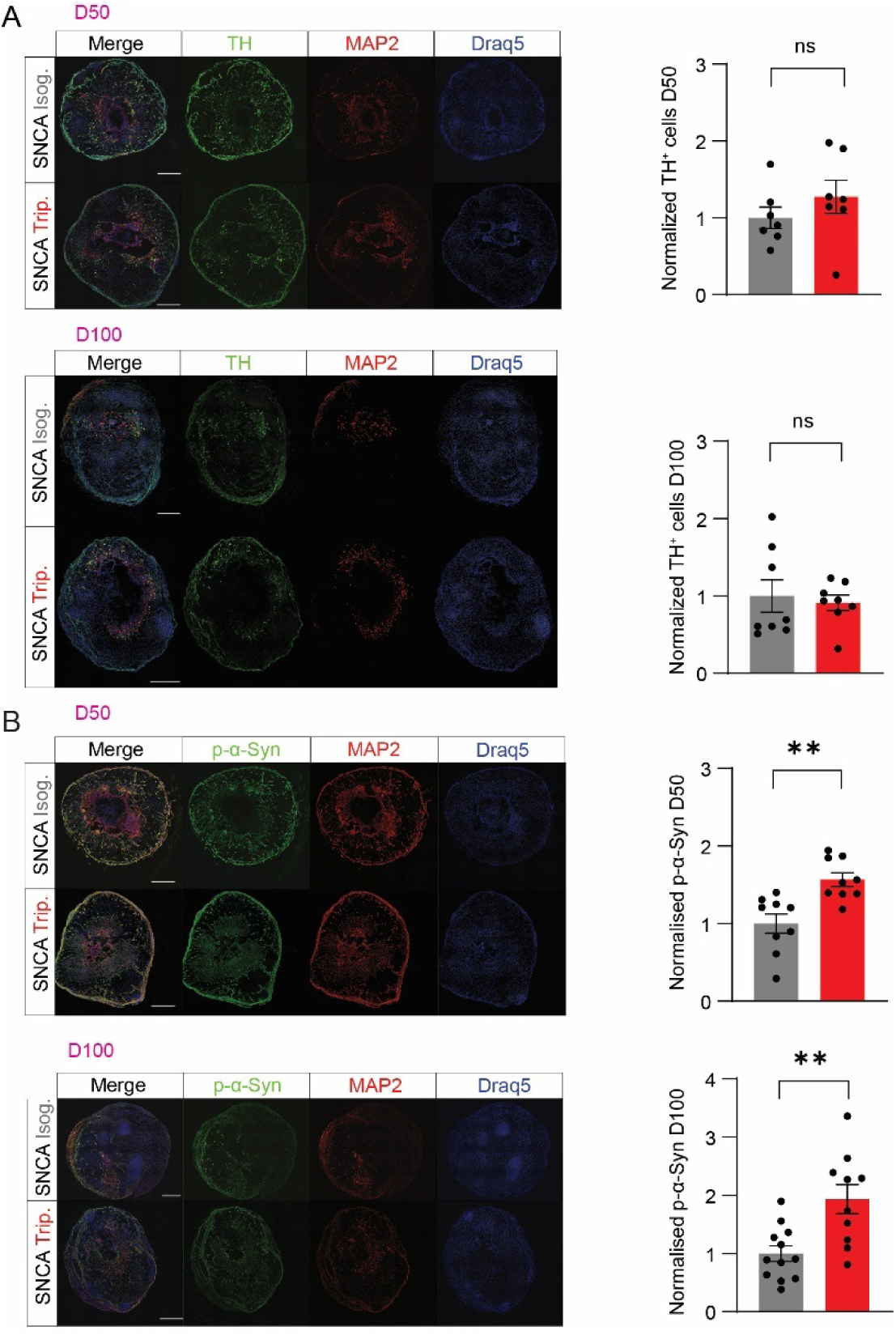
(A) Dopaminergic neuron quantification in *SNCA* Trip and Isog organoids. Representative immunofluorescence images showing tyrosine hydroxylase-positive (TH^+^, green) dopaminergic neurons, MAP2^+^ mature neurons (red), and Draq5^+^ nuclei (blue) in midbrain organoids at day 50 (D50) and day 100 (D100) of differentiation. Quantification reveals no significant difference in normalized TH^+^ cell numbers between *SNCA* Isog (grey bars) and *SNCA* Trip (red bars) conditions at either time point (D50: *P* >0.05, ns; D100: *P* >0.05, ns). Data represent mean ± SEM with individual data points shown. (B) Phosphorylated α-synuclein accumulation in *SNCA Trip* organoids. Representative immunofluorescence images displaying phosphorylated alpha-synuclein (p-α-Syn, green), MAP2^+^ neurons (red), and Draq5^+^ nuclei (blue) in organoids at D50 and D100. Quantitative analysis demonstrates significantly elevated normalized p-α-Syn levels in *SNCA* Trip organoids compared to isogenic controls at both D50 (\*\**P* <0.01) and D100 (\*\**P* <0.01). The progressive accumulation of phosphorylated alpha-synuclein in *SNCA* Trip conditions occurs without corresponding loss of dopaminergic neurons, indicating early pathological protein aggregation preceding neurodegeneration. Scale bars = 500 μm. Data shown are mean ± SEM with individual; Student’s *t*-test with parametric or non-parametric (*Mann-Whitney*) test; \*\**P* < 0.01, ns = not significant.

**Supplementary Figure 2.** Raw data from immunoblots.

**Supplementary Figure 3.**
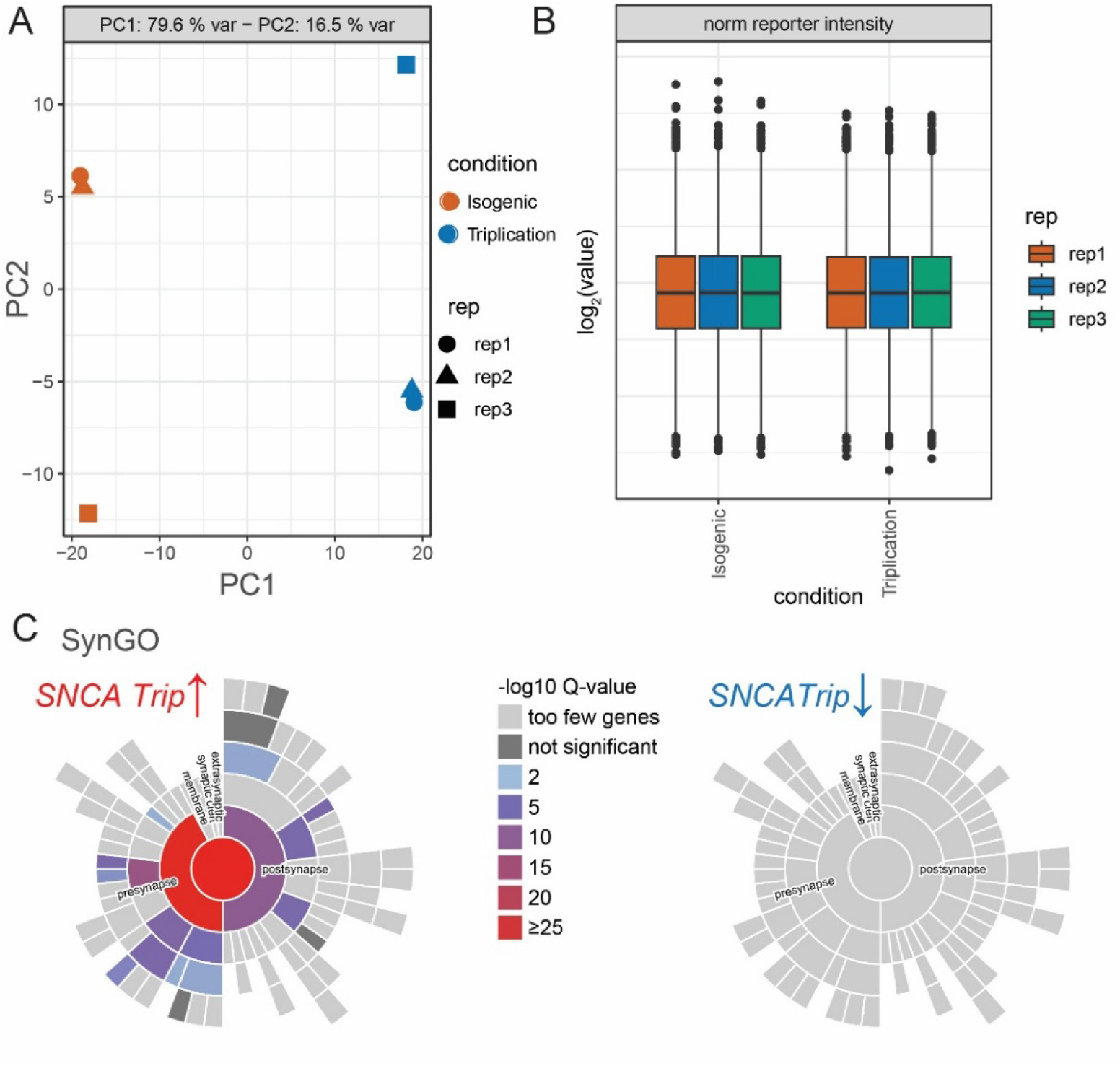
(A) Principal component analysis (PCA) of TMT proteomics data showing clear separation between *SNCA* Trip (blue) and *SNCA* Isog (orange) organoids at D100. PC1 explains 79.6% of variance and PC2 explains 16.5% of variance. (B) Normalized reporter ion intensity distributions across biological replicates (rep1-3) for both conditions, demonstrating consistent sample preparation and data quality. (C) SynGO analysis of significantly altered proteins in *SNCA* Trip organoids. Left panel shows upregulated proteins (*SNCA* Trip↑) with enrichment in postsynaptic and presynaptic compartments. Right panel shows downregulated proteins (*SNCA* Trip↓) with minimal synaptic enrichment. Colour intensity represents -log_10_ Q-value significance, with red indicating highest enrichment (≥25) and grey indicating non-significant or insufficient gene representation.

**Supplementary Figure 4.**
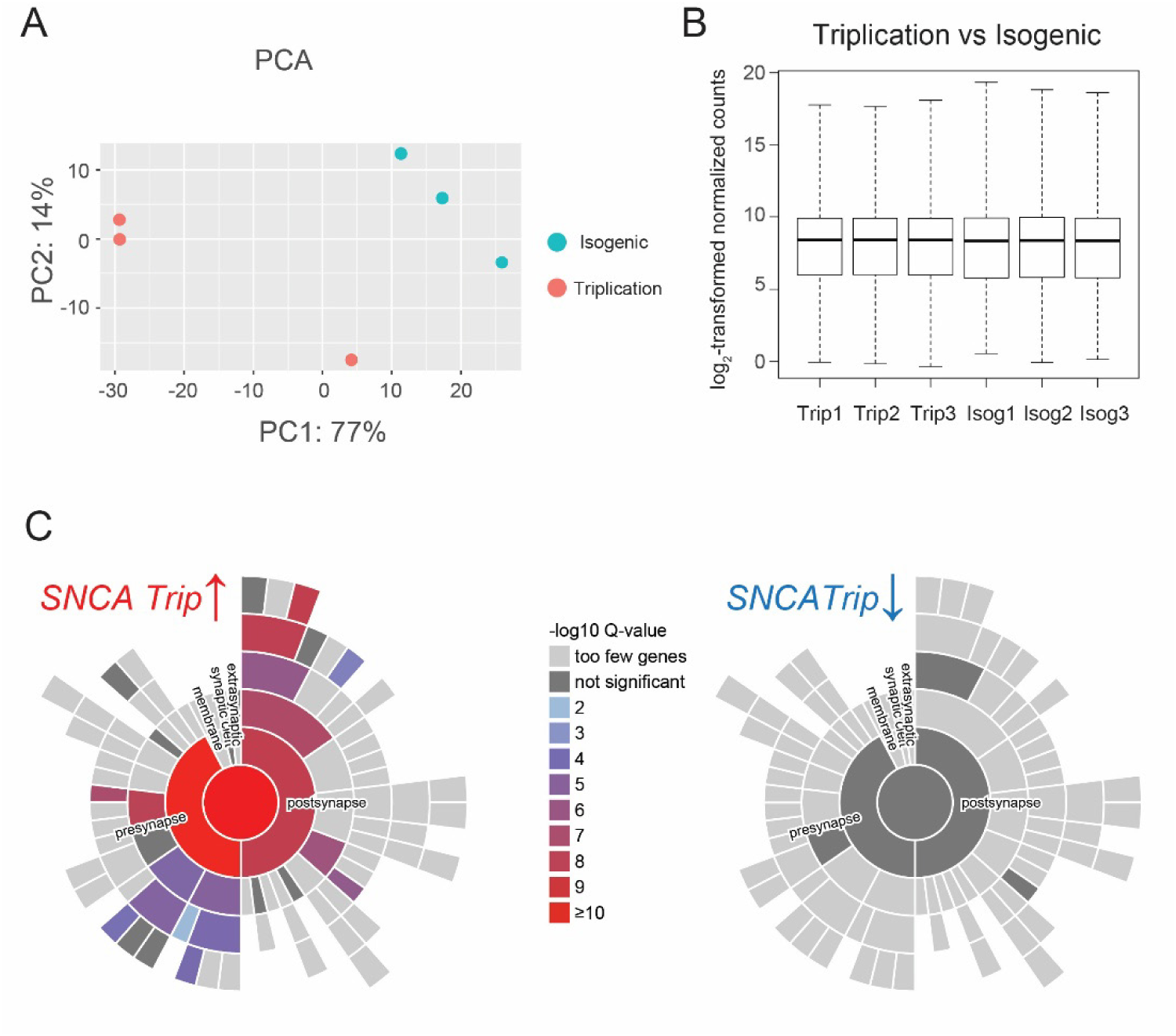
(A) Principal component analysis (PCA) of bulk RNA-seq data from D100 *SNCA* Trip (red circles) and SNCA Isog (teal circles) midbrain organoids. PC1 explains 77% of variance and PC2 explains 14% of variance, demonstrating robust transcriptional separation between genotypes across three biological replicates per condition. (B) Normalized count distributions (log₂-transformed) showing consistent data quality across all samples. Box plots display the distribution of gene expression values for each biological replicate (Trip1-3, Isog1-3), with median expression levels ranging from 6-8 log₂ counts. The uniform distributions confirm successful normalization using DESeq2’s variance stabilizing transformation. (C) SynGO analysis of significantly differentially expressed genes (*Padj* ≤ 10⁻⁴, |log₂FC| ≥ 2). Left panel shows upregulated genes in *SNCA* Trip organoids (*SNCA* Trip↑) with strong enrichment in synaptic compartments, particularly postsynaptic and presynaptic regions (red indicating highest significance, -log₁₀ Q-value ≥10). Right panel shows downregulated genes (*SNCA* Trip↓) with minimal synaptic annotation, predominantly appearing as grey sectors indicating “too few genes” or “not significant” categories. Colour intensity represents statistical significance of synaptic gene ontology enrichment according to the SynGO database.

**Supplementary Figure 5.**
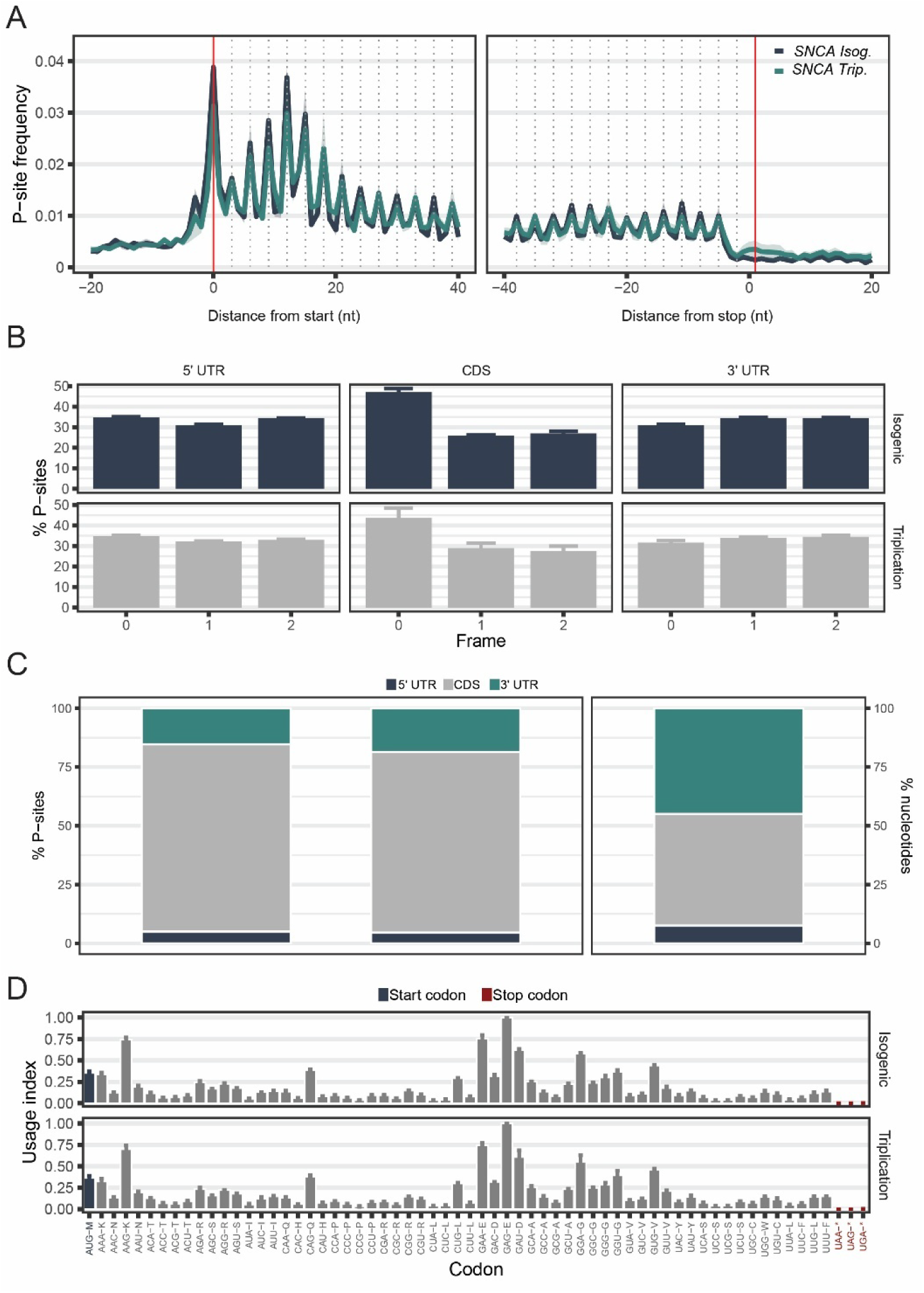
(A) Metaprofiles showing P-site frequency distribution relative to start codons (left panel) and stop codons (right panel) in D100 *SNCA* Trip (teal) and *SNCA* Isog (dark blue) midbrain organoids. Clear 3-nucleotide periodicity around start codons confirms proper ribosome positioning and high-quality ribosome profiling data. Red vertical lines indicate start codon (AUG) and stop codon positions. (B) Reading frame distribution of P-sites across transcript regions showing percentage of ribosome-protected fragments (RPFs) in frames 0, 1, and 2 within 5’ UTR, coding sequence (CDS), and 3’ UTR regions. High enrichment in frame 0 within the CDS (>85%) confirms accurate P-site mapping and translation fidelity in both conditions. (C) Distribution of P-sites across transcript regions showing the percentage of RPFs mapping to 5’ UTR (dark blue), CDS (grey), and 3’ UTR (teal) regions. The predominant enrichment in CDS regions indicates successful capture of actively translating ribosomes with minimal background from non-coding regions. (D) Codon usage index analysis showing the relative usage of start codons (dark blue) and stop codons (red) across all detected codons in *SNCA* Isog (top) and *SNCA* Trip (bottom) organoids. Similar codon usage patterns between conditions confirm consistent ribosome profiling quality and absence of systematic biases. All analyses were performed using riboWaltz with standard quality control parameters, confirming high data integrity across both genotypes.

**Sup. Table 1 Limma results, proteomics**

**Sup. Table 2 Deseq2 results, RNA-seq**

**Sup. Table 3 XTAIL results, RPF**

**Sup. Table 4 Antibodies Table**

**Sup. Table 5 Statistics details**

