## Supplementary Figure 2 for "*SNCA* triplication disrupts proteostasis and extracellular architecture prior to neurodegeneration in human midbrain organoids"

### RAW DATA WESTERN BLOTS

D50

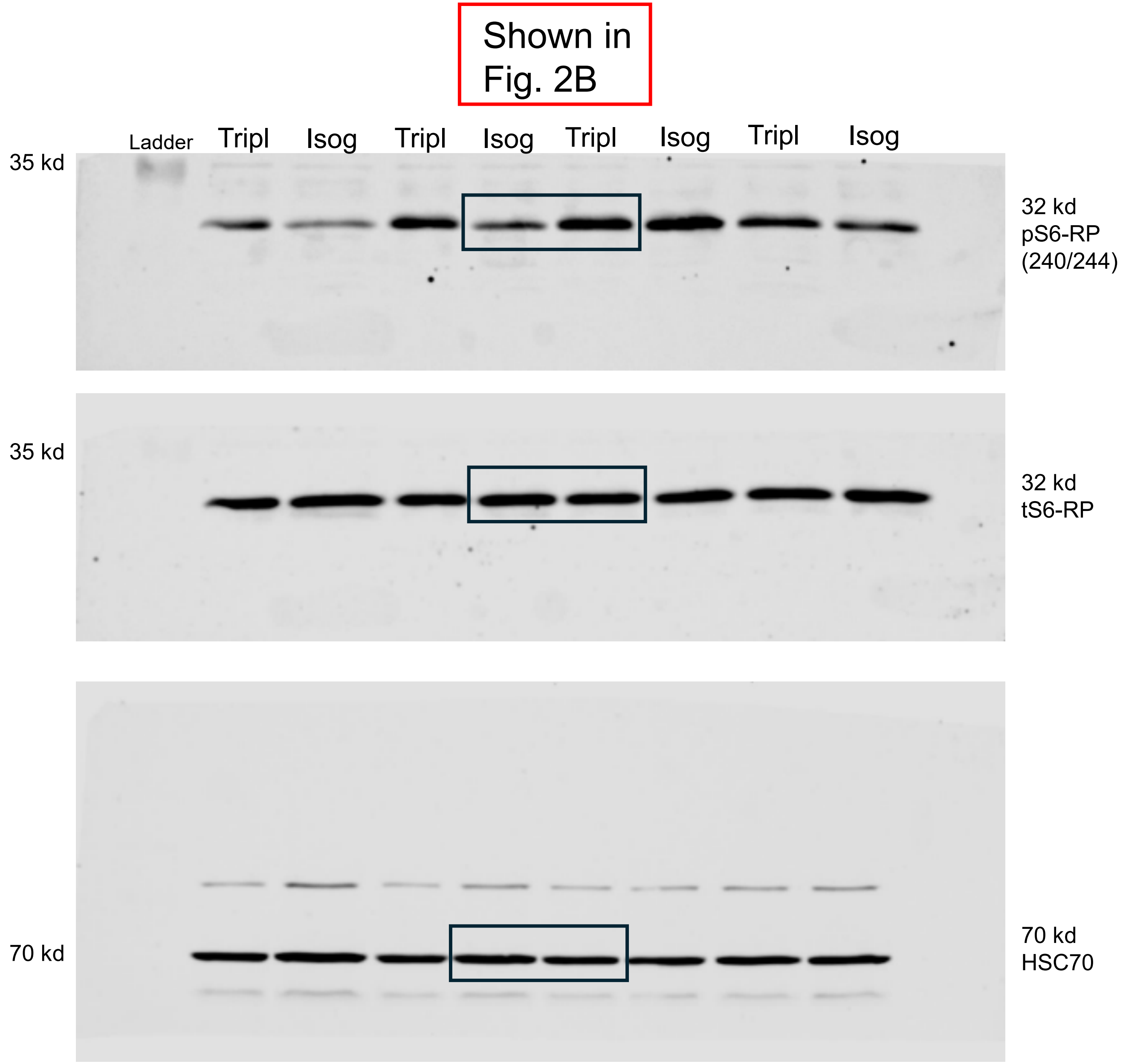

Shown in Fig. 2B
Ladder
Tripl
Isog
Tripl
Isog
Tripl
Isog
Tripl
Isog
35 kd
32 kd
pS6-RP
(240/244)
35 kd
32 kd
tS6-RP
70 kd
70 kd
HSC70

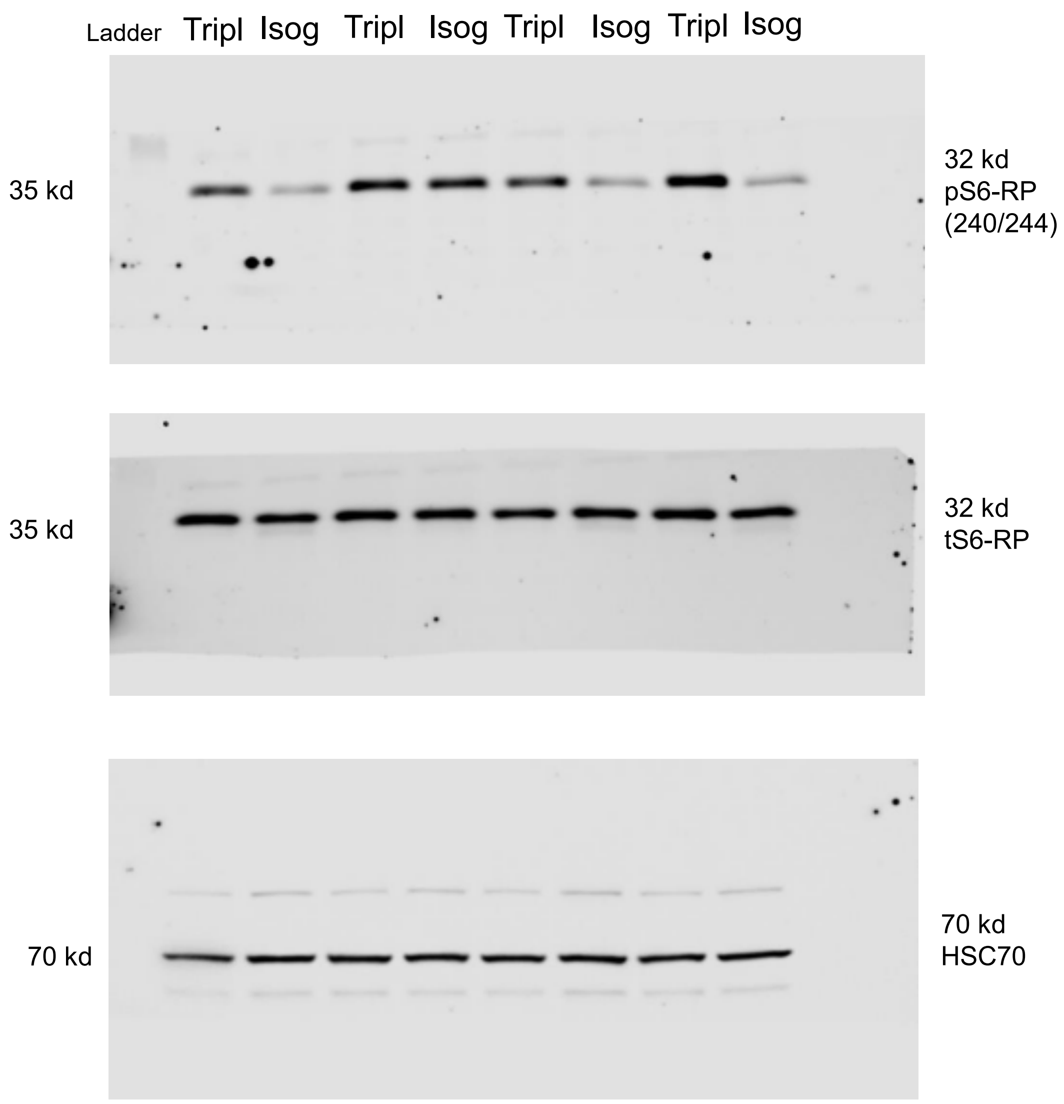
Ladder
Tripl
Isog
Tripl
Isog
Tripl
Isog
Tripl
Isog
35 kd
32 kd
pS6-RP
(240/244)
35 kd
32 kd
tS6-RP
70 kd
70 kd
HSC70

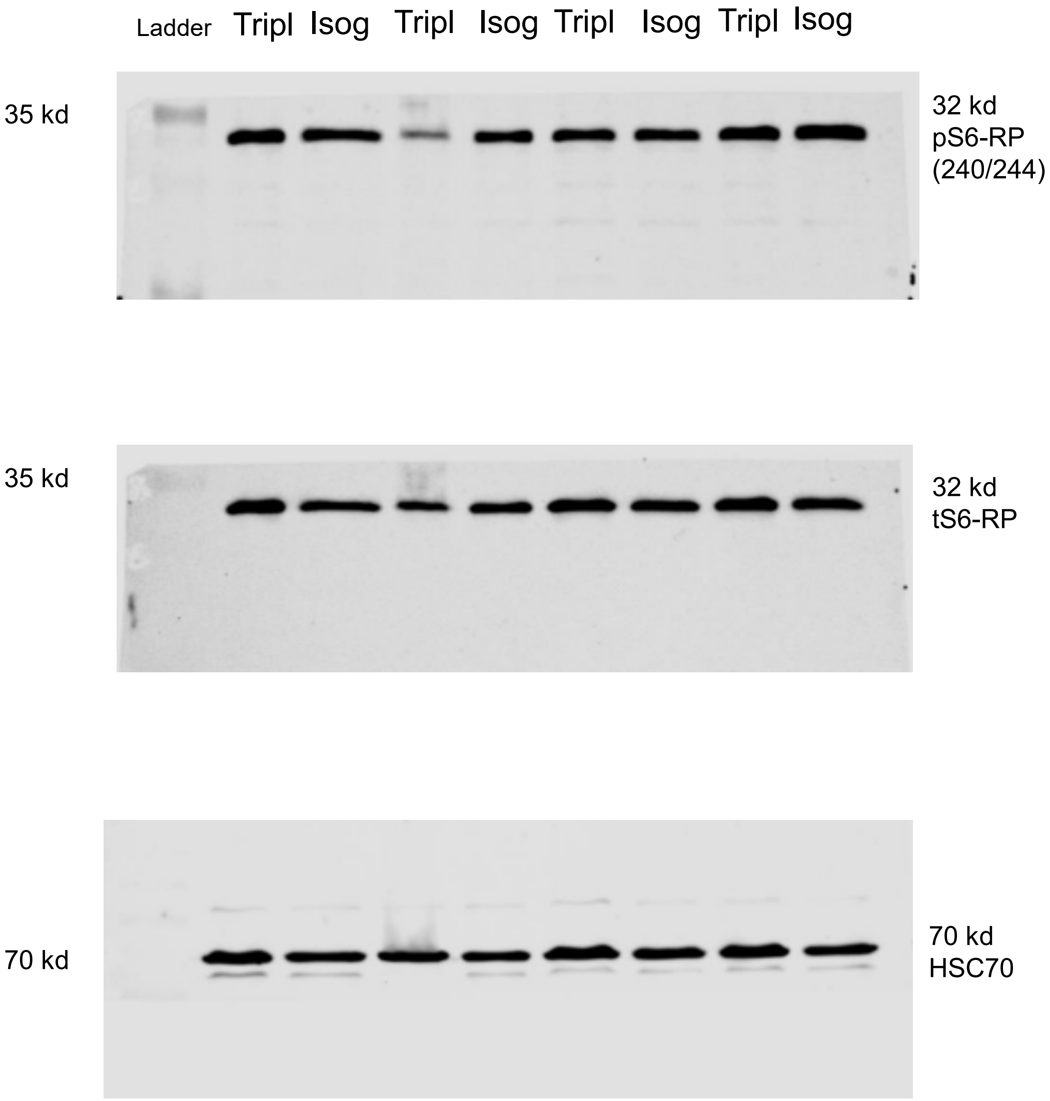

Ladder
Tripl
Isog
Tripl
Isog
Tripl
Isog
Tripl
Isog
35 kd
32 kd
pS6-RP
(240/244)
35 kd
32 kd
tS6-RP
70 kd
70 kd
HSC70

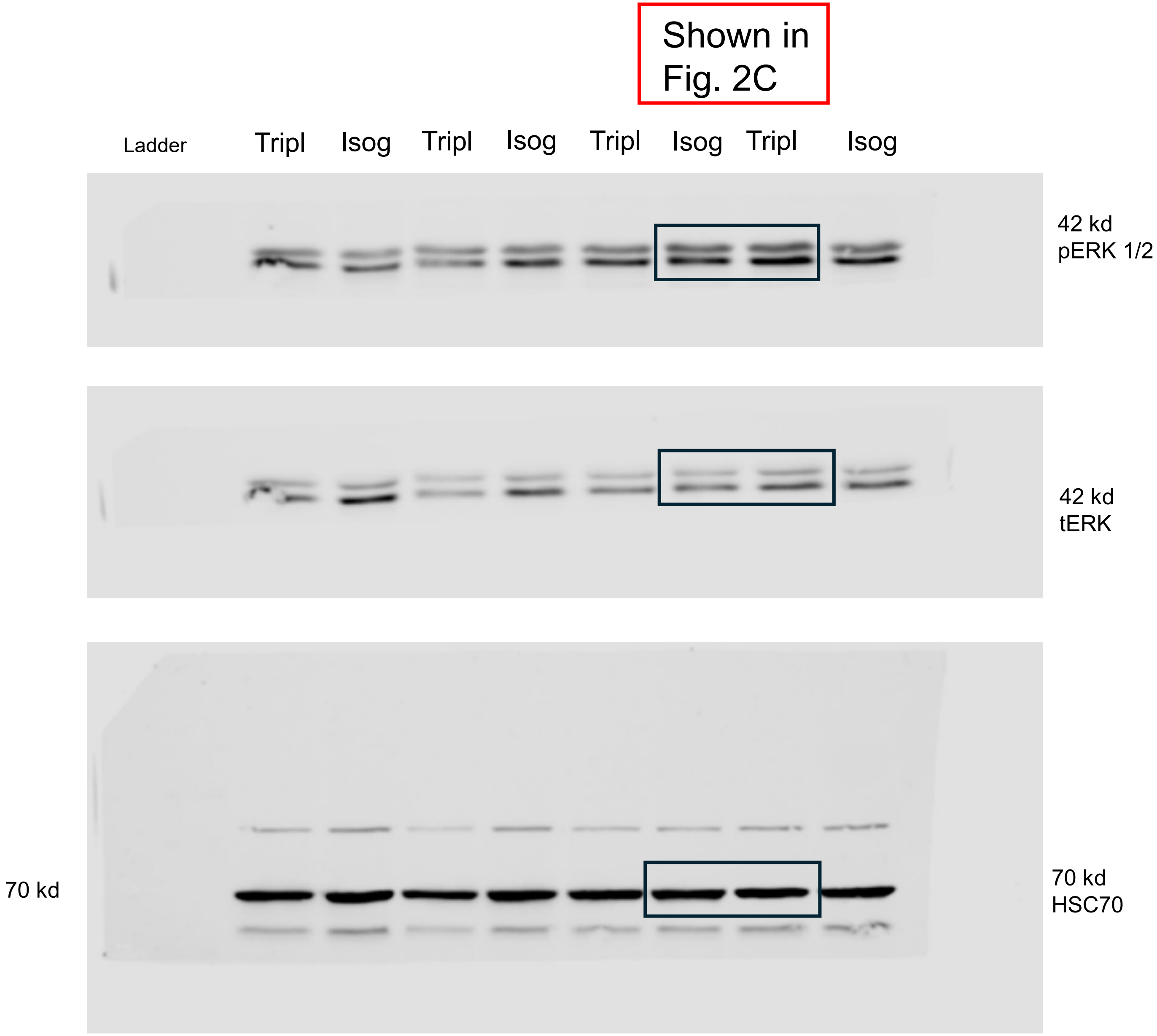

Shown in Fig. 2C
Ladder
Tripl
Isog
Tripl
Isog
Tripl
Isog
Tripl
Isog
42 kd
pERK 1/2
42 kd
tERK
70 kd
70 kd
HSC70

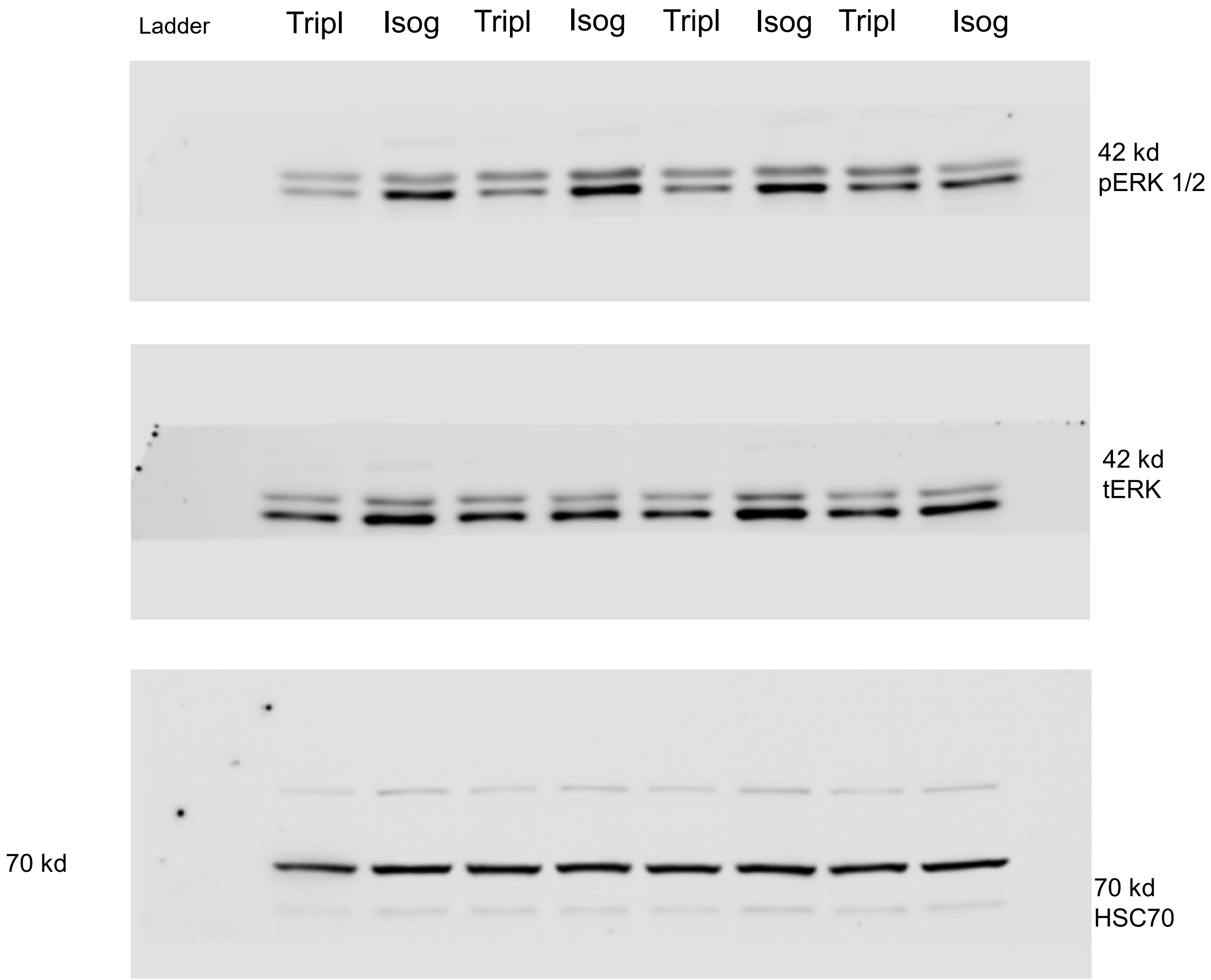
Ladder
Tripl
Isog
Tripl
Isog
Tripl
Isog
Tripl
Isog
42 kd
pERK 1/2
42 kd
tERK
70 kd
70 kd
HSC70

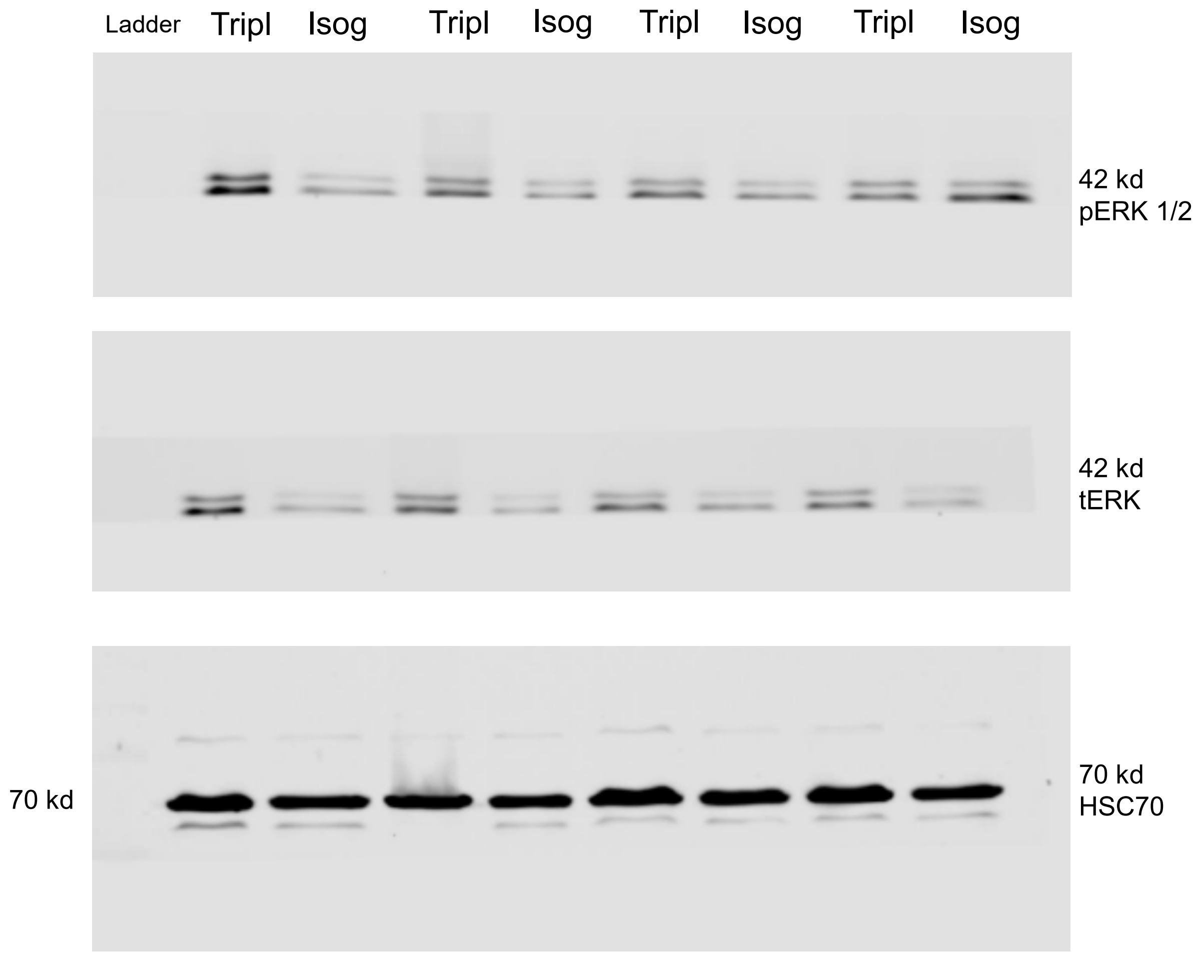

Ladder
Tripl
Isog
Tripl
Isog
Tripl
Isog
Tripl
Isog
42 kd
pERK 1/2
42 kd
tERK
70 kd
70 kd
HSC70

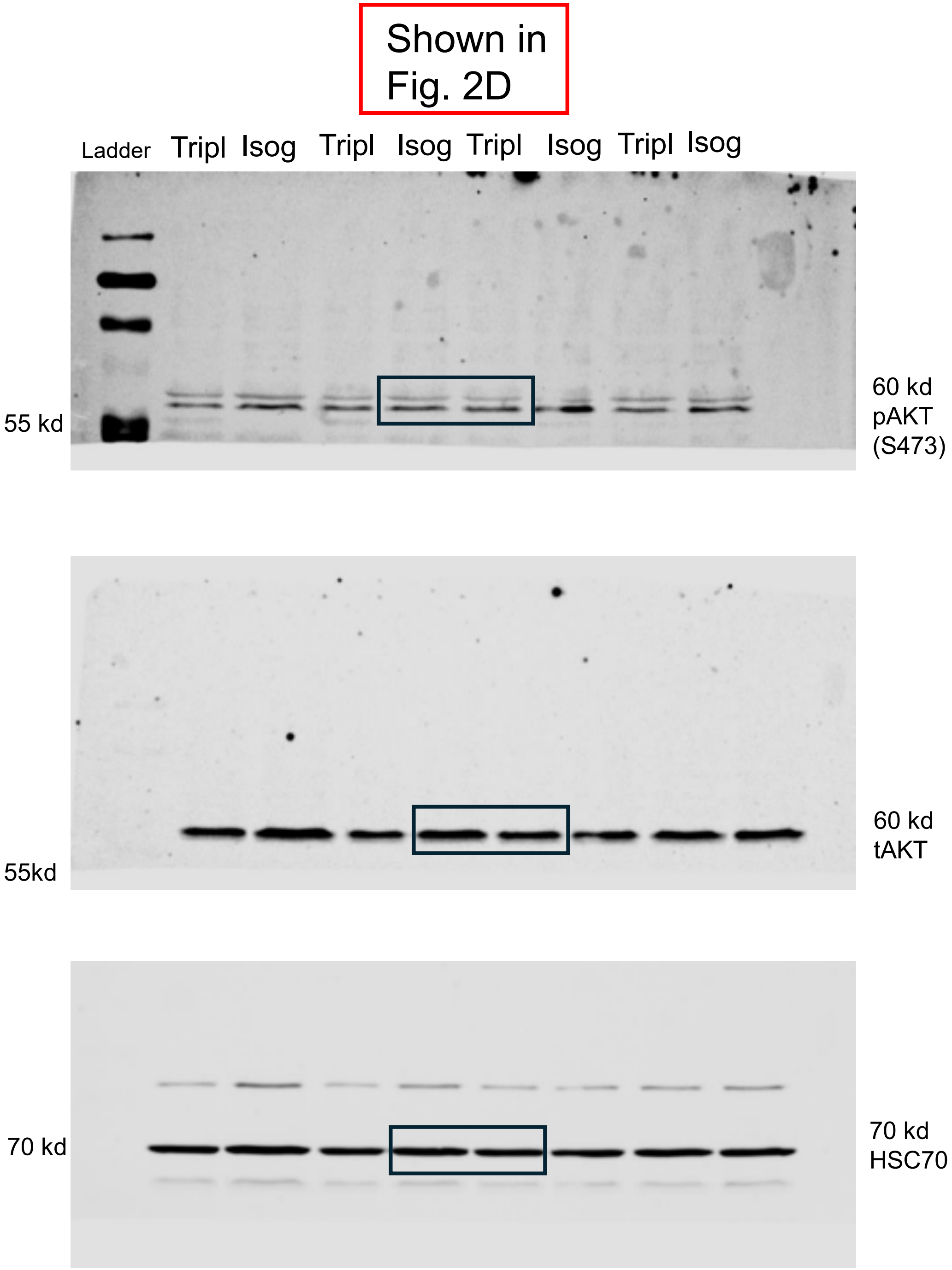

Shown in Fig. 2D
Ladder
Tripl
Isog
Tripl
Isog
Tripl
Isog
Tripl
Isog
55 kd
60 kd pAKT (S473)
55kd
60 kd tAKT
70 kd
70 kd HSC70

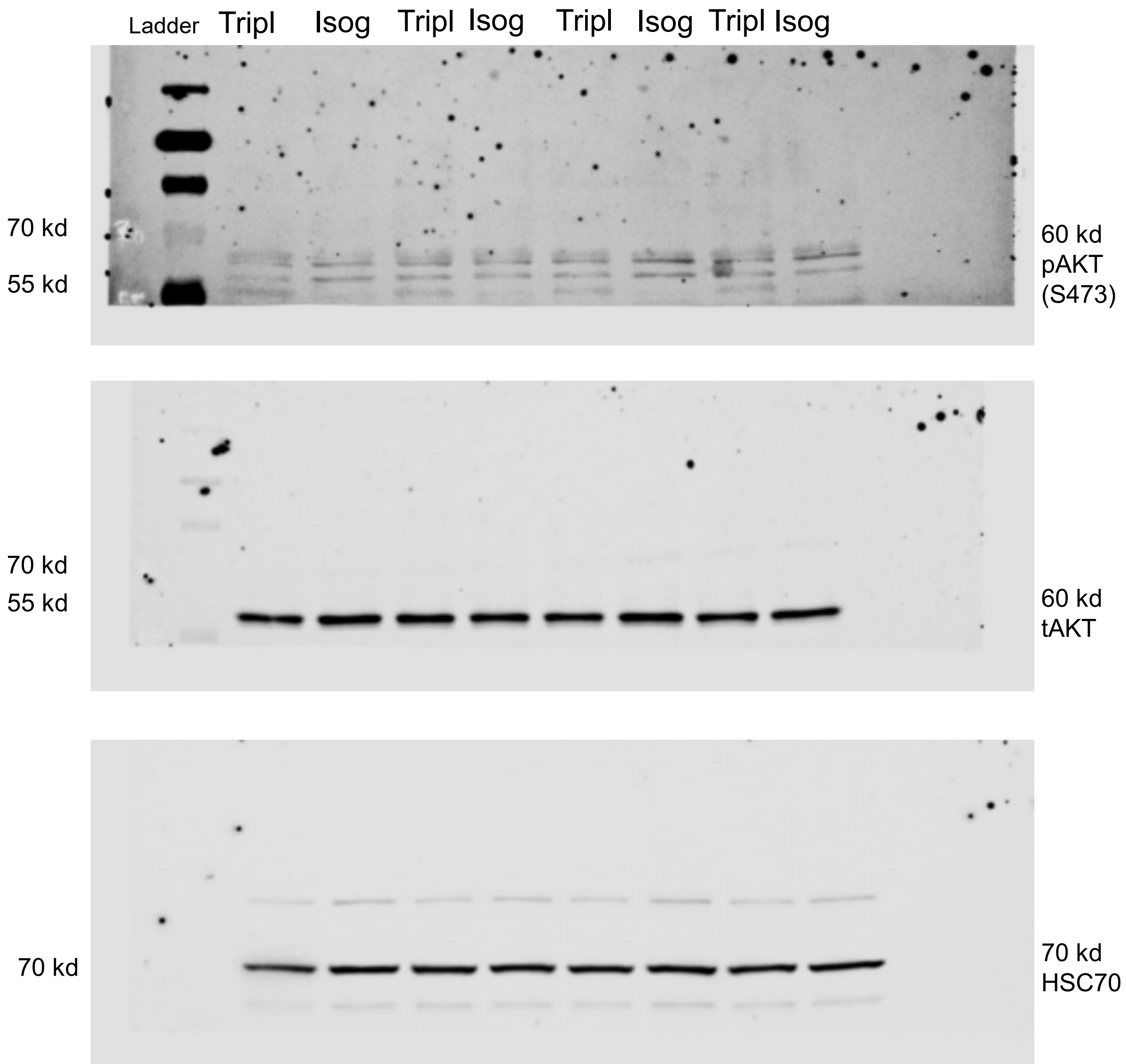
Ladder
Tripl
Isog
Tripl
Isog
Tripl
Isog
Tripl
Isog
70 kd
55 kd
60 kd
pAKT
(S473)
70 kd
55 kd
60 kd
tAKT
70 kd
70 kd
HSC70

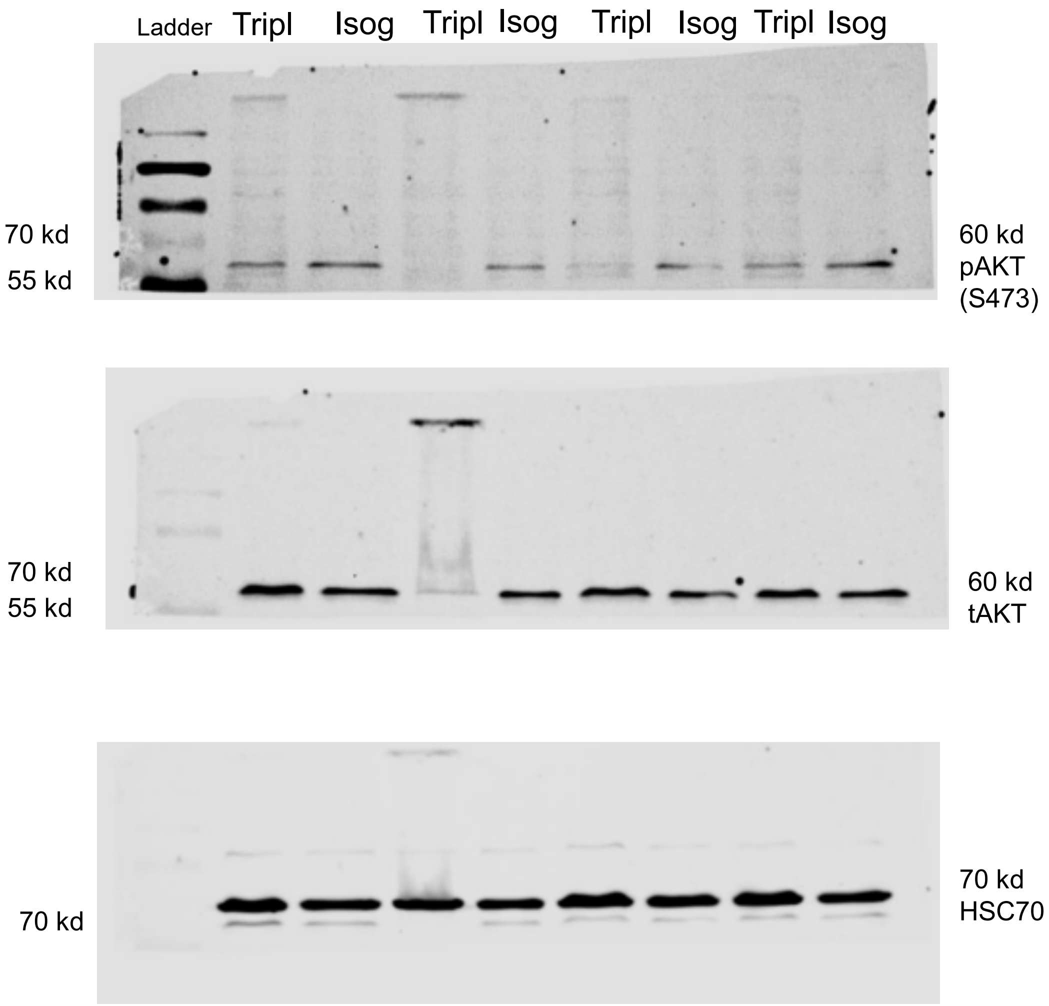
Ladder
Tripl
Isog
Tripl
Isog
Tripl
Isog
Tripl
Isog
70 kd
55 kd
60 kd
pAKT
(S473)
70 kd
55 kd
60 kd
tAKT
70 kd
70 kd
HSC70

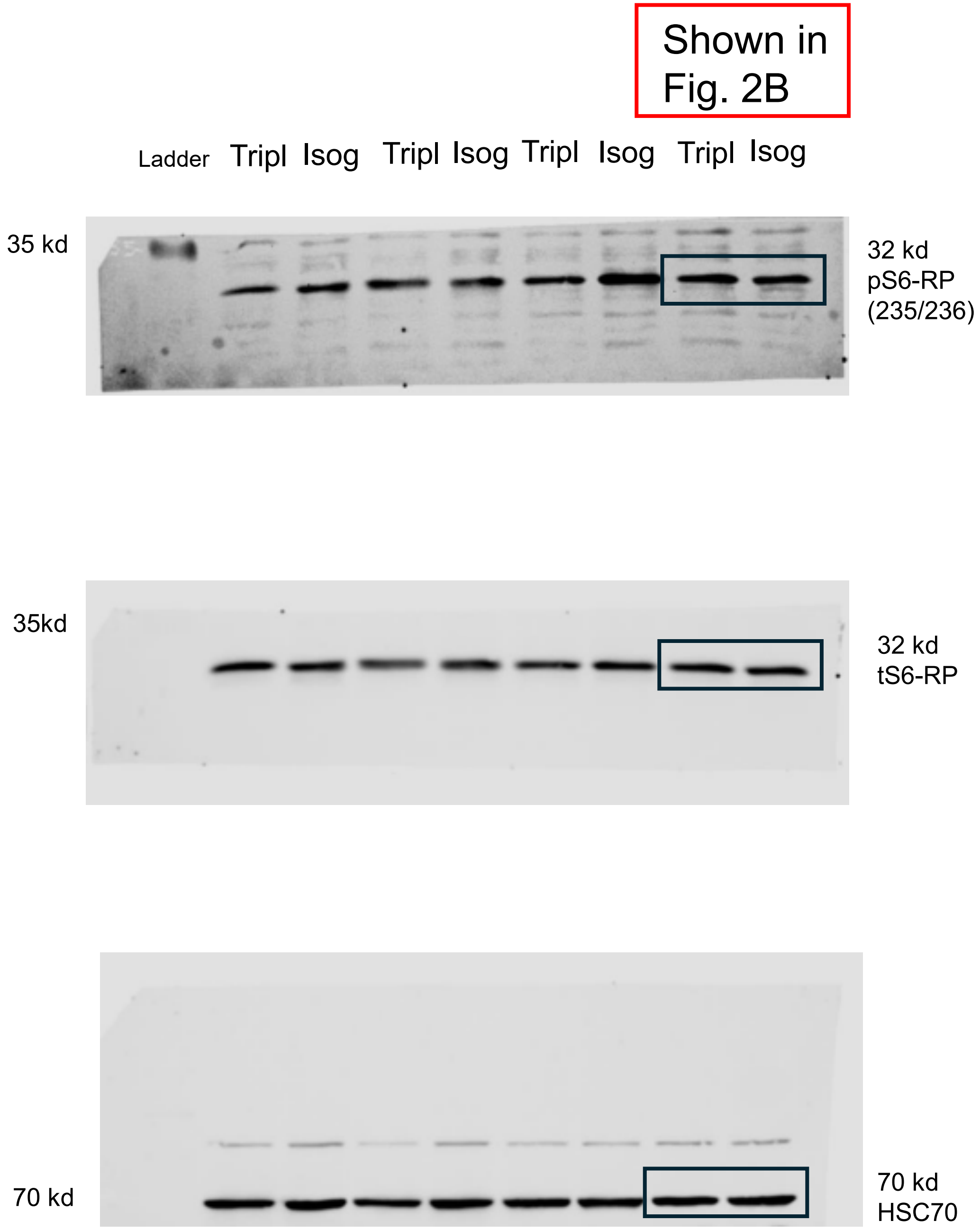

Shown in Fig. 2B
Ladder
Tripl
Isog
Tripl
Isog
Tripl
Isog
Tripl
Isog
35 kd
32 kd
pS6-RP
(235/236)
35kd
32 kd
tS6-RP
70 kd
70 kd
HSC70

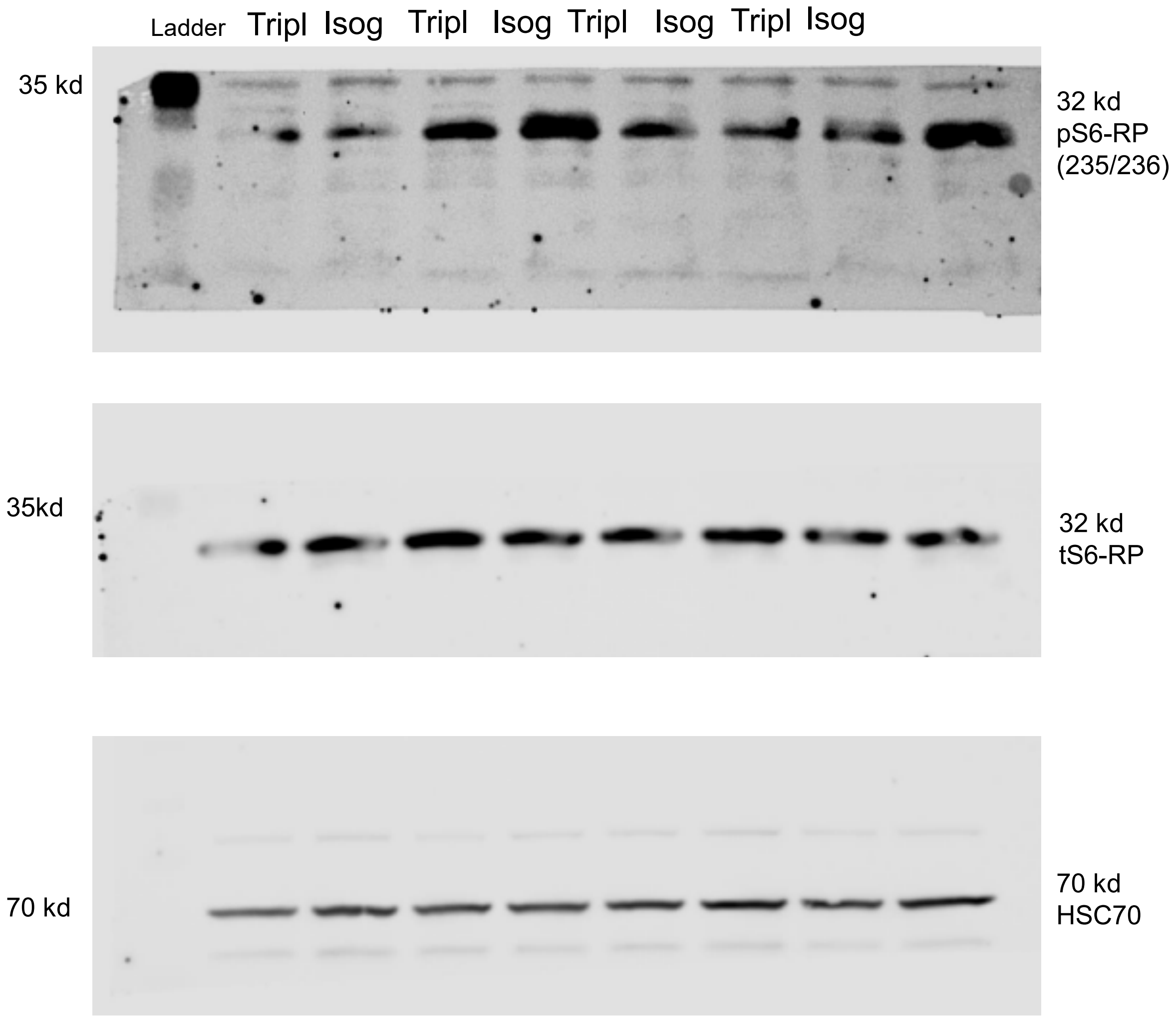
Ladder
Tripl
Isog
Tripl
Isog
Tripl
Isog
Tripl
Isog
35 kd
32 kd
pS6-RP
(235/236)
35kd
32 kd
tS6-RP
70 kd
70 kd
HSC70

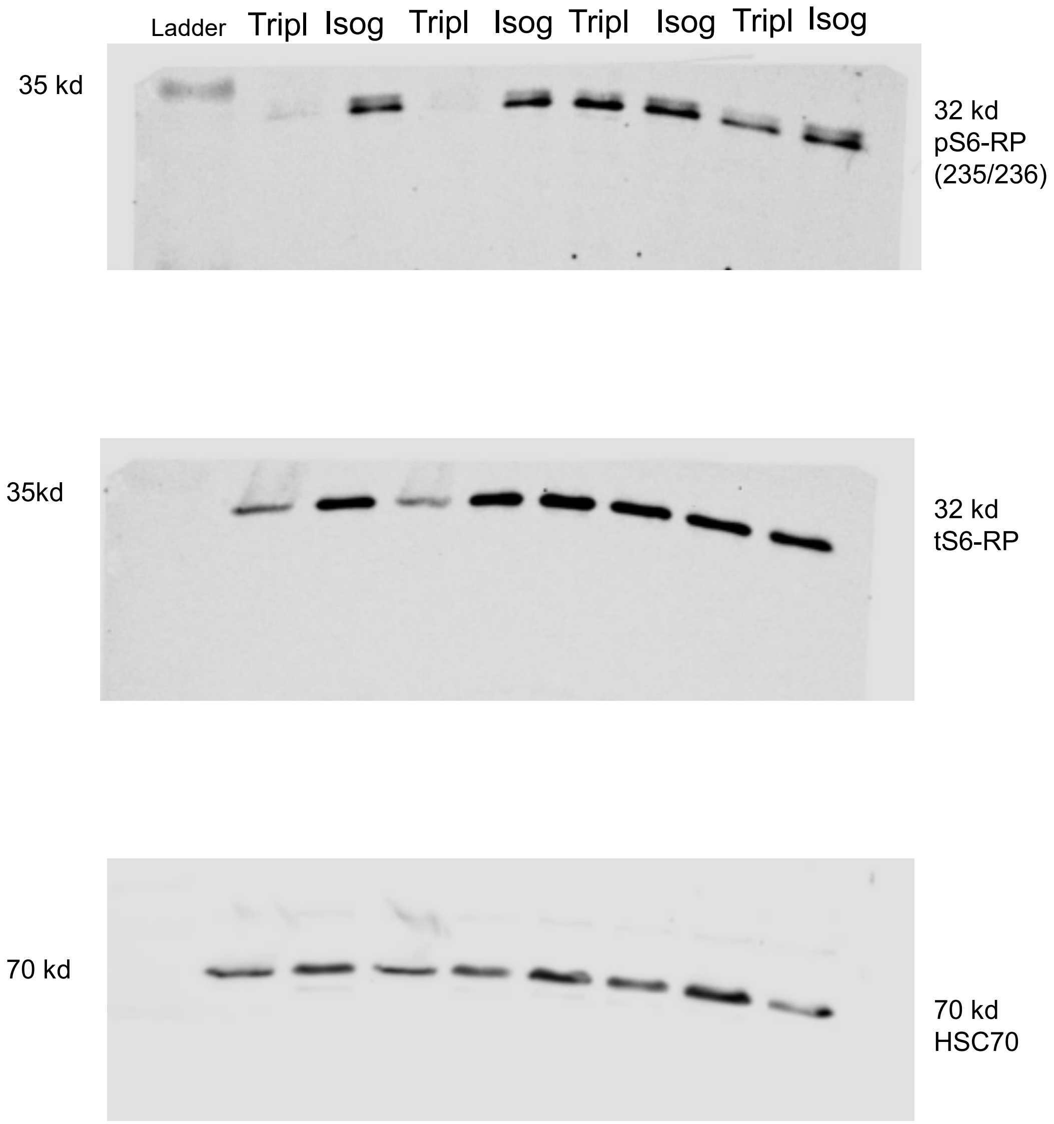

Ladder
Tripl
Isog
Tripl
Isog
Tripl
Isog
Tripl
Isog
35 kd
32 kd
pS6-RP
(235/236)
35kd
32 kd
tS6-RP
70 kd
70 kd
HSC70

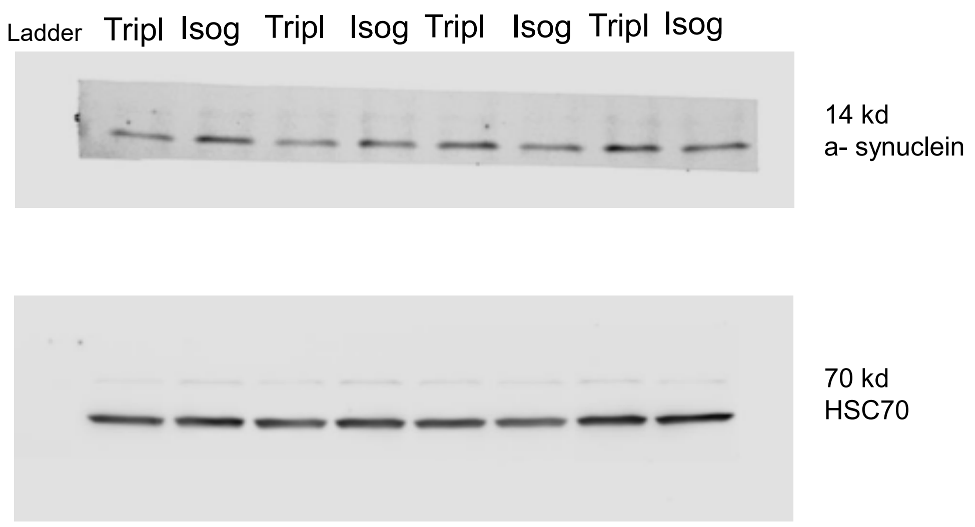

Ladder
Tripl
Isog
Tripl
Isog
Tripl
Isog
Tripl
Isog
14 kd
a- synuclein
70 kd
HSC70

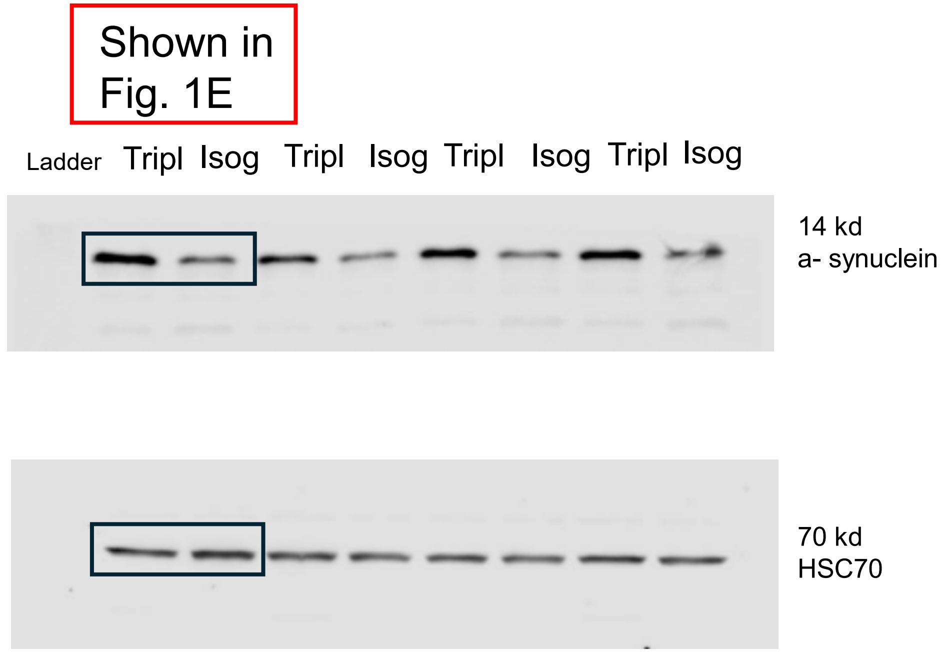

Shown in Fig. 1E
Ladder
Tripl
Isog
Tripl
Isog
Tripl
Isog
Tripl
Isog
14 kd
a- synuclein
70 kd
HSC70

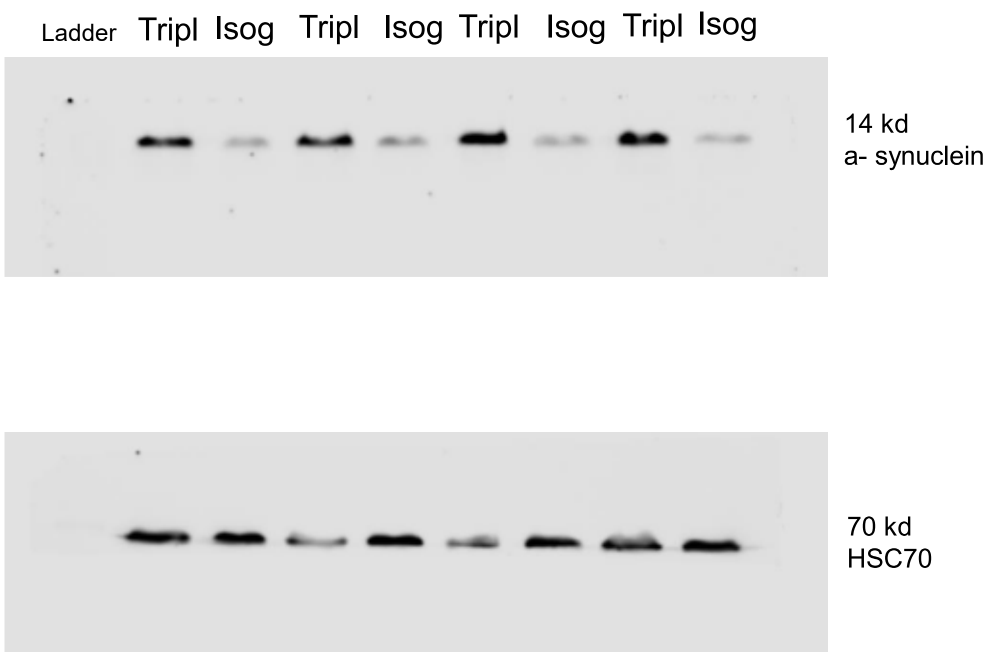
Ladder Tripl Isog Tripl Isog Tripl Isog Tripl Isog
14 kd
a- synuclein
70 kd
HSC70

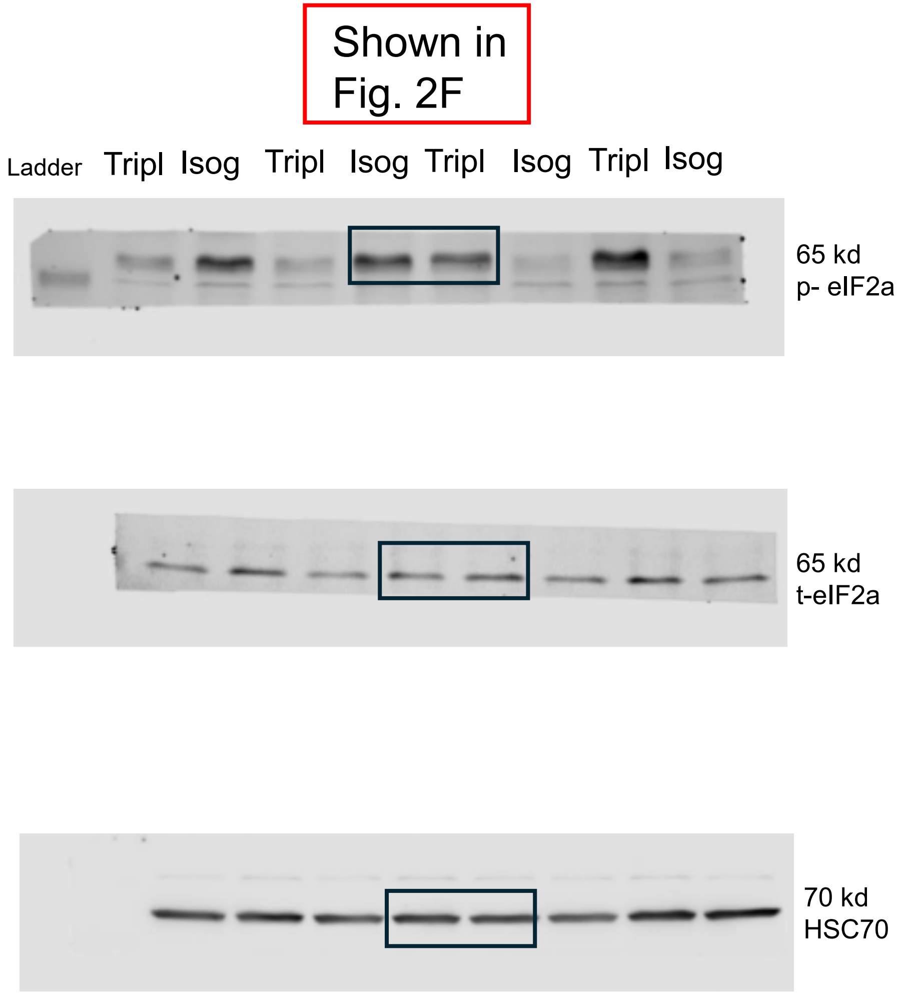
Shown in Fig. 2F
Ladder
Tripl
Isog
Tripl
Isog
Tripl
Isog
Tripl
Isog
65 kd
p- eIF2a
65 kd
t-eIF2a
70 kd
HSC70

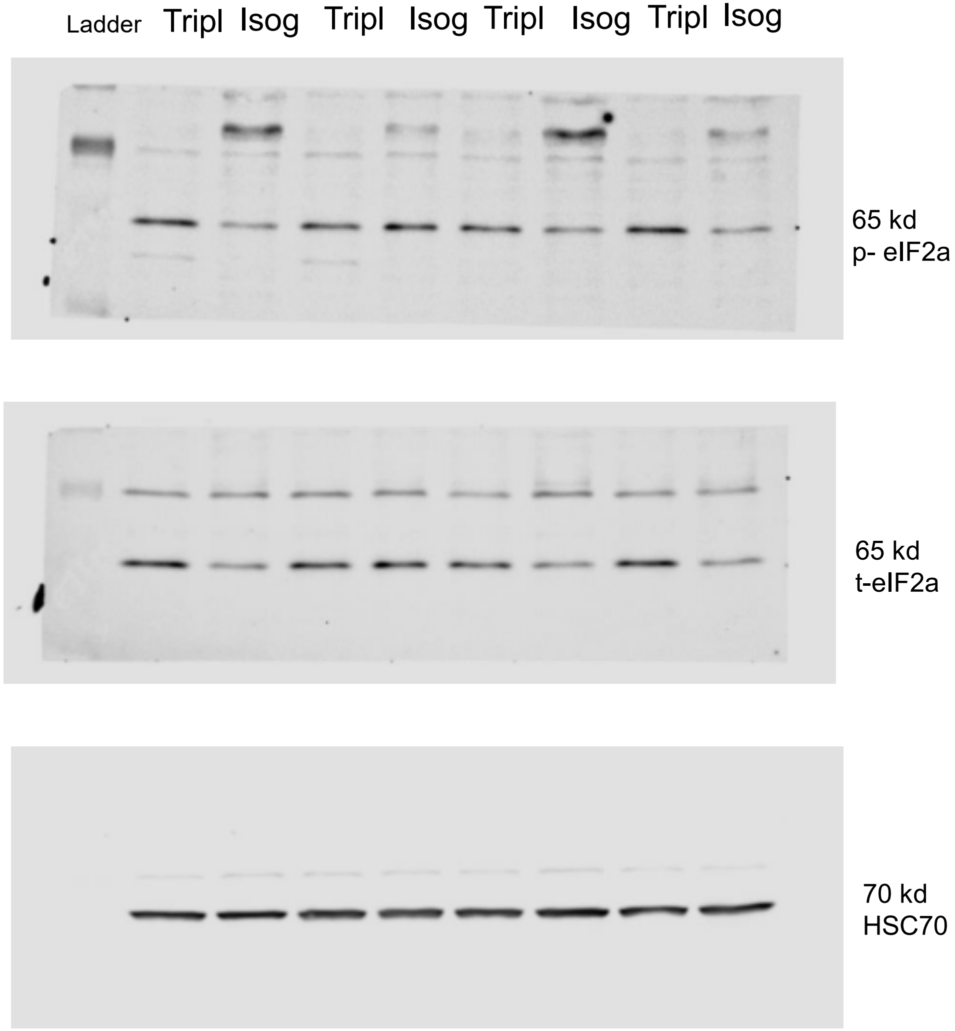
Ladder
Tripl
Isog
Tripl
Isog
Tripl
Isog
Tripl
Isog
65 kd
p- eIF2a
65 kd
t-eIF2a
70 kd
HSC70

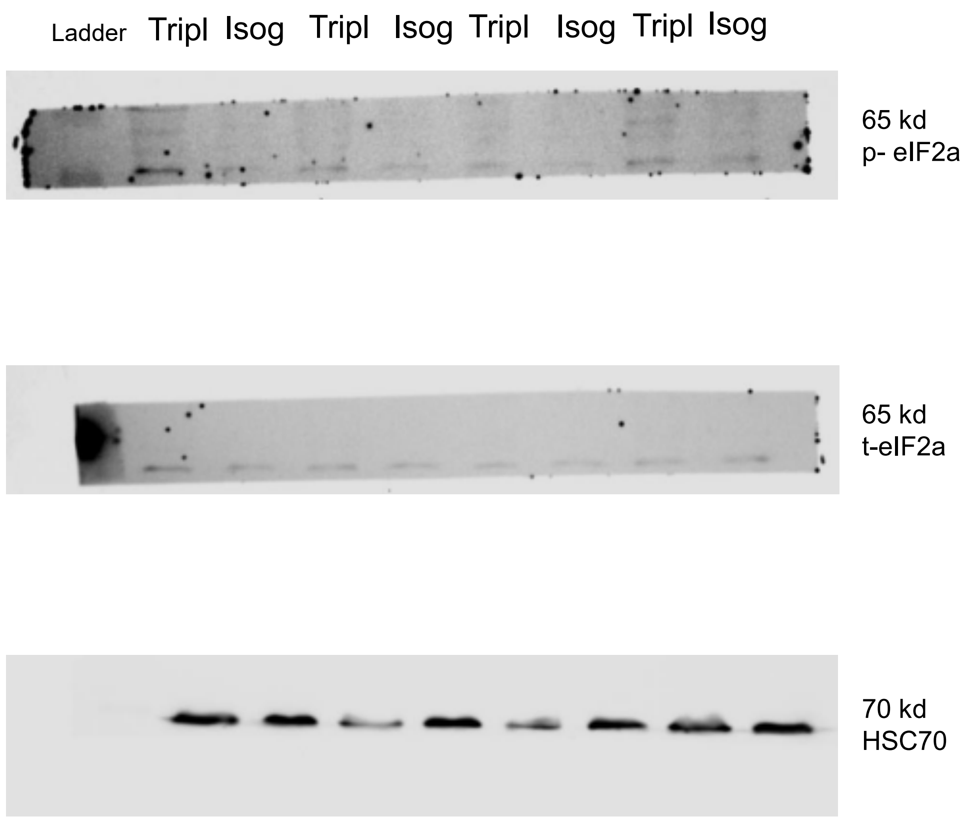
Ladder
Tripl
Isog
Tripl
Isog
Tripl
Isog
Tripl
Isog
65 kd
p- eIF2a
65 kd
t-eIF2a
70 kd
HSC70

### RAW DATA WESTERN BLOTS

D100

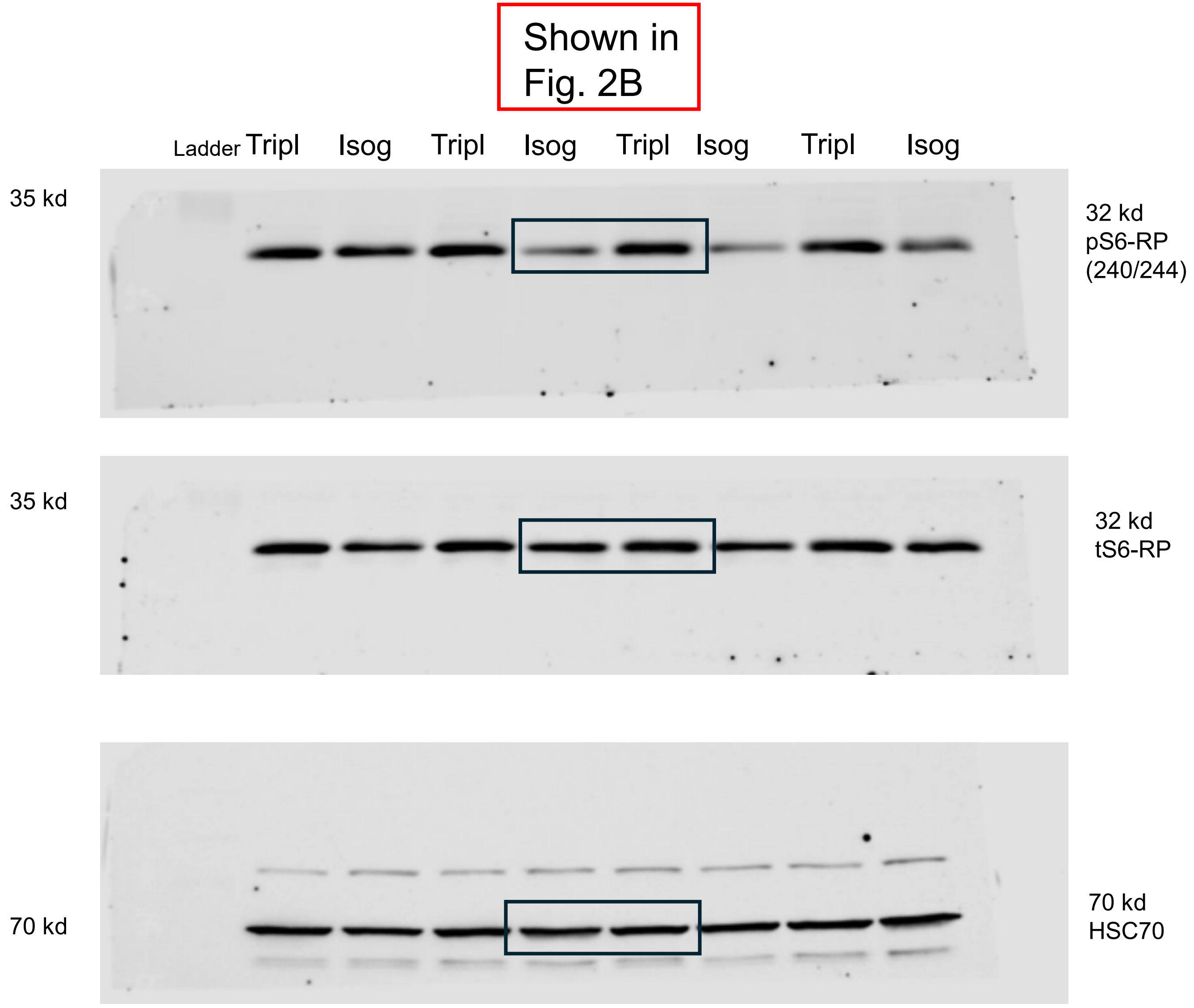
Shown in Fig. 2B
Ladder
Tripl
Isog
Tripl
Isog
Tripl
Isog
Tripl
Isog
35 kd
32 kd pS6-RP (240/244)
35 kd
32 kd tS6-RP
70 kd
70 kd HSC70

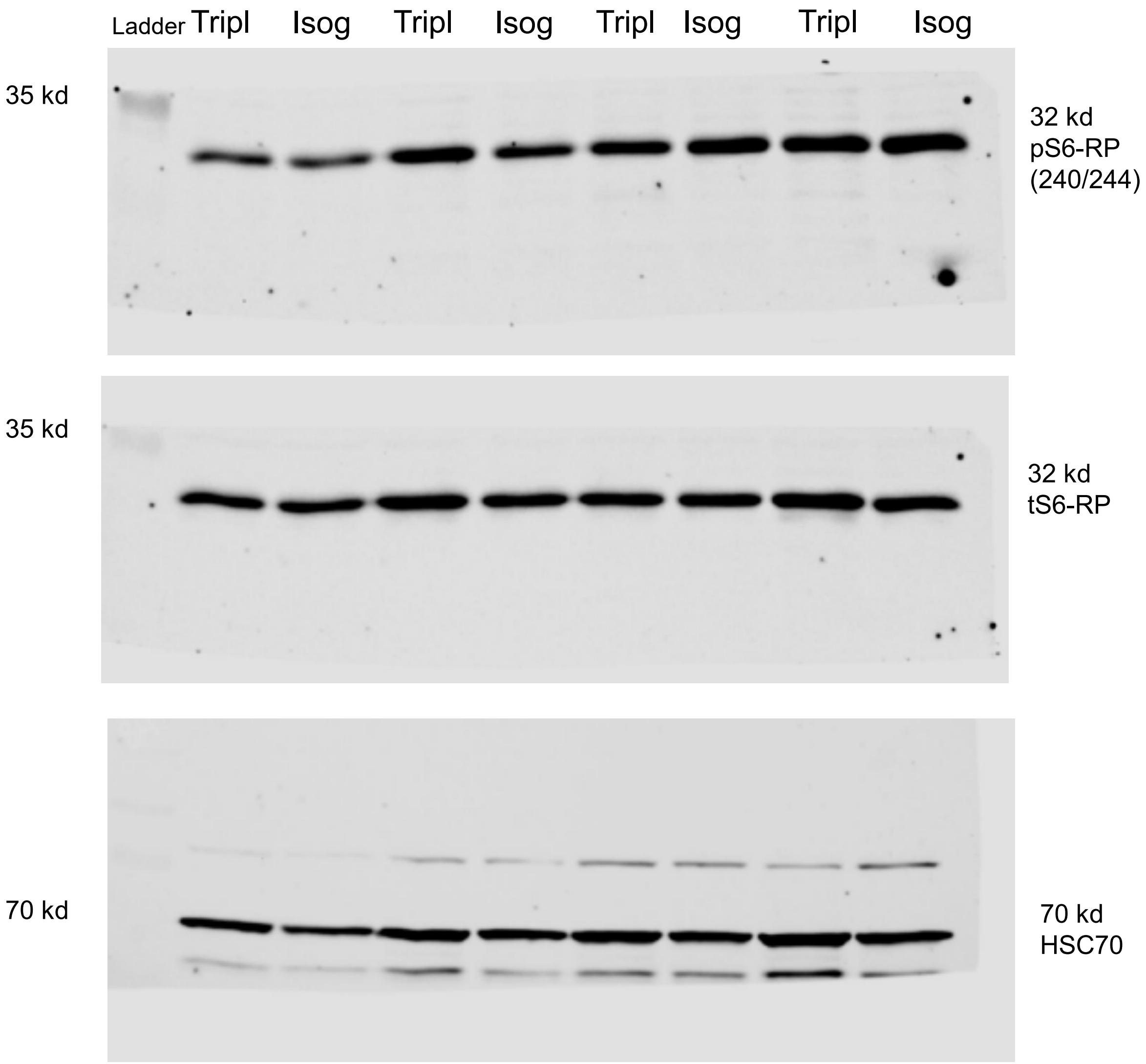
Ladder
Tripl
Isog
Tripl
Isog
Tripl
Isog
Tripl
Isog
35 kd
32 kd
pS6-RP
(240/244)
35 kd
32 kd
tS6-RP
70 kd
70 kd
HSC70

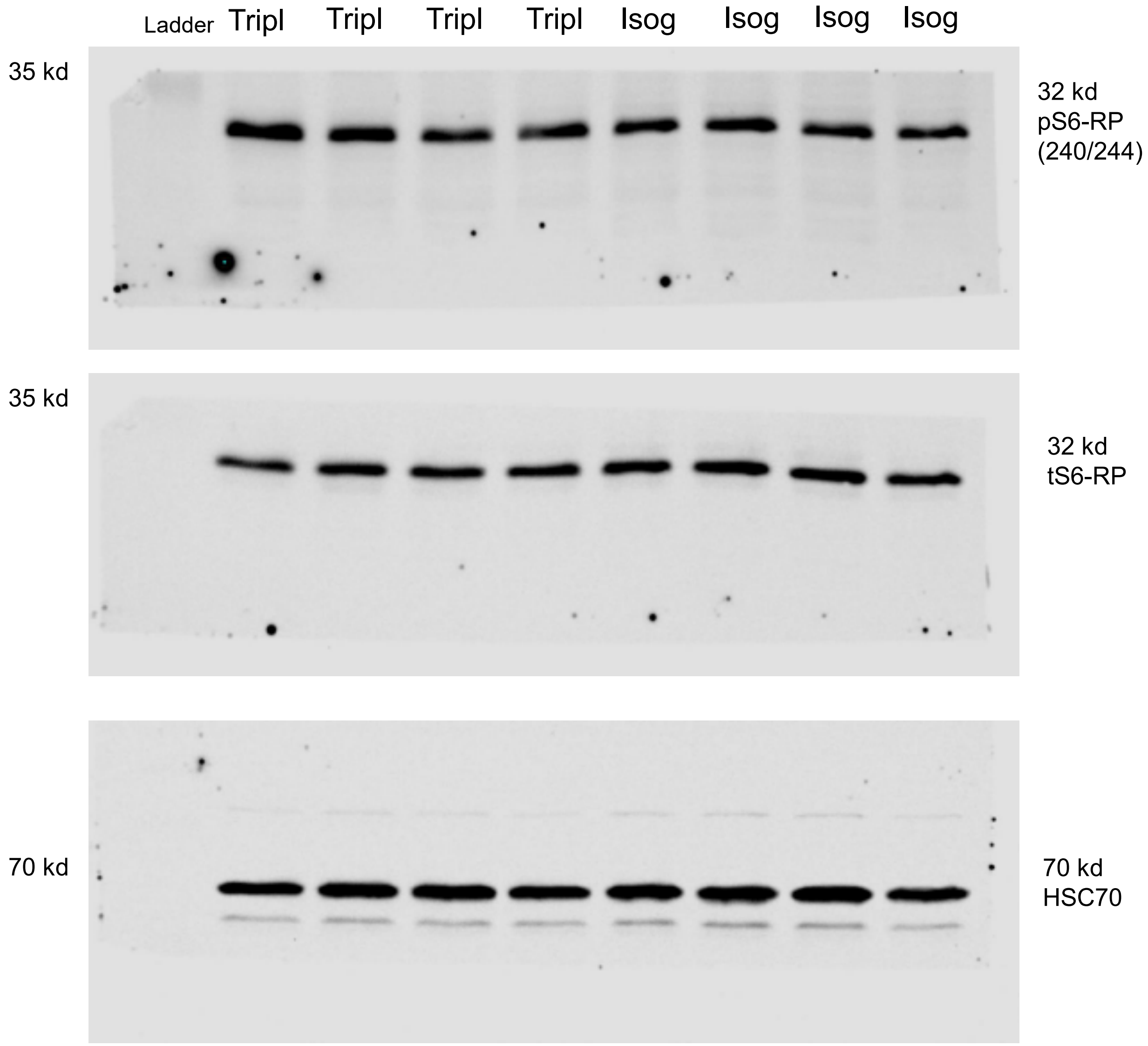

Ladder
Tripl
Tripl
Tripl
Tripl
Isog
Isog
Isog
Isog
35 kd
32 kd
pS6-RP
(240/244)
35 kd
32 kd
tS6-RP
70 kd
70 kd
HSC70

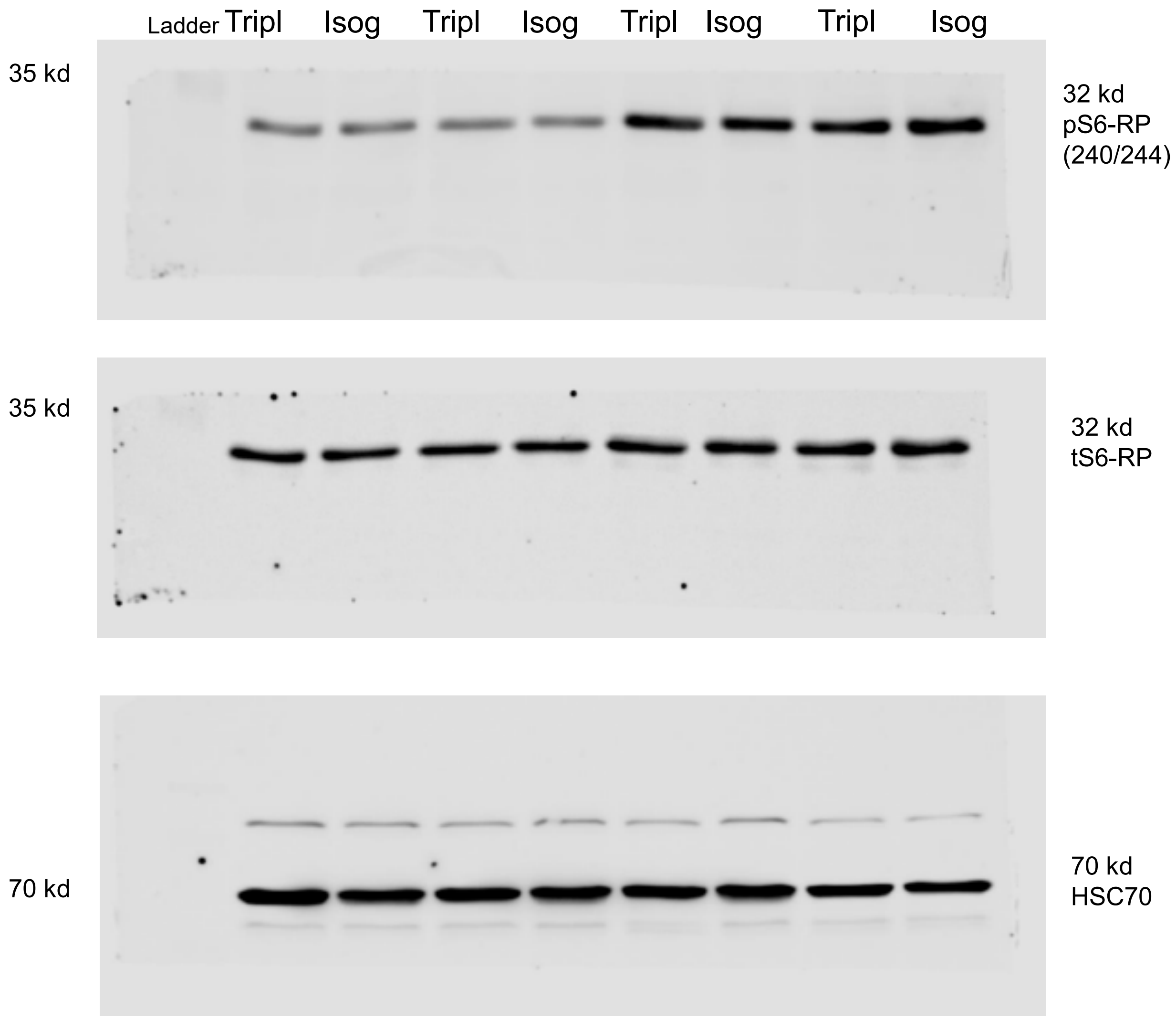
Ladder
Tripl
Isog
Tripl
Isog
Tripl
Isog
Tripl
Isog
35 kd
32 kd
pS6-RP
(240/244)
35 kd
32 kd
tS6-RP
70 kd
70 kd
HSC70

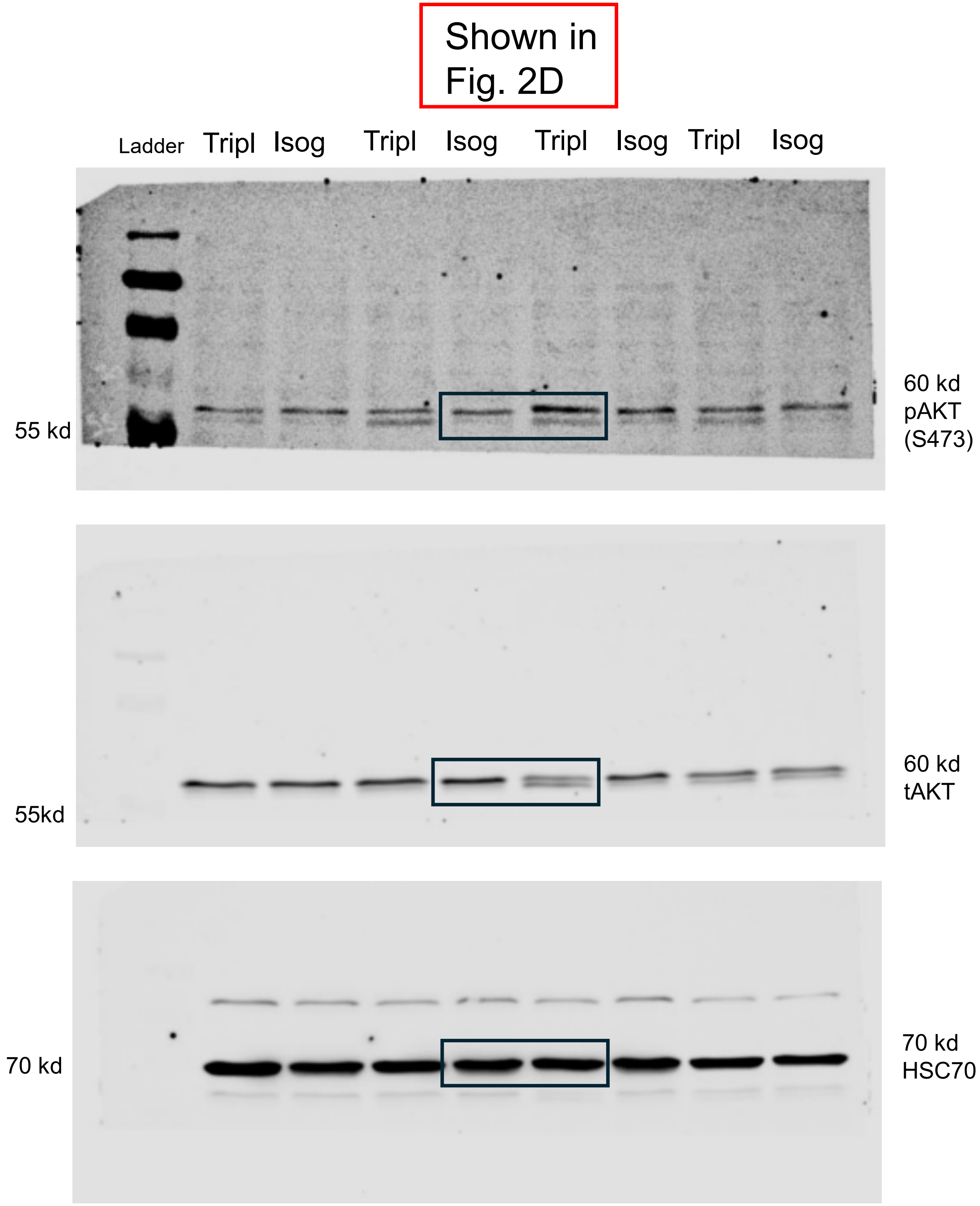
Shown in
Fig. 2D
Ladder
Tripl
Isog
Tripl
Isog
Tripl
Isog
Tripl
Isog
55 kd
60 kd
pAKT
(S473)
55kd
60 kd
tAKT
70 kd
70 kd
HSC70

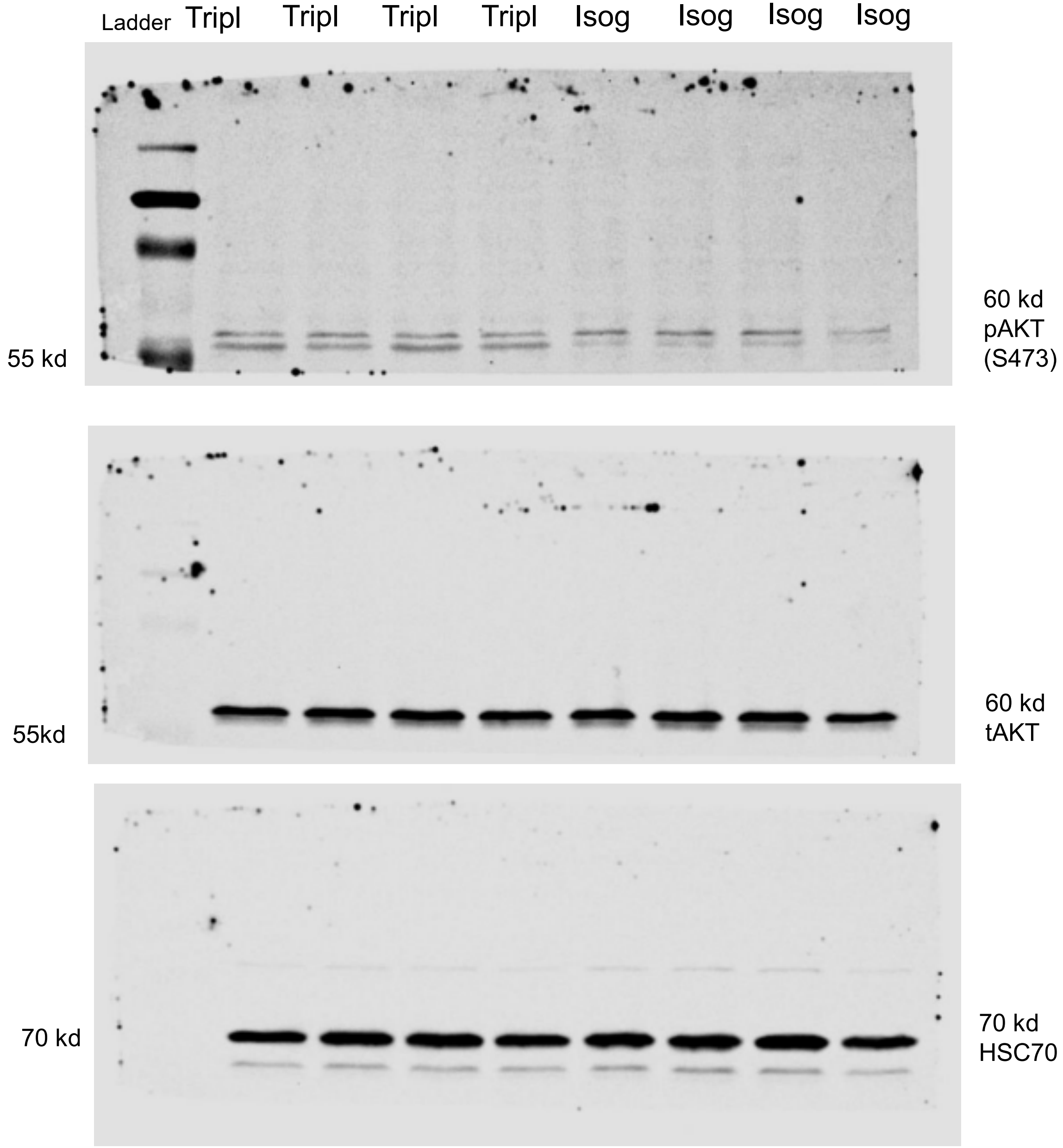
Ladder
Tripl
Tripl
Tripl
Tripl
Isog
Isog
Isog
Isog
55 kd
60 kd
pAKT
(S473)
55kd
60 kd
tAKT
70 kd
70 kd
HSC70

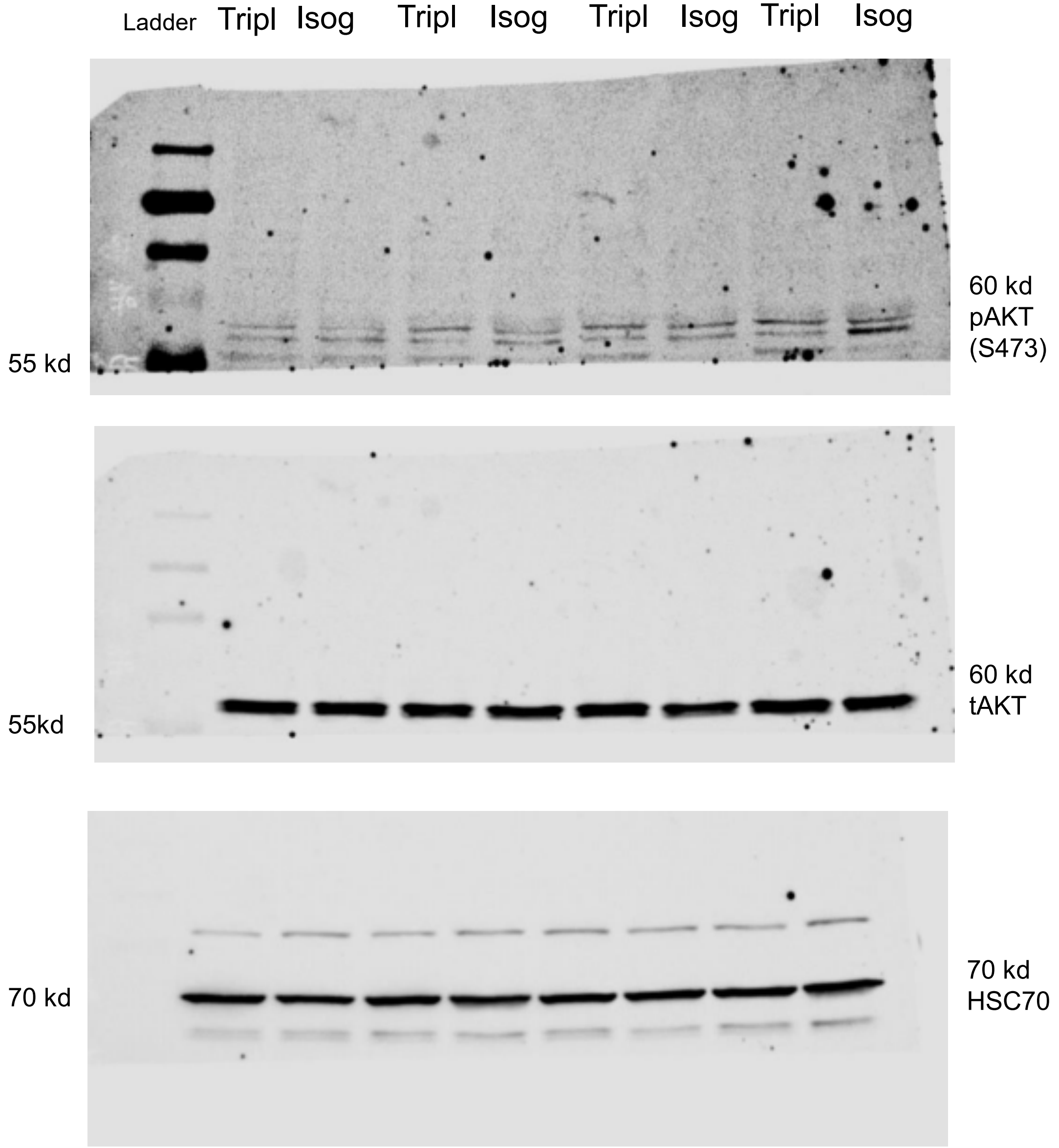
Ladder
Tripl
Isog
Tripl
Isog
Tripl
Isog
Tripl
Isog
55 kd
60 kd
pAKT
(S473)
55kd
60 kd
tAKT
70 kd
70 kd
HSC70

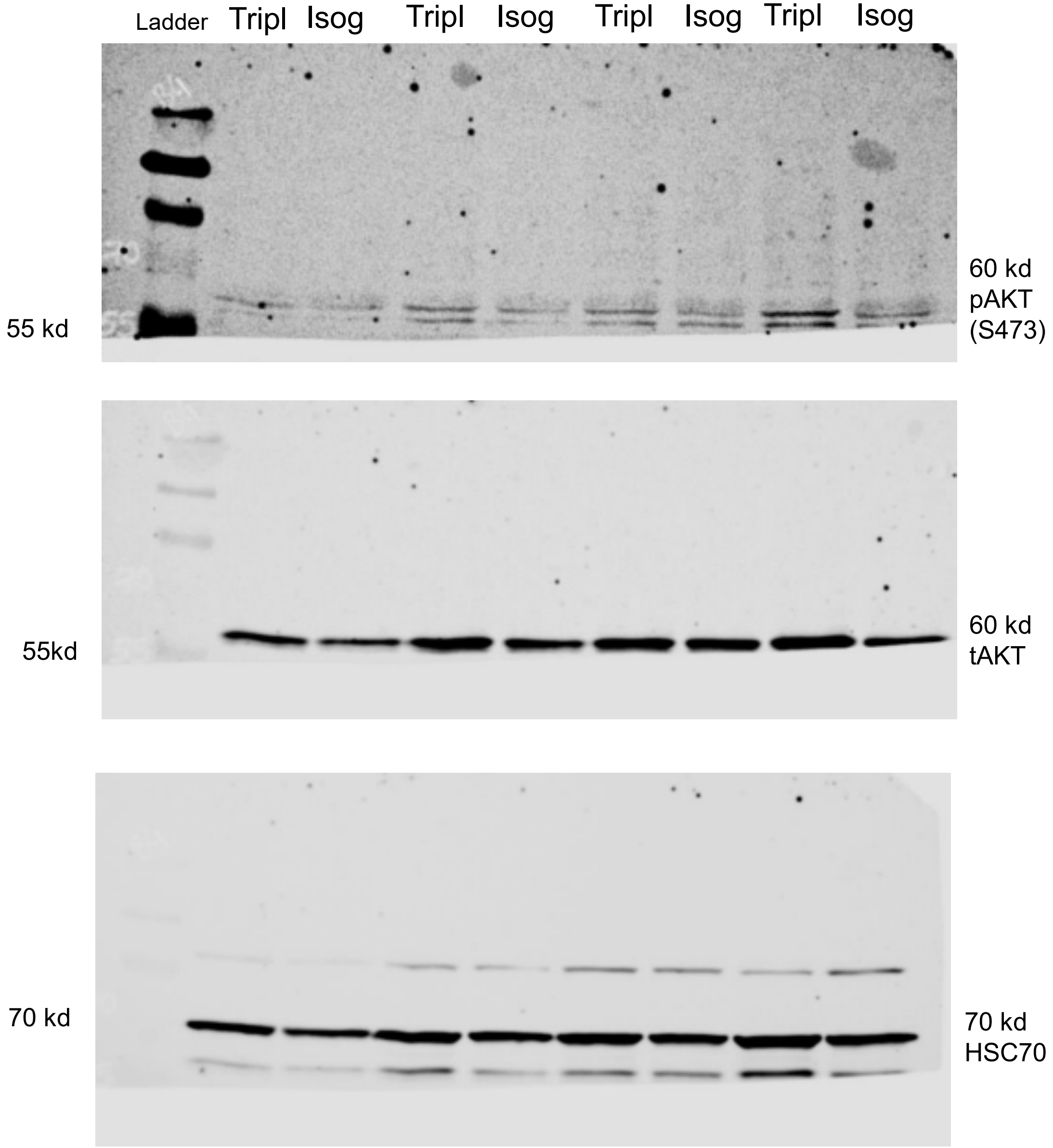
Ladder
Tripl
Isog
Tripl
Isog
Tripl
Isog
Tripl
Isog
55 kd
60 kd
pAKT
(S473)
55kd
60 kd
tAKT
70 kd
70 kd
HSC70

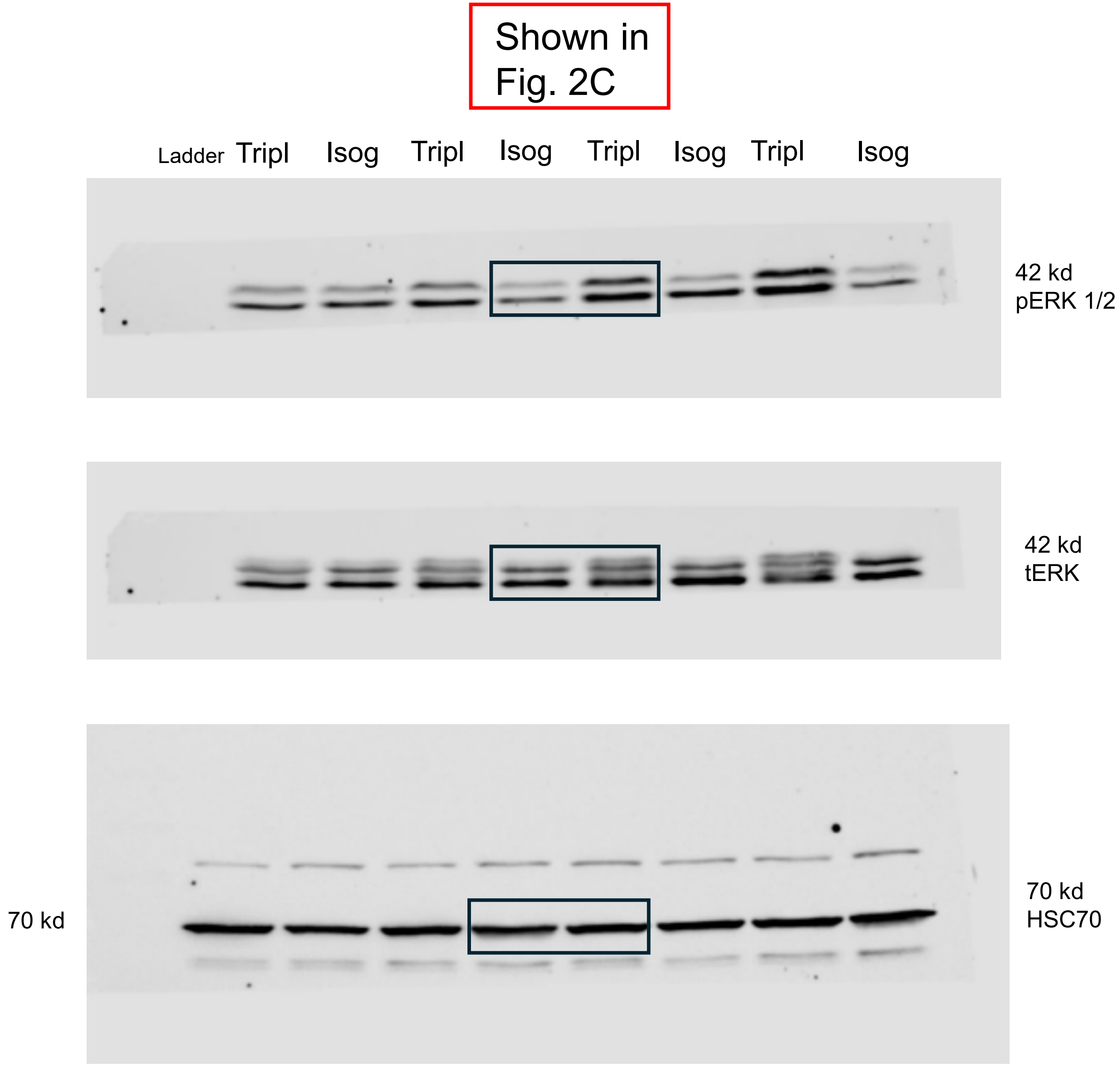

Shown in Fig. 2C
Ladder
Tripl
Isog
Tripl
Isog
Tripl
Isog
Tripl
Isog
42 kd
pERK 1/2
42 kd
tERK
70 kd
70 kd
HSC70

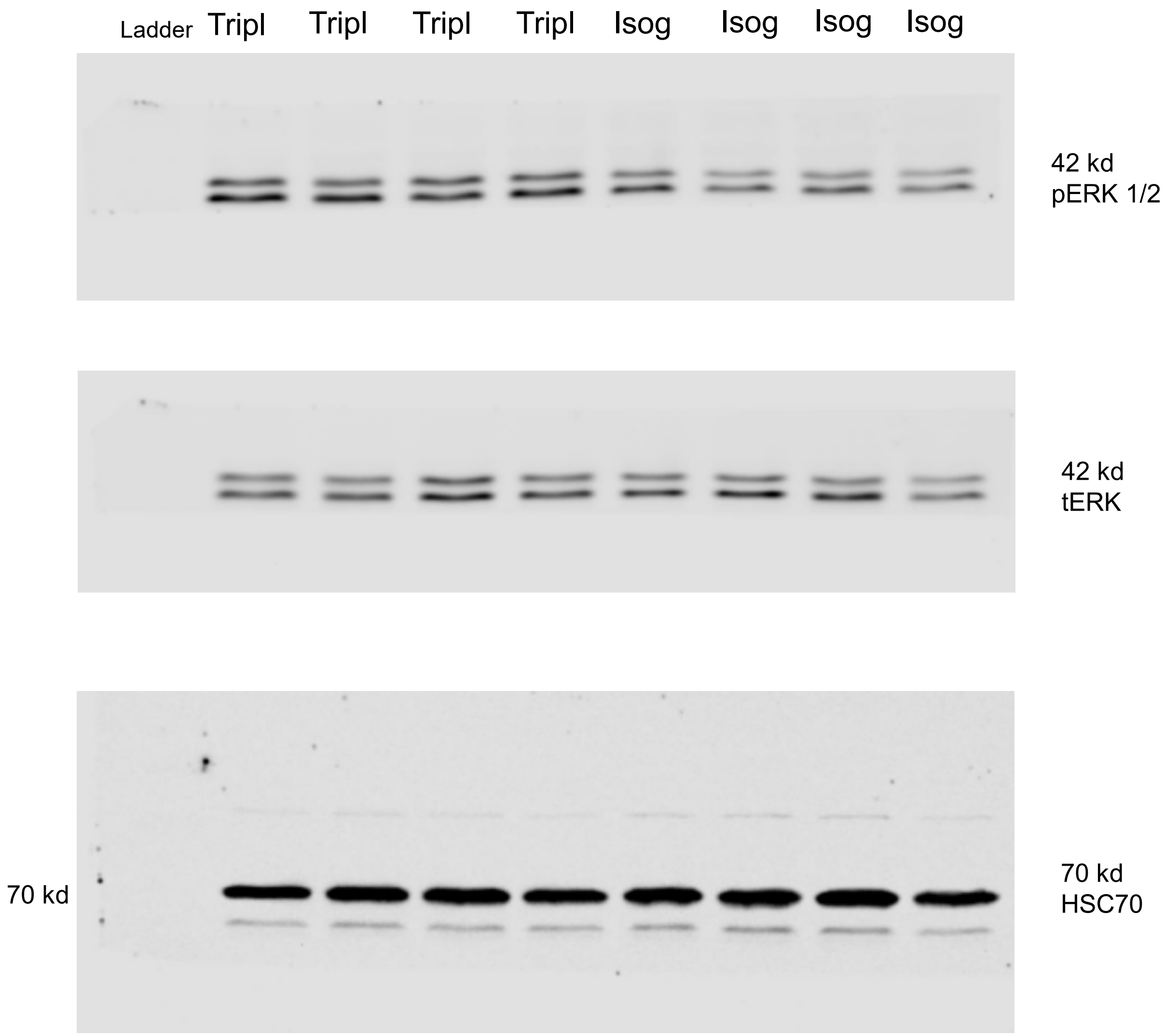
Ladder
Tripl
Tripl
Tripl
Tripl
Isog
Isog
Isog
Isog
42 kd
pERK 1/2
42 kd
tERK
70 kd
70 kd
HSC70

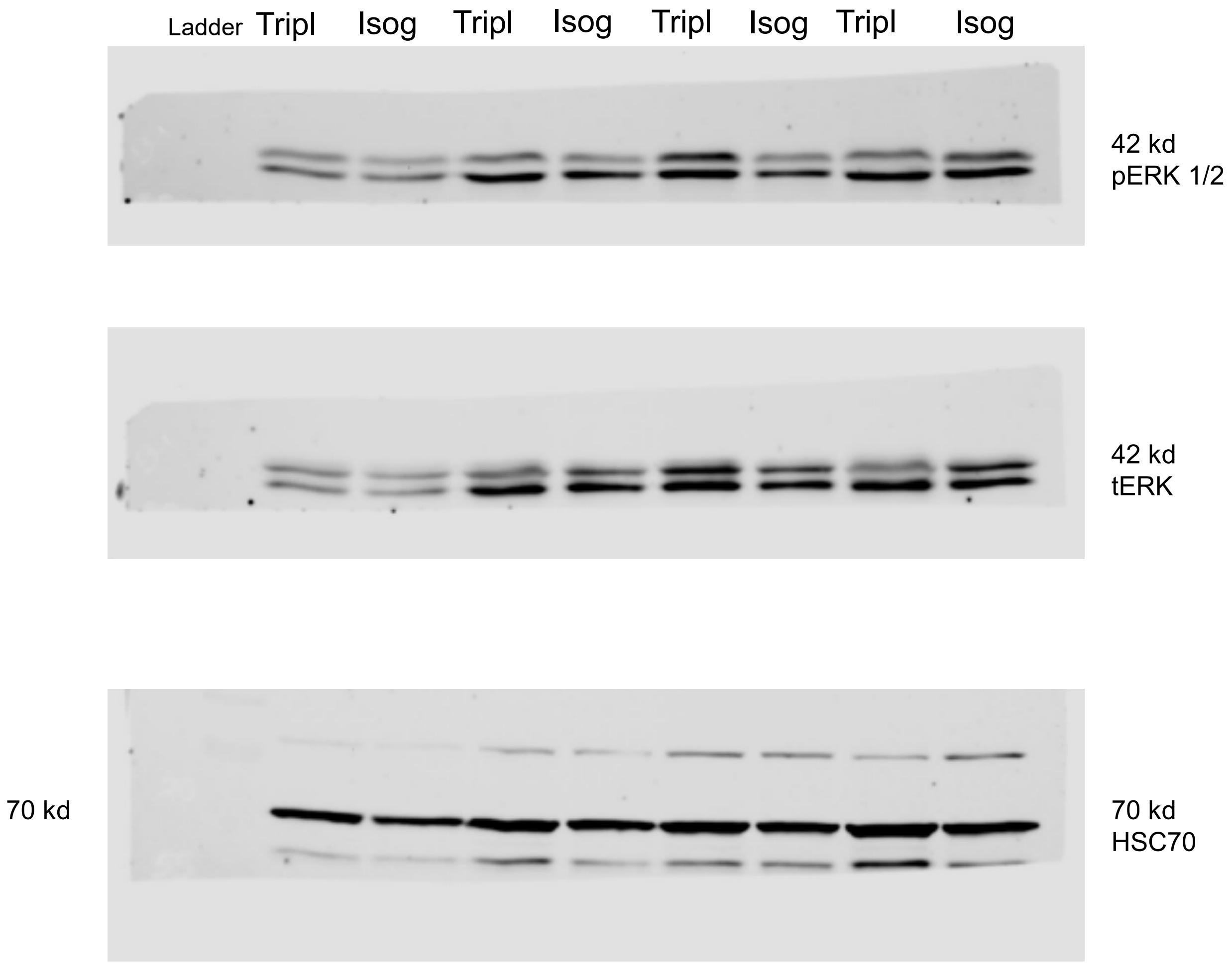

Ladder
Tripl
Isog
Tripl
Isog
Tripl
Isog
Tripl
Isog
42 kd
pERK 1/2
42 kd
tERK
70 kd
70 kd
HSC70

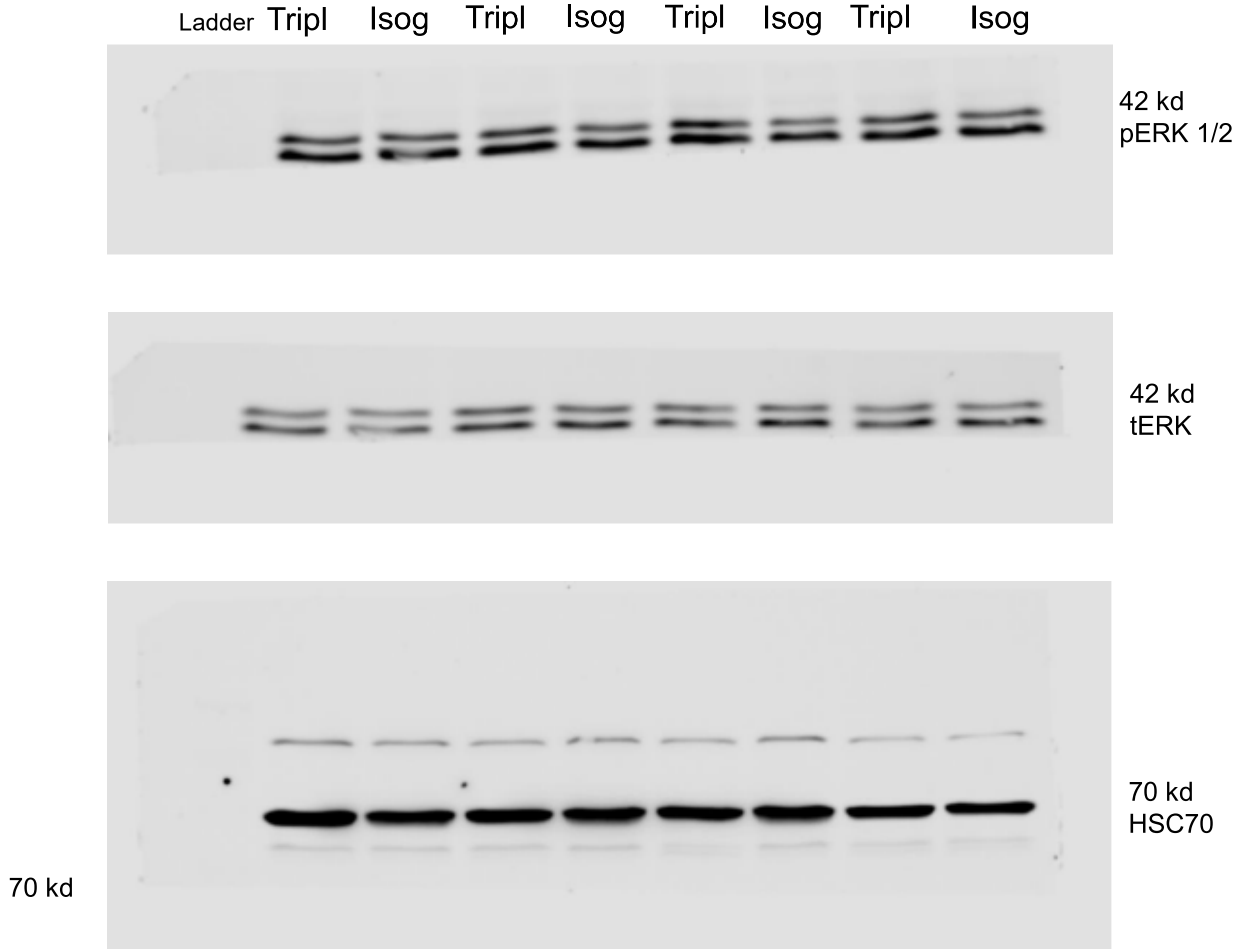
Ladder
Tripl
Isog
Tripl
Isog
Tripl
Isog
Tripl
Isog
42 kd
pERK 1/2
42 kd
tERK
70 kd
HSC70
70 kd

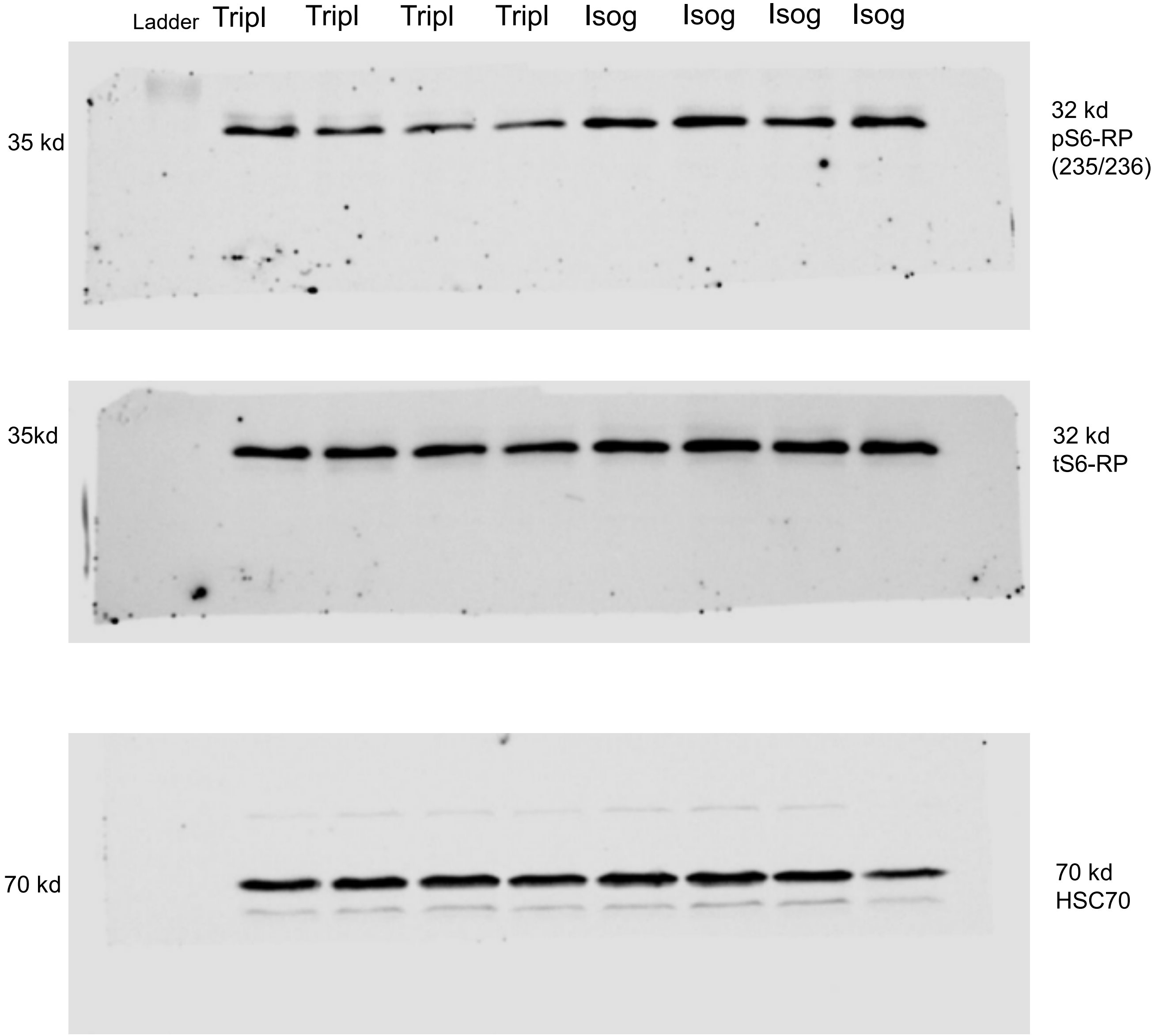

Ladder
Tripl
Tripl
Tripl
Tripl
Isog
Isog
Isog
Isog
35 kd
32 kd
pS6-RP
(235/236)
35kd
32 kd
tS6-RP
70 kd
70 kd
HSC70

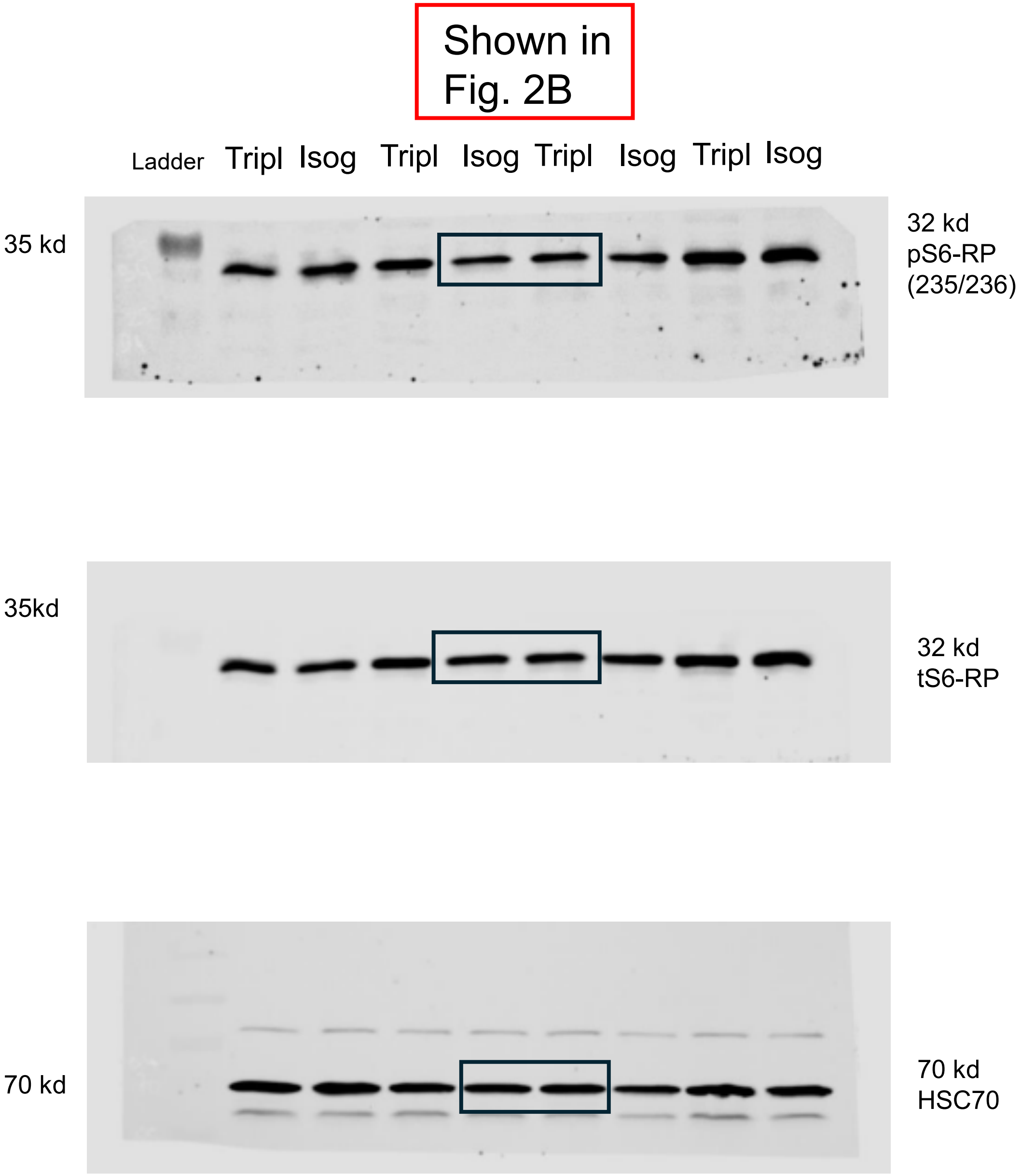
Shown in Fig. 2B
Ladder
Tripl
Isog
Tripl
Isog
Tripl
Isog
Tripl
Isog
35 kd
32 kd
pS6-RP
(235/236)
35kd
32 kd
tS6-RP
70 kd
70 kd
HSC70

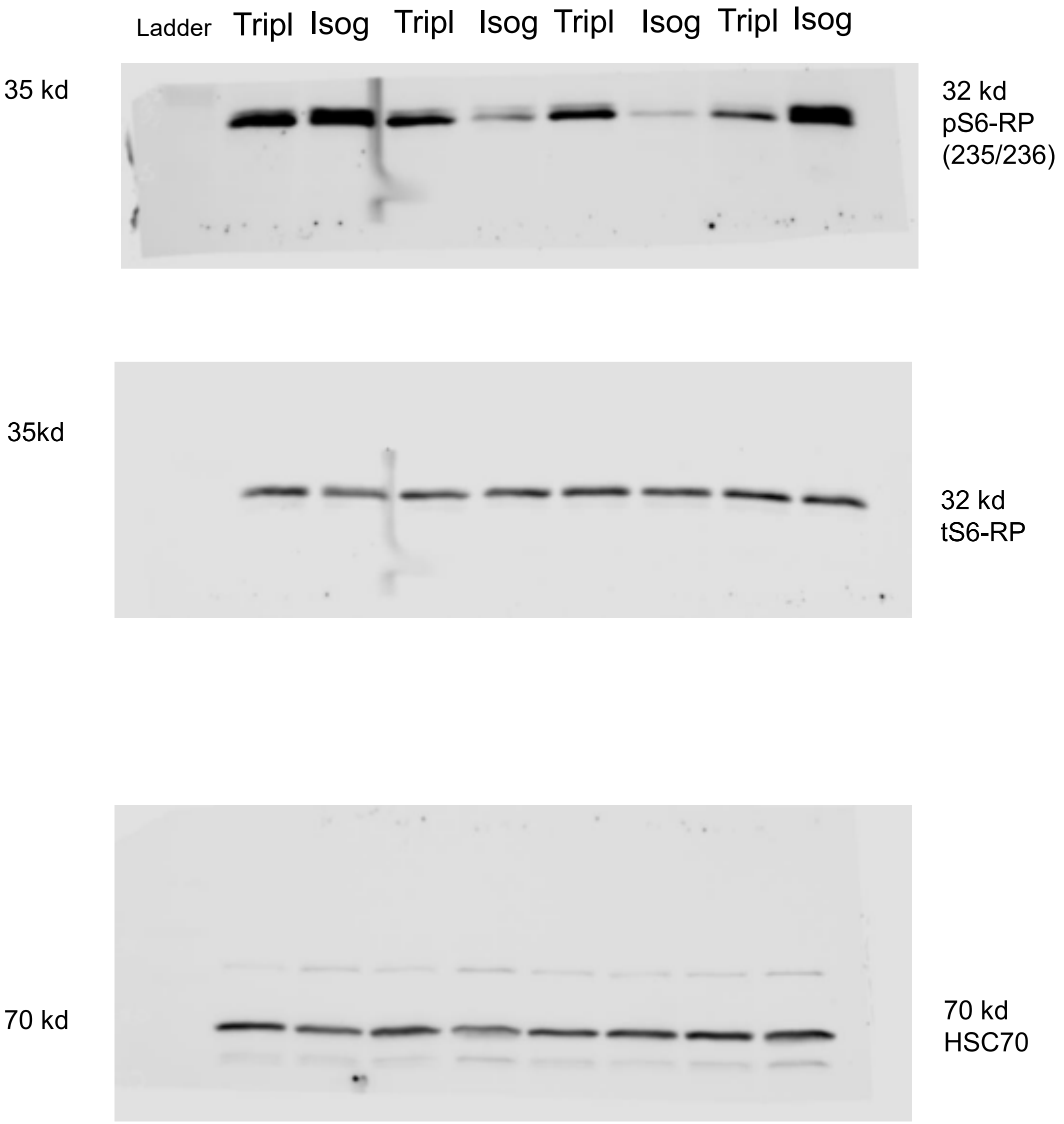
Ladder
Tripl
Isog
Tripl
Isog
Tripl
Isog
Tripl
Isog
35 kd
32 kd
pS6-RP
(235/236)
35kd
32 kd
tS6-RP
70 kd
70 kd
HSC70

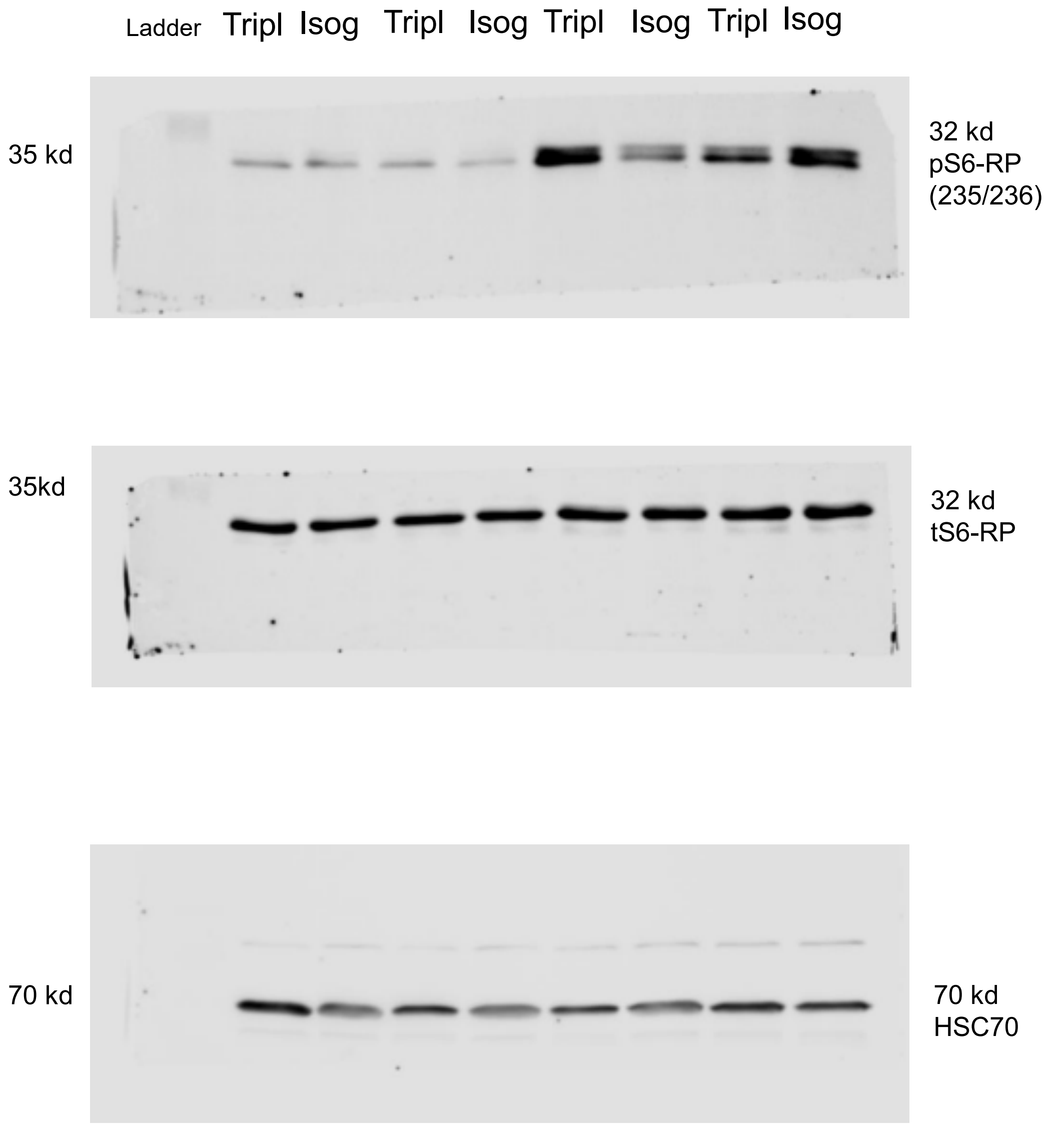
Ladder
Tripl
Isog
Tripl
Isog
Tripl
Isog
Tripl
Isog
35 kd
32 kd
pS6-RP
(235/236)
35kd
32 kd
tS6-RP
70 kd
70 kd
HSC70

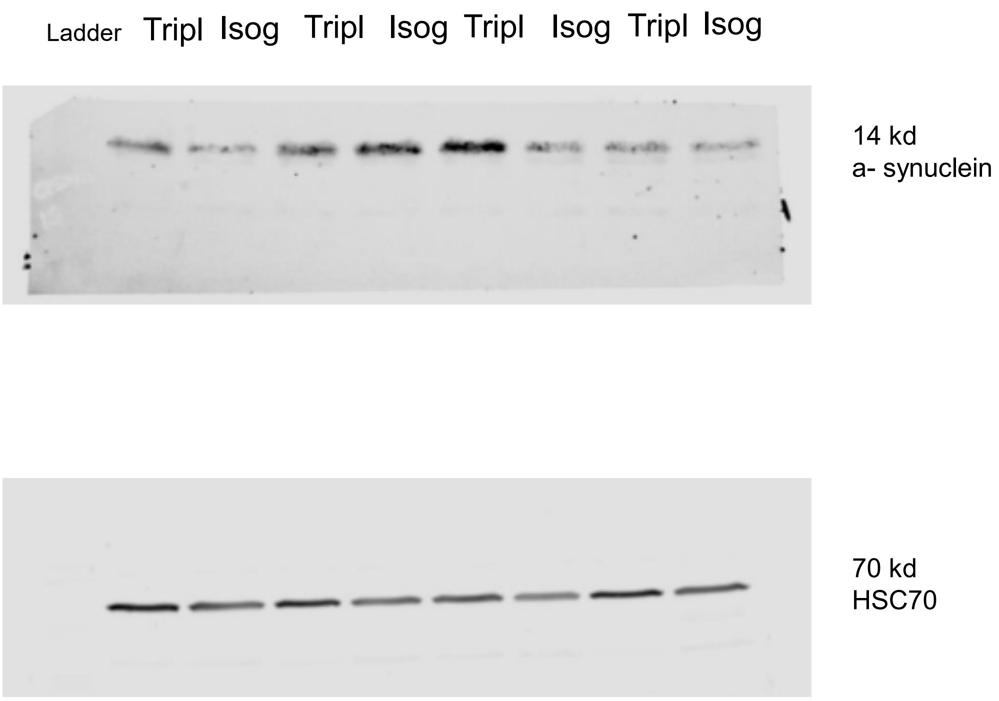
Ladder
Tripl
Isog
Tripl
Isog
Tripl
Isog
Tripl
Isog
14 kd
a- synuclein
70 kd
HSC70

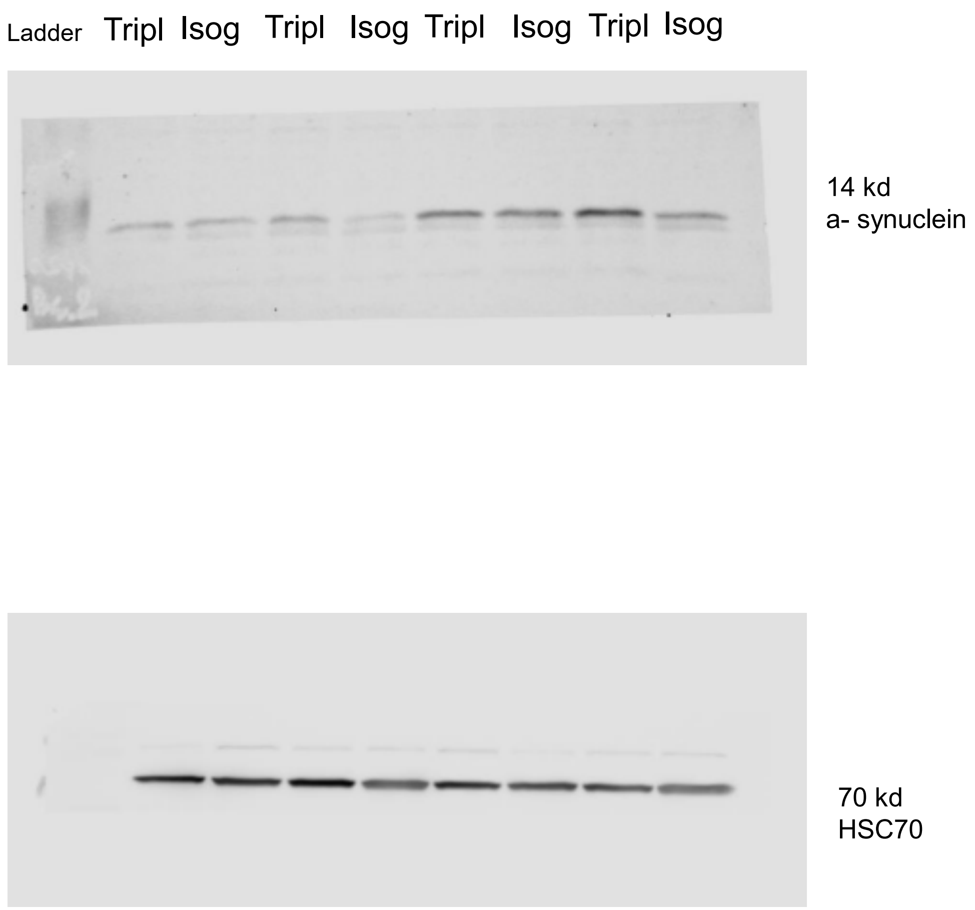
Ladder
Tripl
Isog
Tripl
Isog
Tripl
Isog
Tripl
Isog
14 kd
a- synuclein
70 kd
HSC70

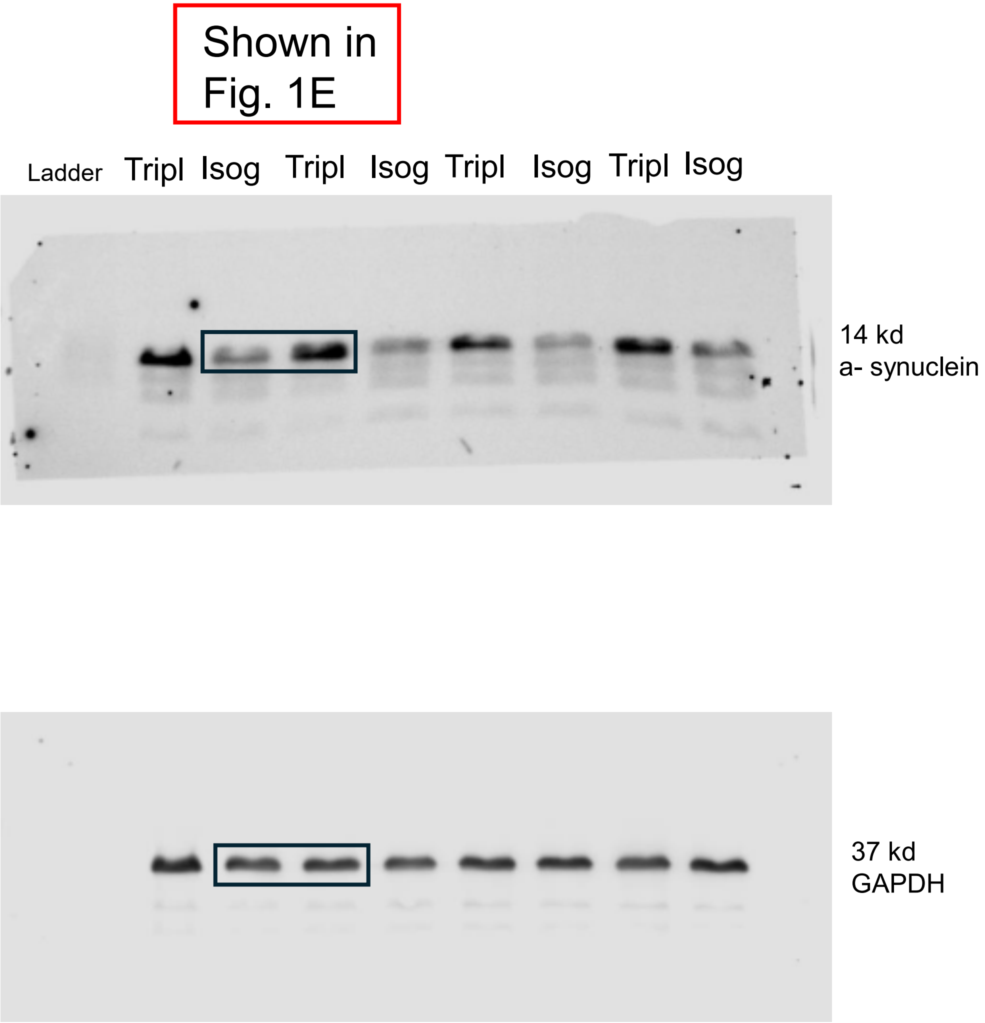
Shown in Fig. 1E
Ladder
Tripl
Isog
Tripl
Isog
Tripl
Isog
Tripl
Isog
14 kd
a- synuclein
37 kd
GAPDH

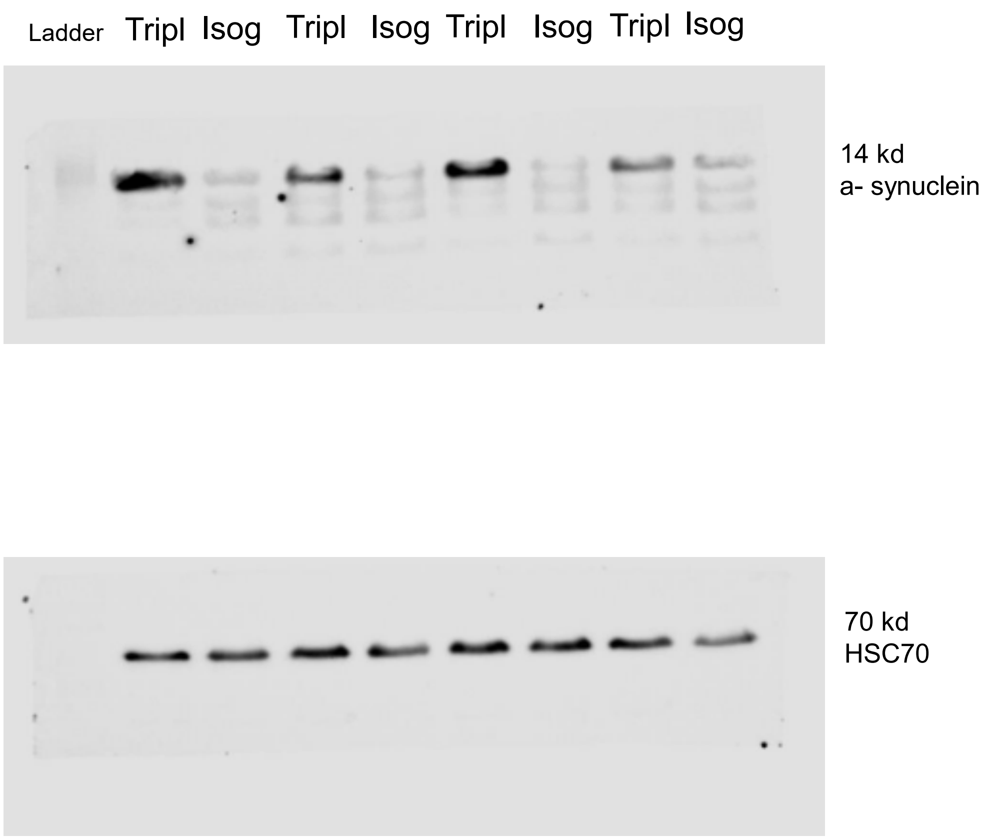
Ladder Tripl Isog Tripl Isog Tripl Isog Tripl Isog
14 kd
a- synuclein
70 kd
HSC70

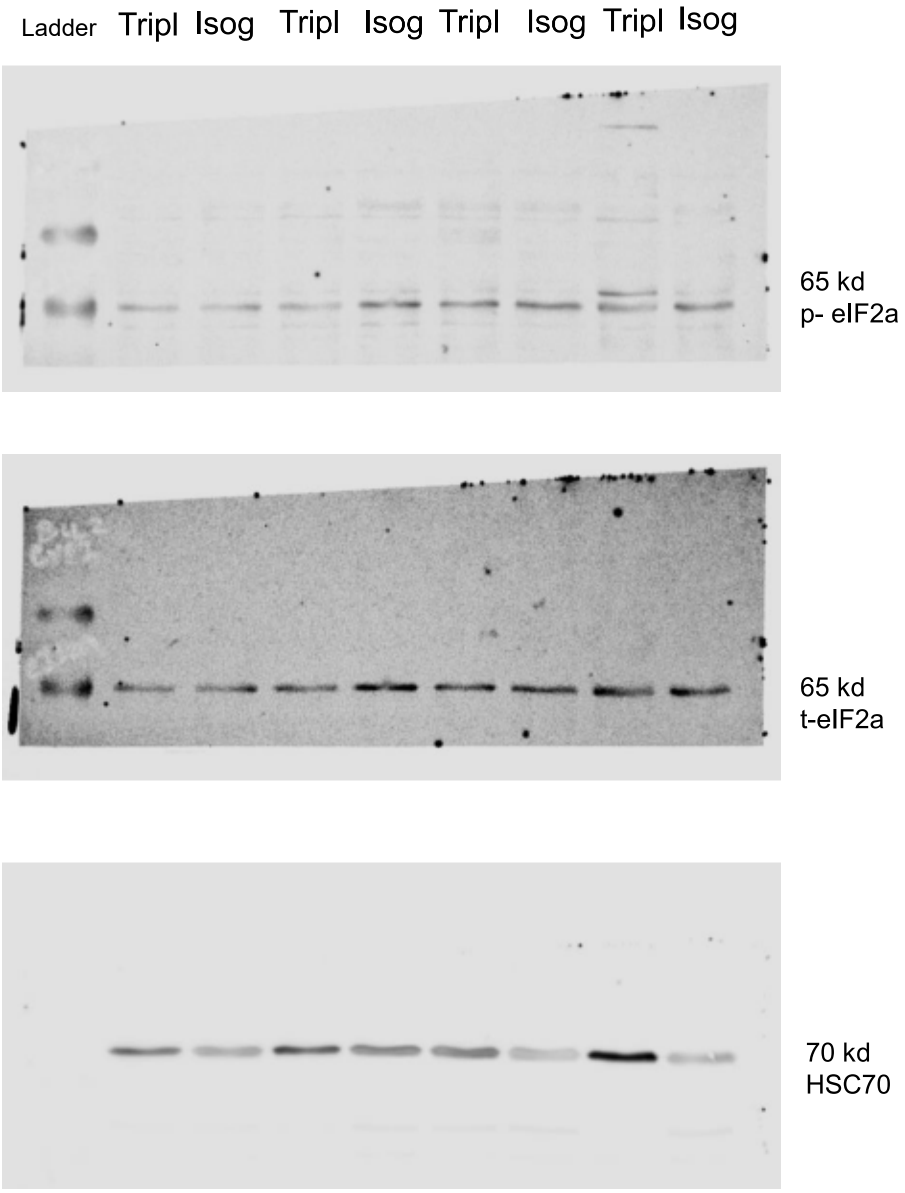

Ladder
Tripl
Isog
Tripl
Isog
Tripl
Isog
Tripl
Isog
65 kd
p- eIF2a
65 kd
t-eIF2a
70 kd
HSC70

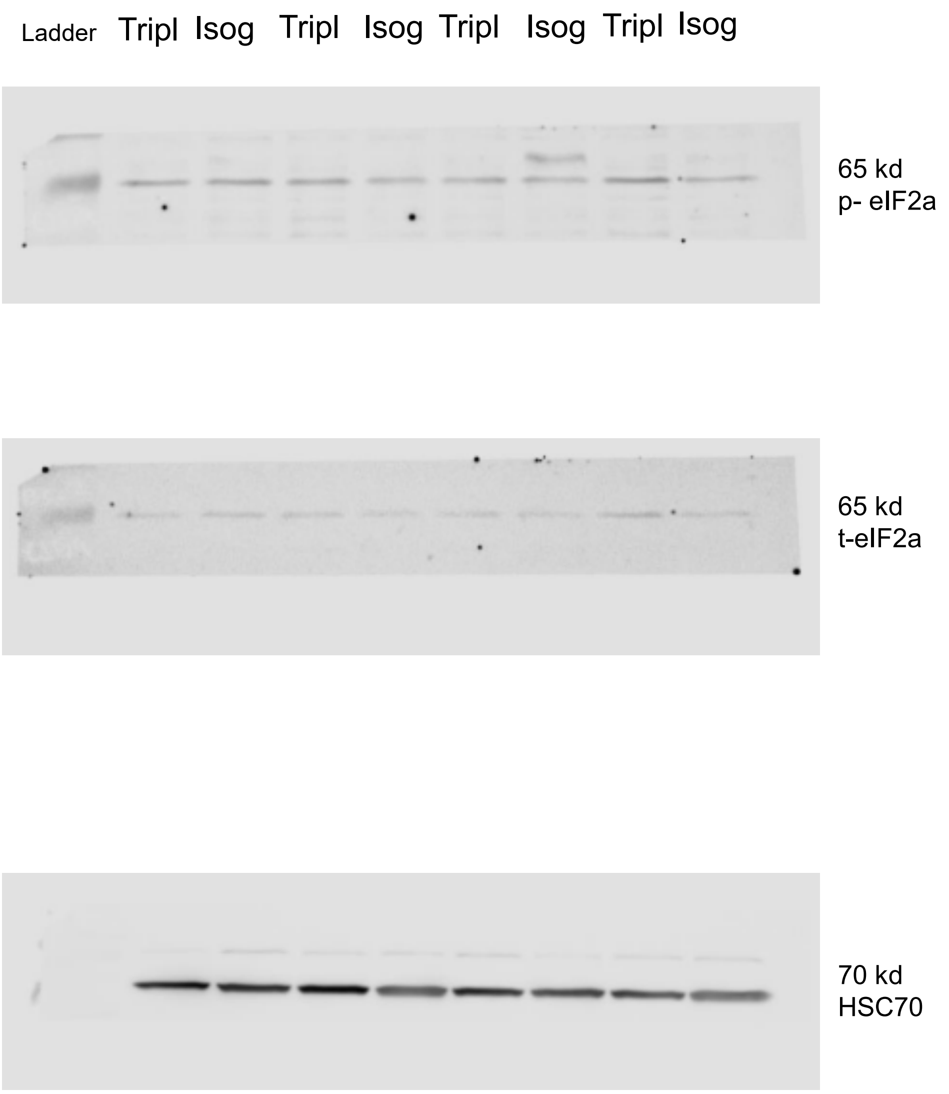
Ladder
Tripl
Isog
Tripl
Isog
Tripl
Isog
Tripl
Isog
65 kd
p- eIF2a
65 kd
t-eIF2a
70 kd
HSC70

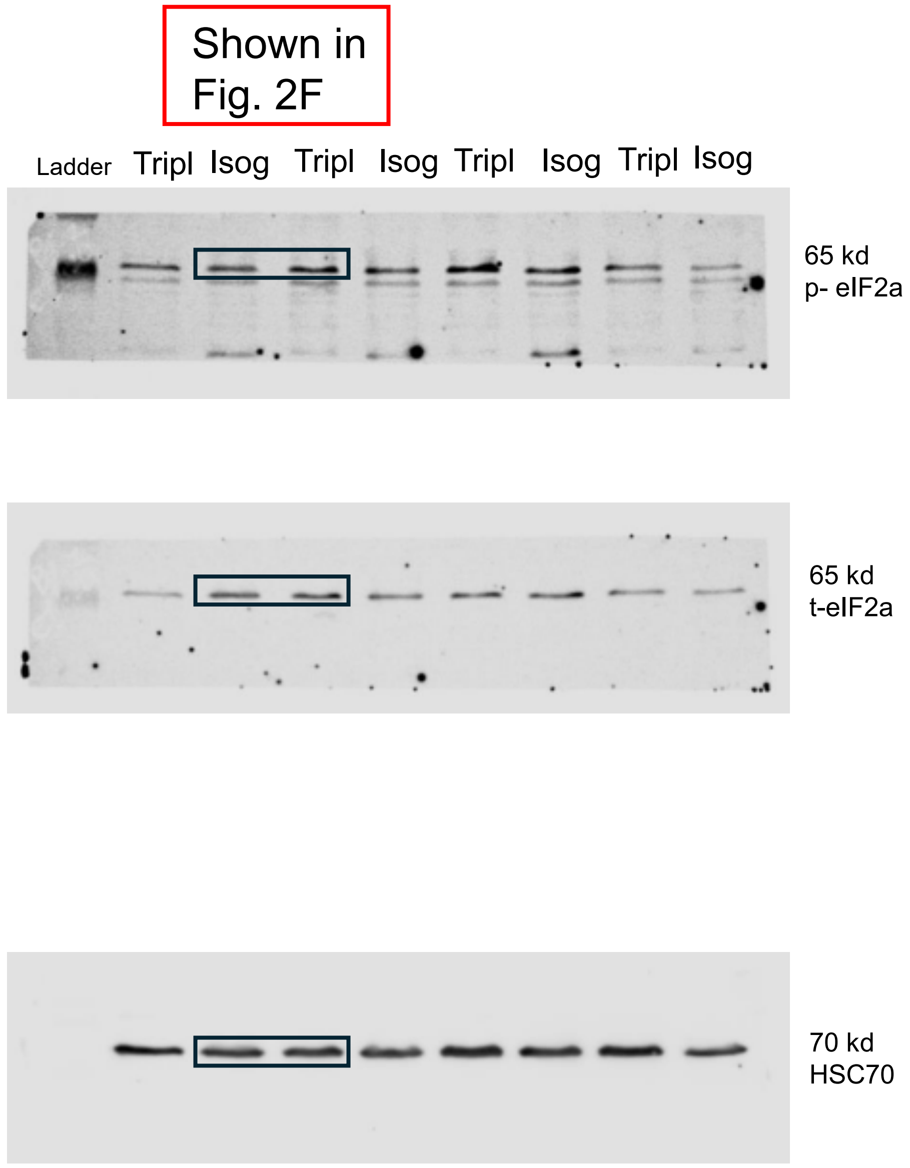
Shown in Fig. 2F
Ladder
Tripl
Isog
Tripl
Isog
Tripl
Isog
Tripl
Isog
65 kd
p- eIF2a
65 kd
t-eIF2a
70 kd
HSC70
